# A Complete Telomere-to-Telomere Diploid Reference Genome for South Asian Population

**DOI:** 10.1101/2025.07.12.664550

**Authors:** Prasad Sarashetti, Josipa Lipovac, Qi Jia, Ling Wang, Zhe Li, Lovro Vrcek, Ying Chen, Fei Yao, YY Sia, Dehui Lin, Xinyi Zhang, Daniel Muliaditan, Hwee Meng Low, See Ting Leong, Chiea Chuen Khor, Eleanor Wong, Weiling Zheng, Kresimir Krizanovic, John C Chambers, William Y K Hwang, Patrick Tan, Jonathan Göke, Mile Sikic, Jianjun Liu

## Abstract

Human reference genomes have been instrumental in advancing genomic and biomedical research, but South and Southeast Asian populations are underrepresented, despite accounting for a large proportion of world population. As a part of effort on generating reference genomes for these populations, we present the first gapless, telomere-to-telomere (T2T) diploid genome assembly created by using a trio sample set of Indian ancestry (I002C), with NG50 of 154.89 Mb and 146.27 Mb for the maternal and paternal haplotypes, including the fully assembled rDNA array for the maternal chromosome 21 and Y chromosome. With the Merqury QVs of 82.05, 83.08 and 82.64 for the maternal, paternal and haploid assemblies respectively, I002C represents the highest-quality human genome assembled in both diploid and haploid forms to date. Compared to CHM13, the I002C genome displays substantial sequence diversity, resulting in 14,943 structural variants, including 3,236 novel variants absent from public databases. Analysis of trio-phased haplotypes further revealed elevated inter-haplotype divergence within centromeric and subtelomeric regions, along with identification of differentially methylated regions (DMRs) as candidates for novel imprinting loci. As a result of substantial SVs between them, I002C is a more suitable reference than CHM13 for the genomic analysis of South Asian samples with less reference bias and better performance in mapping and variant calling, particularly for long read sequencing data. As the first high-quality T2T diploid reference genome for Indian, the largest world’s population, I002C contributes to the growing set of population-specific reference genomes and helps to overcome a significant gap in human genome diversity.

## Introduction

A human reference genome serves as the foundation for genomic analyses, such as gene and regulatory sequence annotations, transcriptome quantification, and variation analysis, making it an essential resource for biomedical and clinical research to understand human genetic diversity, disease mechanisms, and evolutionary biology, and enable precision medicine. The most widely used reference genome, GRCh38, was assembled from a few donor individuals, primarily of African and European descent^1^. Despite its widespread use, GRCh38 still contains unresolved regions, including ribosomal DNA (rDNA) arrays, as well as pericentromeric and subtelomeric regions^2^, which are critical for understanding fundamental cellular processes. Trying to overcome these limitations, there have been several efforts on generating new human reference genomes with better completeness and quality, such as CHM13^2^, HG002^3^, CN1^4^ and YAO^5^ by taking the advantages of long read sequencing technologies and new algorithms for de novo assembling analysis. More recently, recognizing the limitation of current references in capturing the genetic diversity information, efforts have gone beyond the generation of complete, telomere-to-telomere (T2T) and phased reference genomes to build pangenome reference consisting of a panel of multiple reference genomes, such as the project led by the Human Pangenome Reference consortium (HPRC)^6,7^.

These efforts are, however, mainly focused on analyzing individuals of European and East Asian ancestries, with over 85% of studies involving populations of European descent^8,9^. Despite representing around 34% of the world population, South and Southeast Asian populations are very much under-represented in these efforts, with only very few studies being carried out^10–18^, and currently no complete T2T reference genome is available. It has been increasingly recognized that alignment biases, errors in variant calling, misinterpretation of pathogenic mutations, and incorrect inferences may arise when the reference genome for analysis is very different from the samples under study^4,7,19^. These limitations of reference bias hamper genetic studies and translation to clinical applications in underrepresented populations and call for the generation of ancestry-specific T2T reference genomes.

Here we present the first high-quality, gapless, telomere-to-telomere, haplotype-resolved reference genome (I002C) with 44 autosomes and XY chromosomes for South Asia, using a biologically normal Indian sample, from a father-mother-child trio samples set recruited in Singapore. We employed state-of-the-art sequencing technologies, including PacBio HiFi circular consensus long-read sequencing and Oxford Nanopore ultralong sequencing, and advanced algorithms for de novo assembling, including Verkko^20^ and Hifiasm^21,22^ to construct chromosome-level contigs, including the long and highly repetitive centromeric regions and the short arms of five acrocentric chromosomes. We utilized both parental sequencing data information for phasing and generating the paternal and maternal haploid genomes, and we used long read RNA-Seq data combined with reference genome annotations to provide a comprehensive set of transcripts for the I002C genome. We further demonstrated that with this reference genome, South Asian (SAS)-specific novel variants can be identified with greater accuracy, and sequencing results from SAS samples can be analyzed with higher mapping rates and lower clipping rates. This effort has helped to overcome a significant gap in the presentation of human genome diversity by providing the first reference genome for the SAS population, the largest one in the world by accounting for 25% of the world population.

## Results

### Sample and Sequencing Data

To assemble a diploid, telomere-to-telomere (T2T) reference genome, a healthy Singaporean male and corresponding parental samples with self-reported Indian ancestry for all the grandparents were recruited. Karyotyping at a resolution of 450 bands confirmed the diploid nature of the child sample (I002C) (Additional file 1: Figure S1). Using the DNAs of the child sample cell line, we generated 318.62 Gb (103x) of Pacbio HiFi, 193.66 Gb (62x) of Nanopore duplex, and 669.94 Gb (216x) Nanopore ultra-long reads, complemented with 70.53 GB of Nanopore cDNA data. In addition, the short-reads dataset included 297.61 Gb (96x) from MGI, 371.18 Gb (119x) OmniC and 22 Gb total RNASeq from the Illumina platform (Additional file 2: Table S1). PacBio HiFi and Nanopore duplex reads were also generated for the two parental samples (Additional file 2: Table S1). The reference genome generated from this child sample was intended to represent one of the most underrepresented populations in global genomics—SAS (Figure 1). Local ancestry analysis based on data from the 1000 Genomes Project^23^ revealed that over 90% of the I002C genome is of SAS origin, while the second-largest contribution is from European populations, along with scattered markers from East Asian and American, consistent with previous findings^24^ (Additional file 1: Figure S2). Further comparison of ancestry markers between the two haplotypes shows diversity, highlighting inter-haplotype variation. For example, on chromosome 15, the region distal to the centromere is inferred as American in origin on the maternal haplotype, whereas the corresponding paternal region is of European ancestry (Figure 1). We also performed the PCA analysis of I002C and the 5 SAS subpopulation samples from the 1000 Genomes Project. The analysis showed that I002C clustered with the SAS group overall, with a closer affinity to Gujarati Indians from Houston (GIH), Punjabis from Lahore (PJL), and Indian Telugu from the UK (ITU) (Additional file 1: Figure S3). This clustering pattern further supports the South Asian origin of I002C.

**Figure 1:**
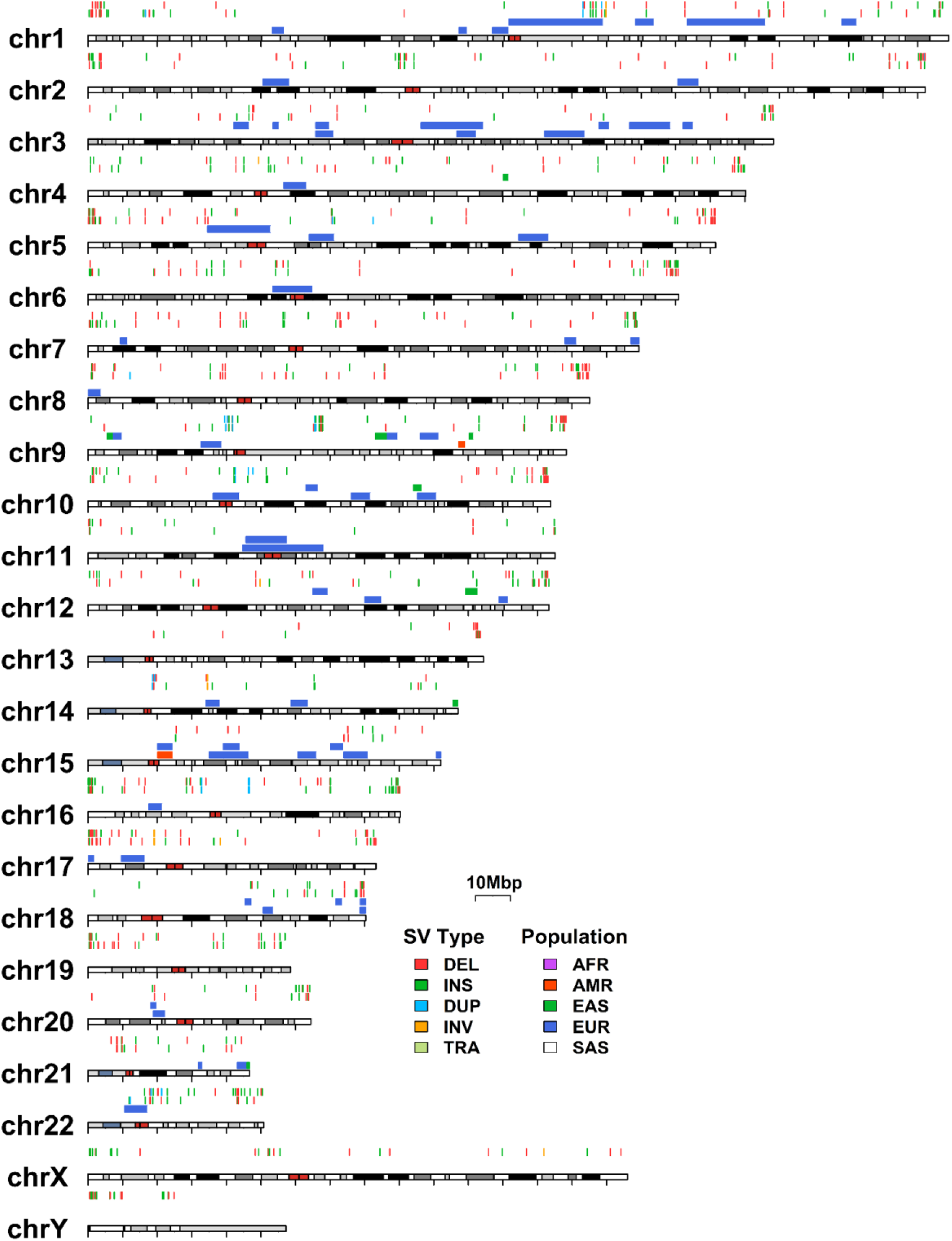
Ancestral composition and unique structural variants of I002C. Overview of ancestry markers and structural variants across I002C haplotypes. Chromosomal regions are coloured by different tracks based on predicted ancestral marker and structural variant types. For each chromosome (4 tracks from bottom to top), (1) ancestral markers for the maternal haplotype; (2) ancestral markers for the paternal haplotype; (3) structural variants unique to the maternal haplotype and (4) structural variants unique to the paternal haplotype, identified relative to CHM13 and absent from gnomAD, HGSVC, and HPRC. **Populations:** AFR, African; AMR, American; EAS, East Asian; EUR, European; SAS, South Asian. **Structural variant types:** DEL, deletion; INS, insertion; DUP, duplication; INV, inversion; TRA, translocation.

### A telomere-to-telomere diploid assembly, polishing, and assessment

Initial assemblies were generated using Verkko^20^ and Hifiasm^21,22^ in trio mode. For each haplotype, we selected chromosome-scale sequences from either assembler based on quality, contiguity and presence of telomere motif on both ends. This resulted in 20 maternal chromosomes (excluding the p-arms of chromosomes 13, 15 and 22) with 39 gaps, and 19 paternal chromosomes (excluding the p-arms of chromosomes 4, 13, 14 and 15) with 14 gaps (Additional methods: **Contig construction)**. The remaining chromosomes were generated using template-based scaffolding (Additional Methods: **Scaffolding**). The gaps were closed using combinatorial methods, including TGS-Gapcloser^25^, contigs from local reassemblies of binned reads and ultra-long read alignments (Additional methods: **Gap-filling**). The remaining telomeres were patched using subtelomere ultra-long read mapping and local assemblies (Additional methods: **Telomere patching and extension**) (Additional file 2: Table S2). The rDNA arrays were verified and corrected by local re-assembly and manual curation (Additional methods: **rDNA**). The final assembly includes full T2T sequences for all 22 autosome pairs, both sex chromosomes (X and Y), and the mitochondrial genome.

To further improve assembly quality, we performed multiple rounds of polishing using binned HiFi, ONT reads (duplex, corrected and uncorrected ultra-long), and MGI short reads (Additional methods: **Read binning**, Additional file 2: Table S3-S4). The polishing workflow included two rounds of automated polishing^26^, one round of manual curation (Additional file 1: Figure S4), and one round of polishing using assemblies from corrected ultra-long reads. This process increased the assembly quality (QV) from an initial 68.05 to 82.65 (average of two haplotypes)(Figure 2a), as assessed by Merqury^27^ using hybrid k-mers (k=21) obtained from MGI short reads and HiFi reads. For more than two-thirds of the chromosomes in both maternal and paternal haplotypes, the Merqury QV exceeded the genome-wide average of 83, with 25 chromosomes reaching a QV of 100 or infinity, indicating full k-mer support from the input reads (Additional file 1: Figure S5). A range of evaluation tools confirmed the high quality of the final I002C assembly (Additional file 1: Figure S6-S7). Notably, 99.24% (5.93 Gbp) of the genome was flagged as haploid and error-free, while 44 Mbp (0.75%) were identified as duplicated or collapsed regions, and only 0.93 Mbp (0.02%) showed coverage-based errors detected by Flagger^7^ (Figure 2b). In addition, we curated platform-specific (HiFi, Duplex and ultra-long) coverage-bias and clipped-read regions from mapping of binned reads, and annotated them following previously described method^26^ (Additional file 1: Figure S8). The final polished I002C genome achieved an NG50 of 154.89 Mb for the maternal haplotype and 146.27 Mb for the paternal haplotype, comparable to previous high-quality human assemblies (Figure 2c). Merqury based quality assessment (k=21) resulted in genome wide QV scores of 82.05 for the maternal and 83.08 for the paternal I002C assemblies, exceeding other high-quality human genomes that are either published or publicly released (Figure 2c). To increase the sensitivity in homopolymer-rich regions, we also performed QV estimation using k-mer size of 31-mers. As expected, QV scores based on 31-mers were generally lower than those from 21-mers across all the genomes. Despite the overall decrease, I002C still exhibited higher QV values than the other genomes (Additional file 1: Figure S9). A haploid version of the I002C genome was also constructed by selecting the higher-quality assembly for each autosome and both sex chromosomes (X and Y) (Additional file 2: Table S5), resulting in a QV of 82.64 (k=21) and 70.16 (k=31). These results establish I002C as the highest quality human genome currently available.

**Figure 2:**
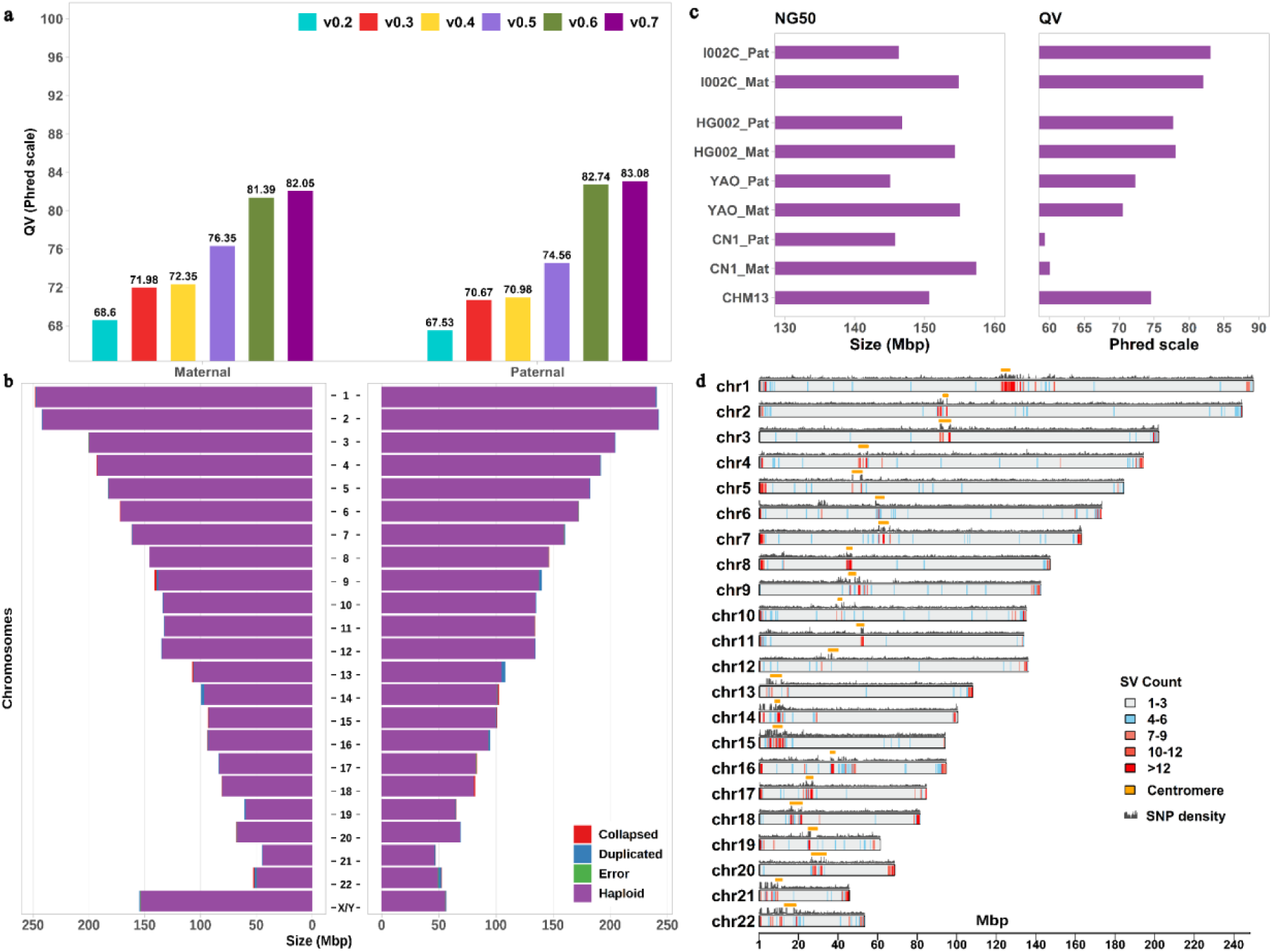
Overview of the haplotype-resolved I002C genome. **a)** Progression of I002C genome versions with polishing, showing improvements in QV scores assessed using hybrid k-mers. b) Assessment of haplotype reliability based on read mapping. For each chromosome, the left bar represents the maternal haplotype and the right bar the paternal haplotype. Regions marked as haploid (purple) are considered reliable, comprising over 99% of each haplotype on average. c) NG50 and QV (k=21) values for I002C haplotypes compared with other high-quality human genomes (HG002, YAO, CN1 and CHM13), ordered by year of publication or public release with most recent shown at the top. d) Genome-wide heterozygosity profile of I002C based on structural variant counts in 500 kb windows, overlaid with SNP density between maternal and paternal haplotypes. Higher diversity is observed near centromeric and subtelomere regions (highlighted by gold bars above the SNP density track).

### Annotation of Repeats, Centromeres and rDNAs

The haplotype-resolved genome enabled the annotation and analysis of repeat variation between the two haplotypes and the comparison with CHM13^2^. Genome-wide repeat content in I002C (including 22 autosomes and XY) was comparable to CHM13, with 53.29% of sequences being repetitive in the maternal and 53.43% in the paternal genomes, relative to 54.2% in CHM13 (Additional file 2: Table S6). We focused on the repeat families from peri/centromeric regions, where alpha-satellite (αSat), beta-satellite (βSat), gamma-satellite (γSat), and human satellites 1–3 (HSat1–3) together accounted for 5.09% and 5.96% of the maternal and paternal genomes, respectively, compared to 6.45% in CHM13 (Additional file 2: Table S7-S8). The annotated αSat arrays ranged from 1.40 to 6 Mbp in the maternal and 0.49 to 7.80 Mbp in the paternal genomes. These lengths, when compared to 165 samples from previous studies^28,29^, highlighted variation in both haplotypes and across individuals (Figure 3a).

**Figure 3:**
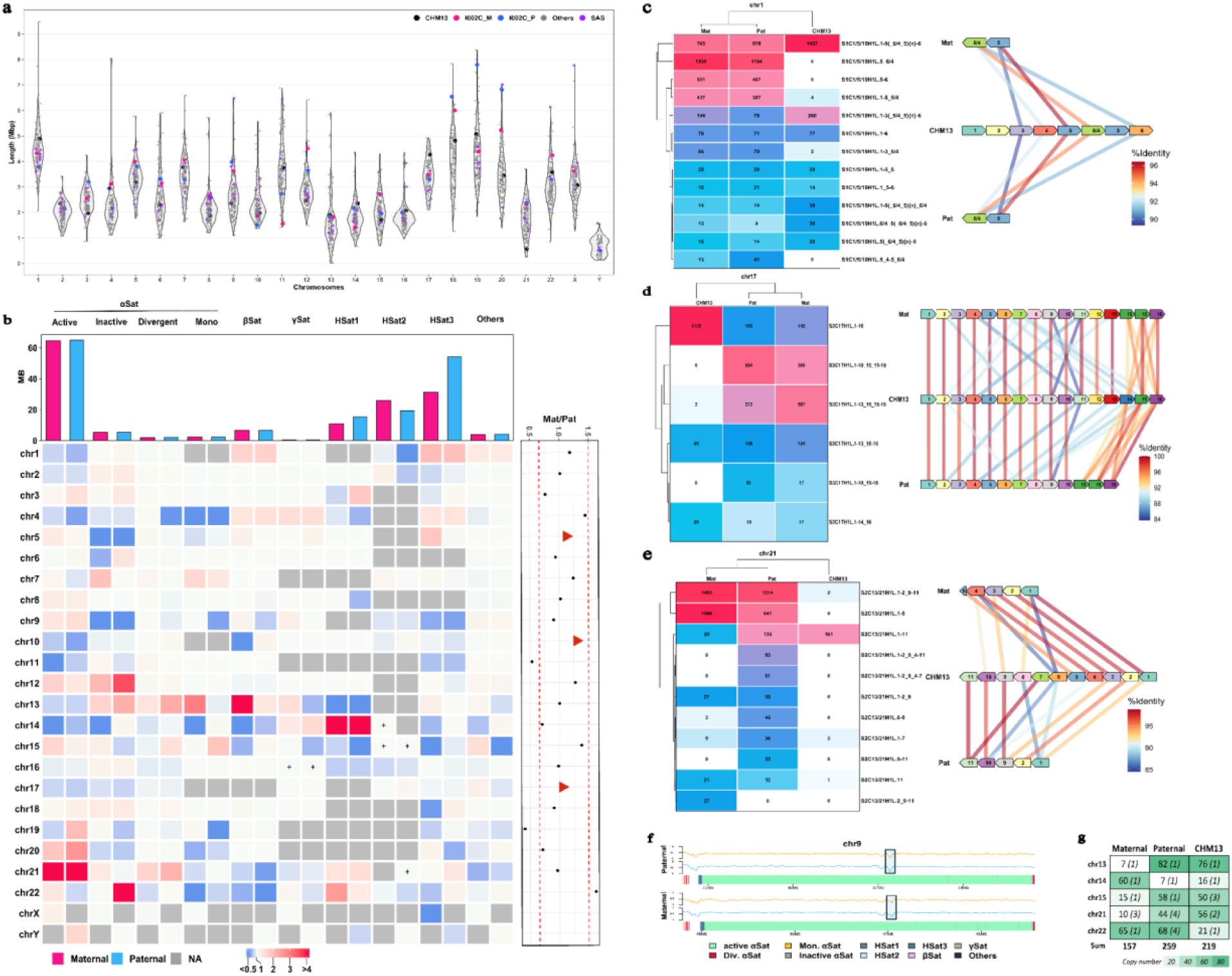
Peri/Centromeric satellite diversity, methylation, and rDNA variation among the I002C maternal, paternal and CHM13. **a)** Comparison of HOR lengths across I002C haplotypes and 165 human genomes. **b)** Heatmap showing the ratio of peri/centromeric satellite lengths in I002C haplotypes relative to CHM13. “+” denotes the presence of a satellite repeat in I002C but absence in CHM13. A dot plot compares active α-satellite (αSat) lengths between maternal and paternal haplotypes of I002C; chromosomes with >1.5-fold difference are marked with red dotted lines, and those with distinct HOR structures are indicated with triangles. **c-e)** Left: Heatmaps showing the abundances of top 10 most frequent HORs in each of I002C haplotypes and CHM13. Right: Structural variation and sequence similarity of the most abundant HORs. **f)** DNA methylation profiles across centromeric regions, measured from PacBio HiFi (blue) and Oxford Nanopore (yellow) data. Methylation levels (range 0–1) are averaged in 10 kb windows and normalized using adjacent windows. Centromere dip regions (CDRs) are indicated by shaded boxes. **g)** Heatmap of ribosomal DNA (rDNA) copy numbers, as well as major morph count [CN (morph count)] across I002C haplotypes and CHM13.

To assess the integrity and completeness of the annotated centromeric regions, we first mapped the 10 kb peri-centromeric regions from CHM13 to both I002C haplotypes and confirmed the presence of corresponding loci on each chromosome (Additional file 2: Table S9-S10). Next, all long-read datasets (HiFi, duplex, and ultra-long ONT) were aligned to the combined I002C reference, revealing uniform coverage across the annotated centromeric regions, consistent with accurate representation and assembly continuity (Additional file 1: Figure S10-S11). Centromere repeat annotation showed that the overall repeat composition in I002C is similar to CHM13, mainly consisting of αSat made up of high-order repeats (HORs) classified as active/live, inactive, divergent, and monomeric, along with βSat, γSat, and HSat1–3. At the individual chromosome level, we observed the variations in repeat families and repeat array lengths (Figure 3b). On average, the active αSat arrays in I002C are 1.33 times longer than in CHM13. In particular, nine autosomes from both haplotypes and the X and Y chromosomes of I002C are more than 1.8 times longer, with chromosome 21 showing the largest difference of 6.4-fold (Additional file 2: Table S11). However, these differences fall within the known range of human variation^23^ (Figure 3a). While satellite length generally differed, the largest difference was observed in the I002C paternal chromosome 14 HSat1, which spans 1.8 Mbp compared to 143.8 Kbp in CHM13. In contrast, the maternal chromosome 14 monomeric αSat measured 6.1 Kbp, substantially shorter than the 91.6 Kbp in CHM13. Despite these length differences, most peri/centromeric satellites are conserved between the I002C and CHM13 genomes, although some are uniquely present or absent. For example, maternal chromosome 14 and paternal chromosome 21 carry around ∼2 Kbp of HSat2 not found in CHM13, and both I002C haplotypes also include about ∼2 Kbp of HSat2 on chromosome 15 and 55 bp of γSat on chromosome 16 not present in CHM13 (Figure 3b, Additional file 1: Figure S12-S15).

To further explore the αSat HOR array structure between genomes, we compared centromeres of chromosome pairs between I002C and CHM13. This analysis showed differences in HOR organization between CHM13 and I002C genomes, including HOR SVs that are deviated from canonical patterns ^30,31^ (Additional file 2: Table S12). Notably, some HOR monomers found in I002C were completely absent from CHM13. For instance, CHM13 chromosome 1 was primarily composed of 6-mer HORs such as S1C1/5/19H1L.1-5(_6/4_5){n}-6, S1C1/5/19H1L.1-6 while I002C haplotypes contained distinct 2-mer variants like S1C1/5/19H1L.5_6/4 and S1C1/5/19H1L.5-6, which were missing from CHM13 (Figure 3c). The active HOR array of chromosome 17 in CHM13 mainly consists of the canonical 16-mer, whereas I002C haplotypes contain a mix of variant 16-mers along with 15-, 14-, and 13-mers. Notably, a novel 13-mer HOR variant (S3C17H1L.1-10_15_15-16) was found in both I002C haplotypes but not in CHM13 (Figure 3d). In contrast, chromosome 21 in both I002C haplotypes was primarily made up of 5-mer variants, compared to the canonical 11-mer in CHM13. A 5-mer variant (S2C13/21H1L.1-5) observed in I002C was completely absent in CHM13, and additional HORs unique to the paternal haplotype were not found in either the maternal genome or CHM13 (Figure 3e). Despite these differences, monomers within these HOR variants remain sequence-identical, suggesting structural polymorphism without extensive sequence divergence.

Similar differences were observed among the I002C maternal and paternal haplotypes (Additional file 1: Figure S16). Overall, 52% of the unique HOR SVs to I002C were specific to either the maternal or paternal haplotype. The dominant αSat HORs also differed between haplotypes—for instance, chromosome 5 had an 8-mer and a 4-mer; chromosome 10 showed a 6-mer and an 8-mer; and chromosome 17 carried a 16-mer and 13-mers in maternal and paternal haplotypes, respectively (Figure 3b). Together, these findings highlight the polymorphic nature of centromeric components among different human genomes and across haplotypes within the same genome.

Taking advantage of single-molecule long-read sequencing, which preserves native DNA methylation signals, we investigated methylation patterns across centromeric regions of both haplotypes. To identify putative kinetochore regions within the centromeres, we analyzed methylation patterns for signatures associated with centromere function. A characteristic centromeric methylation dip—marked by a sharp decrease in methylation levels within the active HOR subregions relative to their flanking sequences-was observed consistently across all chromosomes (Figure 3f, Additional file 1: Figure S17). The size of the methylation dip ranged from 0.1 to 0.4 Mb, consistent with previous reports in CHM1 and CHM13^32^ (Additional file 1: Figure S18).

Ribosomal DNA (rDNA) arrays located on the short arms of acrocentric chromosomes are among the most challenging regions to assemble due to their high repeat content, extensive segmental duplications, and complex satellite structures. The total rDNA copy number (CN) in I002C was estimated to be 416 for the diploid genome using digital PCR, which was consistent with estimations from the k-mer-based approach^2^. Based on the coverage of binned rDNA reads, we estimated 157 and 259 CNs for maternal and paternal haplotypes, respectively. Chromosome-level rDNA CNs were inferred by integrating read clustering with coverage-based and fluorescence in situ hybridization (FISH)-based estimates, and further cross-validated using independent FISH results (Additional methods: rDNA). Comparison of the estimated CN with the CN observed in the assembled genome revealed that the CNs of rDNAs were underrepresented for most of the chromosomes, except for the maternal chromosomes 13 and 21 (Additional file 2: Table S13). The maternal chromosome 21 rDNA sequence was found to be completely assembled in the initial assembly, supported by the coverage based estimates, FISH estimates, and full coverage of overlapping ultra-long reads. As for maternal chromosome 13, the reassembled rDNA sequence was nearly identical to the assembled genome in terms of both sequences and CNs, despite the absence of the full coverage of the region by overlapping ultra-long reads. The rDNA sequences from the assembled genome were retained for the maternal chromosomes 13 and 21. As for the remaining acrocentric chromosomes, due to the difficulty of resolving base-level variation among highly similar rDNA units, rDNA morphs, defined as representative sequences of one or more nearly identical units, were identified through further read clustering (Additional methods: **rDNA**). Arrays were reconstructed using consensus morph sequences scaled by their estimated CNs. These reconstructed arrays were then incorporated into the final assembly by replacing the originally assembled rDNA segments (Figure 3g, Additional file 2: Table S14).

In general, the typical rDNA array structure consisted of consecutive ∼45 kb rDNA units, flanked by a ∼4 kb rDNA fragment upstream and a ∼6 kb rDNA fragment downstream, and further bordered by the distal and proximal junction sequences (DJ, PJ). In total, we identified 19 major rDNA morphs that are different with >200 edit distance, including 7 maternal and 12 paternal morphs in I002C. Comparison with the 8 major morphs in CHM13 revealed sequence diversity, with all major morphs in I002C showing >200 edit distance^33^ from their CHM13 counterparts (Additional file 1: Figure S19). Despite this divergence, several large SVs were shared between I002C and CHM13 major morphs when compared to the canonical rDNA reference (Additional file 1: Figure S20). For example, a 1,530 bp deletion in the tandem repeat block 1 (TR1) is shared among I002C Mat_chr22_A, chm13-model15c and chm13-model22a morphs. In contrast, a 988 bp deletion in the long repeat (LR) region present in chm13-model15a and chm13-model15b was not observed in I002C major morphs. Additionally, we identified a ∼900 kbp deletion in the TR2 region of I002C Pat_chr22_B1 and Pat_chr22_B2 that is not present in CHM13 major morphs.

### Chromosome Y analysis

Assembling the human Y chromosome to a T2T chromosome remains challenging due to its highly repetitive architecture, especially within large heterochromatic regions and was only recently completed^19^. The I002C assembly provides a T2T, gapless Y chromosome from a South Asian individual, representing a valuable addition to the current catalogue of Y chromosome diversity^34^. The total length of I002C-Y is 56.6 Mb, consistent with other recent Y chromosome assemblies ranging from 45 to 85 Mb^34^ (Additional file 1: Figure S21). Notably, it is comparable in size to other South Asian individuals, including HG03009 (58.8 Mb), HG02492 (51.66 Mb), HG03732 (51.5 Mb), and HG03492 (51.2 Mb).

Comparative analysis of I002C-Y and T2T-Y from CHM13 (v2.0) showed concordance in overall structure (Additional file 1: Figure S22), with the largest size differences observed in ampliconic and heterochromatic subregions (Additional file 2: Table S15). Specifically, the AMPL2 subregion in I002C-Y is 11.7% (375 kb) shorter, the centromeric HET1_centro2 is 18.5% (327 kb) longer, HET2_DYZ19 is 52.7% (140 kb) longer, and HET3_Yq12 is 18.1% (6.25 Mb) shorter than their counterparts in T2T-Y. The multi-megabase variation observed in Yq12 is expected, as this region is composed of alternating repeat arrays known to be the largest heterochromatic block in the human genome and exhibits extensive inter-individual variability^34^. Sequence identity between subregions of I002C-Y and T2T-Y exceeds 97% for most regions, with the average identity being 96.65%. The most divergent region is HET2_DYZ19, with 89.4% identity, followed by two centromeric regions HET1_centroDYZ3 (93.9%) and HET1_centro2 (94.5%). In addition, we identify two large inversions when compared to T2T-Y, one in AMPL6 and the other one in AMPL7, and the one in AMPL6 was also observed when comparing to GRCh38 (Additional file 1: Figure S22). These inversions belong to palindromes P5 and P1, respectively, and are noticed in other recent Y chromosome assemblies, including South Asian ones^34^.

Finally, haplogroup assignment based on I002C-Y sequence places this individual in the R-Y6 lineage, a Y-chromosome haplogroup prevalent in South Asian populations and estimated to have emerged ∼4,000–4,500 years ago, underscoring the deep-rooted paternal ancestry in the region.

### Structural Variation in I002C Relative to CHM13 and Between the Paternal and Maternal Haplotypes

In addition to variations in peri/centromeric repeats, whole-genome comparison between I002C haplotypes and CHM13 identified 14,943 structural variants (SVs; ≥50 bp) spanning ∼19.50 Mbp in the maternal and 14,243 SVs spanning ∼18.74 Mbp in the paternal genome (excluding large repeats >10kb), with ∼95% of these being insertions and deletions (Additional file 2: Table S16–S17). Comparing I002C SVs against the combined SVs from public databases, including gnomAD^35^, HGSVC^28,36^, and HPRC^7^, showed that approximately 13.63% (3,236) of SVs were found to be unique to I002C (Additional file 2: Table S18). These I002C-specific SVs were primarily duplications and deletions (∼34% each), followed by insertions (30%) (Figure 1; Additional file 2: Table S18). Further repeat annotation showed that 72% of these unique SV regions overlapped with various repeat types (Additional file 2: Table S19).

Comparative analysis of the two complete haplotypes revealed approximately 2.90 million SNVs, 329,328 small indels (<50 bp), and 16,308 SVs. The heterozygosity rate, calculated as the number of SVs per 500 kb, showed that highly divergent regions are enriched in centromeric and subtelomeric regions (Figure 2d). In these regions, the SV density was more than four fold higher, and the SNV density more than two fold higher, compared to the genome-wide average (Additional file 1: Figure S23).

### Population-specific Reference Reduces Variant Calling Bias

To evaluate the impact of reference bias on read mapping and variant calling (SNV and SV), we compared four complete haploid reference genomes: three I002C haplotypes (maternal, paternal, and haploid version including mitochondrial DNA) and CHM13, using both short-read (NGS) and long-read (ONT) sequencing data. We randomly selected a subset of publicly available NGS data from the 1000 Genomes Project (1KGP)^23^, comprising 50 individuals from each of five South Asian (SAS) subpopulations (BEB, GIH, ITU, PJL, STU) (Additional file 2: Table S20). Additionally, we used ONT data from the Singapore HELIOS study^37^, including 26 Chinese, 31 Indian, and 37 Malay individuals, as well as NGS data from the same study including 39 Chinese, 39 Indian and 42 Malay individuals. Raw sequencing reads from these individuals were mapped to each reference genome, and variants were identified from each alignment.

For NGS data, we observed that the rate of uniquely mapped reads was marginally higher on CHM13 in both SAS populations and Singaporeans compared to either the maternal or paternal I002C haplotype. In SAS, the median of the rate reduced 0.09%, 0.02% and 0.06% for maternal, paternal and haploid, respectively, while in Singaporean samples, it reduced 0.02%, 0.05% and 0.02%, respectively. However, for long read (ONT) data of the Asian samples from Singapore the mapping rate is significantly lower on CHM13 than I002C (Additional file 1: Figure S24). The median of the rate increased 0.12% for maternal, 0.06% for paternal and 0.14% for haploid (Figure 4a). Interestingly, in the same SAS samples with short-read data, the number of detected SVs was generally lower when using the I002C haplotypes compared to CHM13 (Figure 4b, Additional file 1: Figure S25). An exception was observed in the PJL subgroup, where a slightly higher number of SVs was identified using the paternal or haploid genome. A similar trend was observed in long-read data and short-read data across the three Singaporean ethnic populations (Chinese, Malays and Indians), although the number of SVs was slightly higher in Chinese samples mapped to the maternal genome in long-read data or to the paternal or haploid genome in short-read data (Additional file 1: Figure S25).

**Figure 4:**
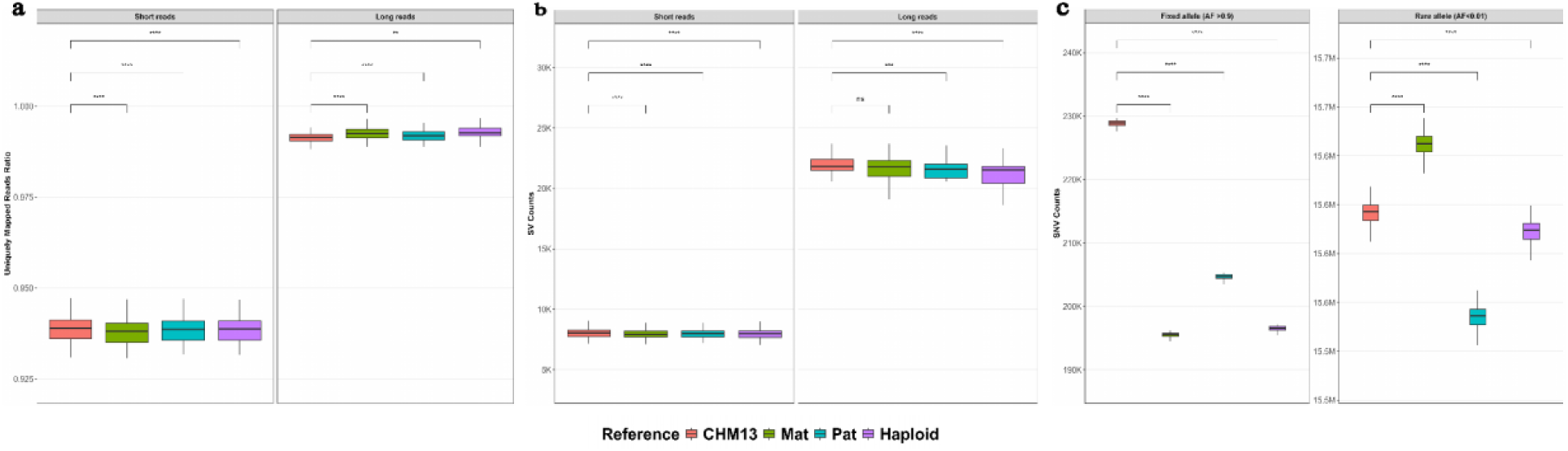
Comparative performance of I002C and CHM13 references in South Asian genomes. **a)** Mapping statistics of short and long-read sequencing data from SAS samples against I002C and CHM13 references. **b)** Allele frequency spectrum showing differences in the number of fixed or nearly fixed variants (AF > 0.9) and rare alleles (MAF < 0.01) across references. c) Comparison of structural variant counts detected using I002C and CHM13 references. The paired Wilcoxon test was used to assess differences in medians between the two groups. Significance levels: ns (P > 0.05), * (P ≤ 0.05), ** (P ≤ 0.01), *** (P ≤ 0.001), **** (P ≤ 0.0001).

To evaluate reference bias in variant calling at both individual and population cohort levels, variants were identified using deepvariant from mapping files of SAS, Singapore Indian, Chinese and Malay samples. For the SAS population including the Singapore Indian samples, an average of 25.76 million bi-allelic SNVs were identified across four reference genomes (Additional File 2: Table S21). However, allele frequency profiles differed significantly, especially for fixed or nearly fixed variants (AF > 0.9) between CHM13 and I002C genomes (Figure 4c). Using CHM13 as reference, 230,031 high-frequency variants were detected, compared with an average of 199,918 when using the I002C assemblies (maternal, paternal, and haploid) (Additional File 2: Table S21). In contrast, I002C references yielded a greater number of rare alleles (MAF < 0.01), consistent with improved representation of population-matched haplotypes. This allele frequency spectrum, indicating reduction in fixed variants and an enrichment of rare alleles, suggests that the I002C assemblies better capture the genetic landscape of the SAS population compared to CHM13 (Figure 4c). Similar results were observed for Singapore Chinese and Malay population (Additional file 2: Table S22-S23, Additional file 1: Figure S26).

We then assessed SVs that were specifically called by using CHM13 as reference (CHM13-specific SVs) after projecting their coordinates to GRCh38. SVs were merged across all SAS samples (250 samples with 183,811 SVs) and all Singaporean samples (94 samples with 138,909 SVs for long reads data, 120 samples with 91039 SVs for short reads data), respectively. Of these SVs, 21.2% (39,051) from SAS short reads data, 21% (19,082) and 15.4% (21,353) from Singaporeans short reads or long reads data were unique to CHM13. Of these CHM13-specific SVs, 5.5% in SAS short-read data, 9.4% and 18.8% in Singaporeans short- and long-read data were shared across at least 50% of the population. These additional SVs identified using CHM13 as the reference are likely variants that are either fixed or highly prevalent in the SAS or Singaporean populations and thus also present in the I002C haplotype genomes, while others may also be truly private to CHM13. Notably, 71 and 152 clinically relevant genes^38^ were affected by fixed or common SVs observed in over 90% of SAS or Singaporean individuals, respectively, indicating potential population-specific relevance in clinical interpretation. Notably, the gene TSC2—included in the ACMG SF v3.2^39^ list of actionable genes recommended for opportunistic screening was affected by an SV present in 99% of the SAS population or 94% of Singaporean individuals. Additionally, two other ACMG genes, TRDN and STK11, were affected by SVs present in 100% of Singaporean individuals. Further, OPRM1 and DPYD, both included in the Clinical Pharmacogenetics Implementation Consortium (CPIC) gene panel, were associated with SVs commonly found in 87% of SAS samples and 91.5% of Singaporeans. These SVs, particularly the clinically relevant ones, would not be called by using I002C as a reference, eliminating the need to do subsequent clinical annotation.

### Genome-wide Analysis of Allelic DNA Methylation

Besides the analysis of methylation patterns within the centromere regions, we also analyzed the genome-wide methylation pattern in the I002C genomes. Single-base resolution DNA methylation patterns were extracted from both Nanopore R10.4.1 and PacBio Revio data. The results showed high concordance between the two data, with a Pearson correlation coefficient of 0.953 when analyzed using 1000 bp sliding windows with 100 bp steps (Additional file 2: Table S24).

To enable parental-origin–aware haplotype-resolved methylation profiles, we assigned sequencing reads to maternal and paternal haplotypes using parental-specific k-mers. This analysis identified 70,400 single-CpG differentially methylated loci (DMLs) between paternal and maternal haplotypes. These DMLs were not randomly distributed, but instead clustered into regions with consistent haplotype-specific methylation. Such differentially methylated regions (DMRs) may have functional significance, and their relevance to disease development is exemplified by well-characterized imprinting disorders, including Prader-Willi syndrome (OMIM 176270)^40–42^, Angelman syndrome (OMIM 105830)^41,43^, Beckwith-Wiedemann syndrome (OMIM 130650)^44,45^, Silver-Russell syndrome (OMIM 180860)^45,46^, Temple syndrome (OMIM 616222)^47^, Kagami–Ogata syndrome (OMIM 608149)^48^, transient neonatal diabetes mellitus (OMIM 601410)^49^, and pseudohypoparathyroidism Ib (OMIM 603233)^50^. Using these known imprinting regions as benchmarks, we calibrated our thresholds and identified 968 autosomal DMRs (Additional file 1: Figure S27), of which 86.4% overlapped regulatory elements such as promoters and enhancers (Additional file 2: Table S25). To evaluate the robustness of our approach, we intersected these DMRs with 93 previously reported imprinting regions supported by multiple studies^51–56^. Reassuringly, all 46 overlapping regions (Additional file 1: Figure S28, Additional file 2: Table S26) matched the reported methylated allele, demonstrating strong concordance. The remaining reported regions were not detected, likely due to the stringent criteria applied (≥97.5% methylation difference at each DML and ≥5 DMLs per DMR).

Intersecting the 968 DMRs with experimentally validated promoter and first exon regions in GRCh38 revealed 2,229 transcripts (596 genes) potentially regulated by these DMRs (Additional file 1: Figure S29). In the I002C cell line, 203 transcripts (67.2% of those expressed) exhibited monoallelic expression, and 142 (70.0% of 203) showed concordant allele-specific methylation and transcription, corresponding to 94 genes (Additional file 2: Table S27). In addition, among the 596 candidate genes, 53 have previously been reported as imprinted^57^, including 22 genes directly linked to the eight imprinting disorders (Additional file 2: Table S28), highlighting the sensitivity of our approach and its potential in suggesting novel candidates of imprinting genes.

### Gene Annotation

To provide a comprehensive set of gene and transcript annotations, we first mapped Gencode annotations (v48) to the I002C maternal and paternal genome assemblies using liftOver^58^ and liftoff^59^. This process successfully transferred 98.92% of annotated transcripts to the I002C maternal and 96.41% to paternal assembly, with an overall 99.39% of annotated transcripts to the I002C (Additional file 2: Table S29).

To obtain a more complete set of genome annotation, we generated long read RNA-Seq data from I002C and used Bambu (v3.10.1)^60^ to build additional transcript models (Figure 5a). We observed that the population-matched reference genome (I002C) resulted in higher alignment rates and lower alignment errors than GRCh38 and CHM13 (Additional file 2: Table S30). Using a more conservative threshold (NDR=0.3), we identify 6190 novel transcripts in the maternal genome, and 5832 in the paternal genome, with more than 21,000 novel transcripts when a sensitive threshold is used (NDR=1, Additional file 2: Table S31). We then combined reference annotations from the maternal and paternal genomes to obtain a single haploid reference genome annotation for I002C), with 24.5% (NDR = 0.3) of transcripts being expressed in the I002C cell line (Figure 5b, c). In addition, 209 annotated transcripts in v48 are re-discovered by RNA-seq analysis, which failed liftOver. This integrated Lifting and long read RNA-seq approach improved annotation completeness, leading to the final numbers of 386477 transcript counts and 79221 genes for maternal haplotype, and 376337 transcripts and 76937 genes for paternal haplotype, when the more conservative threshold (NDR=0.3) is used (Additional file 2: Table S31).

**Figure 5.**
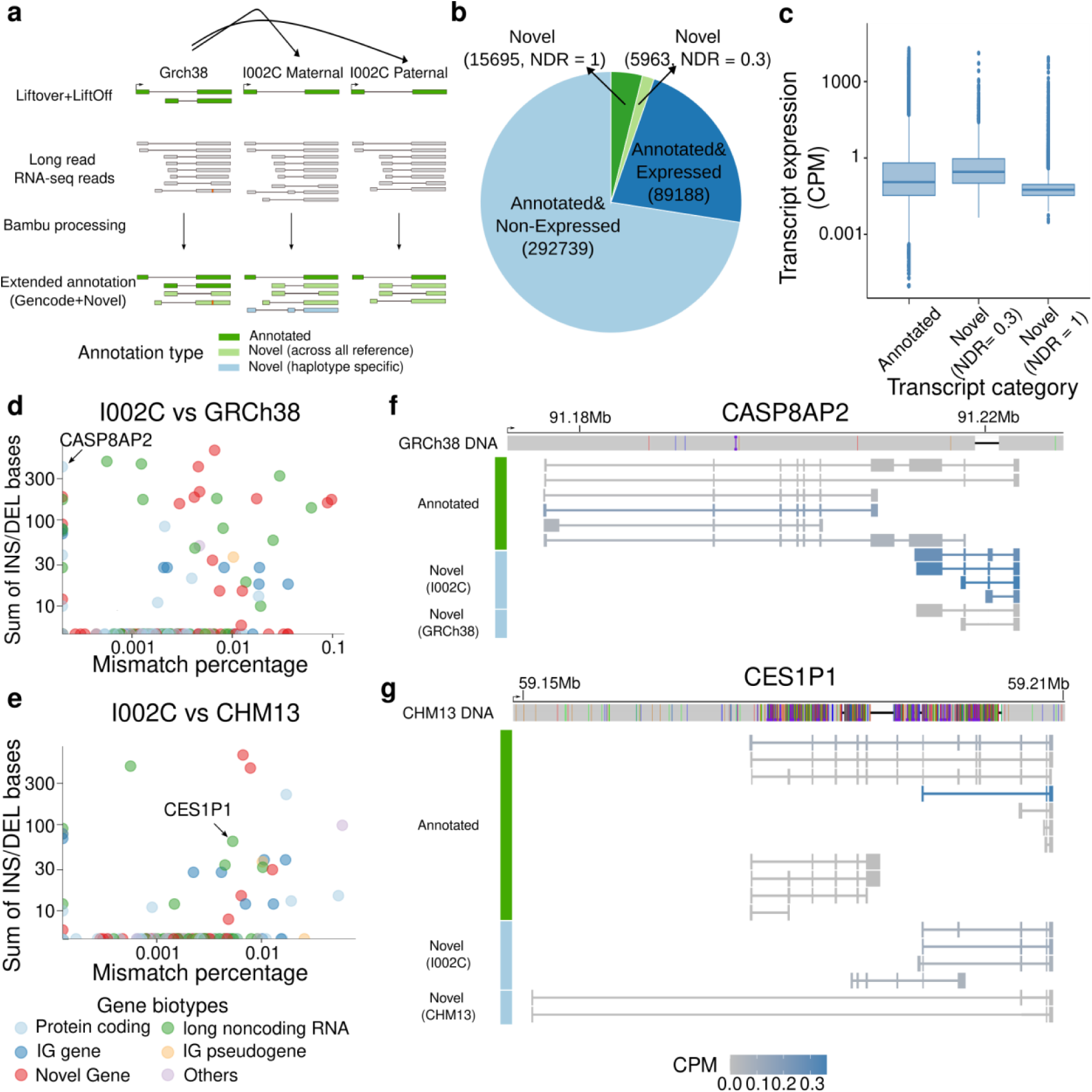
Gene annotation using long-read RNA-seq reads. **a)** Schematic of RNA-seq processing pipeline: 1) Firstly, we performed liftOver and liftoff of annotations from GRCh38 to I002C Maternal and Paternal haplotype separately; 2) We then processed long-read RNA-seq reads with Bambu using GRCh38, I002C Maternal and Paternal as reference respectively; 3) Extended annotation obtained from Bambu can be categorised by annotated, or novel (shared across all references), novel transcripts specific to individual haplotype **b)** Pie chart of I002C haplotype transcript number categorised by annotation and expression level in RNA-seq analysis. Light blue represents annotated and non-expressed (CPM = 0) transcript, dark blue represents annotated and expressed (CPM > 0) transcript, light green represents novel transcript identified with NDR = 0.3, and dark green represents additional novel transcript identified with NDR = 1 **c)** Boxplot of transcript expression in count per million (CPM) for annotated transcript, novel transcript identified when NDR = 0.3, and novel transcript identified when NDR = 1 **d-e)** Scatter plot of percentage of mismatch vs sum of insertion and deletion bases (small indels < 5bp filtered before sum) when I002C haplotype-specific novel transcripts aligned against d) GRCh38 and e) CHM13 reference, focusing on transcripts containing at least 3 exons and CPM greater than 0.04. Note that for each gene, the transcript showing largest deviation (rank by sum of insertion and deletion bases and then followed by mismatch percentage) is kept in the plot for visualization. Light blue represents protein coding genes, dark blue represents IG genes, green represents long noncoding RNAs, orange represents IG pseudogenes, pink represents novel genes, and purple represents other pseudogene types. **f-g)** Top: f) GRCh38 and g) CHM13 DNA sequence aligned against I002C maternal reference. Bottom: From top to bottom, transcript annotations for annotated transcripts, novel transcript specific to I002C maternal haplotype, novel transcripts specific to f) GRCh38, or g) CHM13. Transcript annotations are colored by CPM, with darker values indicating higher CPM.

Next we studied the genome divergence between I002C, GRCh38 and CHM13 for I002C-specific transcript annotations. We aligned the transcript cDNA sequences both annotated and newly discovered by Bambu with different NDR thresholds from I002C against GRCh38 and CHM13, and calculated the number of alignment mismatch positions to estimate the sequence variation, and the number of deletion and insertion positions to estimate structural variation. While the majority of transcripts are shared between I002C and GRCh38 (96%), we observe a number of transcripts with substantial sequence variation (Figure 5d-g). Among these are for example novel transcripts for the CASP8AP2 and CES1P1 gene which contain a previously unannotated exon that is missing from the GRCh38 and CHM13 reference genomes respectively (Figure 5f, g). These novel transcripts will not be quantified if GRCh38 or CHM13 is used as reference, highlighting the value of a population-matched reference genome for transcriptomic analysis.

## Discussion

In this study, we present the first telomere-to-telomere (T2T), fully phased diploid genome assembly of an Indian individual using a trio sample set of Indian ethnicity from Singapore. The majority of Singaporean Indians descend from migrants during the British colonial period, primarily of Telugu and Tamil origin, with some contributions from Northern groups such as Sikhs and Pathans^12^. It is therefore assuring to see that I002C is well embedded within the 5 SAS population samples from the 1000 Genomes Project. More interestingly, I002C is genetically located in the middle area between the Western and Northern SAS samples (GIH and PJL) and the Eastern, Southeastern and Southern SAS samples (BEB, ITU and STU). As such, the I002C genome reflects broader Indian subcontinental ancestry and offers value to individuals of similar backgrounds in South Asia and worldwide. Additionally, it contributes to improving representation for these South Asian populations in human genomic datasets.

Using high-quality long read data, trio-aware assembly, ultra long-read gap filling, and multi-platform polishing, we achieved complete, gapless assemblies of all the autosomes, sex chromosomes, and a fully resolved circular mitochondrial genome representative of the individual. The final assemblies reached NG50 values of over 154 Mb for maternal 146 Mb for paternal and Merqury QV scores exceeding 82 for both haplotypes, establishing I002C as the highest-quality diploid and haploid human genome assembled to date. To the best of our knowledge, the maternal chromosome 21 rDNA sequence of the I002C was the first fully assembled rDNA cluster sequence that has been published. Furthermore, I002C also contributes one of the few complete Y chromosome assemblies from a South Asian individual, expanding current knowledge of Y-linked diversity and structural variation.

As the first reference genome for the South Asian population, I002C highlights important features that remain incompletely characterized in many reference genomes. For example, in addition to providing the first fully assembled rDNA cluster on Chr21 (maternal), I002C also revealed 19 major rDNA morphs, compared to the 8 major rDNA morphs for CHM13. Comparison of I002C genome to CHM13 revealed that over 13% of SVs in I002C are novel, with many falling in repeat-rich regions, which are often sources of structural complexity in the genome. When comparing two I002C haplotypes, the analysis revealed 2.9 million SNVs, more than 329 thousands small indels and more than 16 thousands SVs. Many of these variants were concentrated in centromeric and subtelomeric regions, identifying them as hotspots of structural divergence. In addition, the fully resolved diploid genome enabled the generation of haplotype-resolved methylome and long read RNA expression data. This data enables the identification of parent-of-origin–aware differentially methylated regions (DMRs) that show high concordance with DMRs reported in previous studies, where parent-of-origin information was obtained using strand-seq, germline variant–based allelic determination, or special uniparental samples such as uniparental disomy, complete hydatidiform moles, and ovarian teratomas, which contain chromosomes from a single parent. We have further demonstrated that the DMRs within core transcription regulatory sequences showed high concordance with haplotype-resolved long read RNA expression data. These DMRs have captured many of known imprinting genes, particularly the imprinting disease-associated ones, and suggest additional candidates of novel imprinting genes. Together, these findings highlight the value of fully phased diploid genomes for exploring complex and polymorphic regions.

Our analysis has demonstrated that the I002C is a better reference than CHM13 for the genomic analysis of South Asian samples as well as other Asian samples by reducing population bias. For example, population-specific analysis, based on 250 randomly selected SAS individuals from the 1000 Genomes Project, showed comparable rates of uniquely mapped short reads across references. However, for Singapore samples, the mapping rate for long read sequencing data is higher for I002C than CHM13, which is consistent with the fact that I002C carries a good number of SVs that are unique or much more common in South Asian and Southeast Asian samples than other populations. These SVs are expected to have a negative impact on mapping long reads than short reads by using CHM13 as reference. In addition, analysis of SNVs showed that allele frequency profiles differed between references, with CHM13 yielding more high-frequency variants and I002C assemblies capturing more rare alleles, consistent with improved representation of population-matched haplotypes. Similar patterns were also observed for Singapore Chinese and Malay populations. The number of called SVs was reduced when mapping SAS samples to I002C instead of CHM13, reflecting the presence of population-specific sequences in I002C. Reduction of variant calls by using I002C as a reference can help to reduce the effort on identifying potential pathogenic variant(s) for genomic diagnostic analysis of SAS patients. The additional SVs identified with CHM13 as a reference included variants in medically relevant genes, such as TSC2, which was affected in 99% of SAS samples. Such additional SVs, particularly the medically relevant ones, may lead to misdiagnosis for SAS patients. Together, these findings suggest that the I002C haplotypes more accurately represent the genomic architecture of the SAS population and also serve as a suitable reference for Southeast Asian population particularly the Singapore Malay and Chinese samples. This underscores the value of population-specific assemblies in reducing reference bias and improving accuracy in downstream analyses such as variant discovery and association studies.

By providing a complete, ancestry-relevant reference genome, I002C improves the representation of South Asians in genomic research and enhances the resolutions of both structural and regulatory genomic variation. In addition to supporting more accurate variant interpretation, it refines epigenetic and imprinting analyses. As a first step of our efforts on generating reference genomes for South and Southeast Asian populations, I002C will serve as a linear backbone for constructing locally and Asian relevant pangenomes and contribute to the growing set of population-specific references that are critical for reducing global reference bias and advancing equitable precision medicine.

## Methods

### Sample selection and sequencing

The details of sample selection, Pacbio HiFi, Oxford Nanopore duplex, ultra-long and Omni-C data generation are described in the previous study^61^.

### Karyotyping

Cells were cultured in RPMI 1640 medium (#11875-093, Gibco, USA) supplemented with 10% fetal bovine serum (#A5670801, Gibco) and 1% penicillin–streptomycin (#15140-122, Gibco), and maintained in suspension at 37 °C in a humidified incubator with 5% CO₂. Approximately 5 million cultured cells were treated with KaryoMAX™ Colcemid™ Solution in HBSS (#15210040, Gibco) for 2 hours to induce mitotic arrest. Following standard protocols from *The AGT Cytogenetics Laboratory Manual* (4^th^ Edition)^62^, routine G-banded metaphase preparations were performed^62^.

Karyotype analysis of 20 cells was conducted using the Ikaros karyotyping software (Metasystem) at a 450-band resolution, following the International System for Human Cytogenomic Nomenclature (ISCN 2020) guidelines.

### RNA Sequencing

RNA was extracted using the QIAGEN PAXgene Blood RNA Kit according to the manufacturer’s instructions. Total RNA was used as input material for the RNA sample preparations. rRNA was removed from total RNA by using TIANSeq rRNA Depletion Kit (Animal). Fragmentation was carried out using divalent cations under elevated temperature in the First Strand Synthesis Reaction Buffer. First strand cDNA was synthesized using random hexamer primer and reverse transcriptase, followed by the synthesis of the second strand by adding buffer and dNTPs (with dTTP replaced by dUTP). The synthesized double-stranded cDNA was treated by terminal repair, A-tailing, adaptor ligation, and PCR enrichment after fragment selection. The strand-specific library was obtained by selecting cDNA fragments of 370-420bp by purification with AMPure XP beads. The qualified library was pooled and sequenced on the Illumina NovaSeq X Plus platform with a 150 × 2 paired-end configuration.

### Nanopore cDNA sequencing

Total RNA was extracted from the I002C lymphoblastoid cell line using the Bioline Isolate II RNA Mini Kit (Meridian Biosciences) according to manufacturer’s instructions. Total RNA quantification, purity and integrity assessment was performed using the Qubit 3.0 Fluorometer and Qubit BR RNA Assay Kit (ThermoFisher Scientific), NanoDrop 2000 spectrophotometer (ThermoFisher Scientific) and Agilent 4200 TapeStation system with the RNA screentape assay kit (Agilent). 39µg of total RNA was brought forward to poly(A)+ RNA enrichment using the Dynabeads mRNA Purification Kit (ThermoFisher Scientific) according to manufacturer’s instructions. Nanopore cDNA-PCR library was generated from 50ng of poly(A)+ RNA using the SQK-PCB114.24 kit, (Oxford Nanopore Technologies) according to manufacturer’s instructions. Briefly, complementary strand synthesis and strand switching were performed on input full-length poly(A)+ RNA using kit-supplied oligonucleotides, 16-cycle PCR amplification using primers containing 5’ tags to generate a double-stranded cDNA library, followed by ligase-free attachment of rapid sequencing adaptors. Final cDNA library was loaded onto a FLO-PRO114M R10.4.1 flowcell (Oxford Nanopore Technologies) and sequenced on a PromethION P24 A100 instrument, PromethION release version 25.03.7, Dorado server 7.8.3, equivalent to Dorado release 0.9.2, with live-basecalling in super-accurate (SUP) mode and barcoding trimming.

### WGS Analysis by Next Generation Sequencing (NGS)

WGS PCR-free libraries for the gDNA samples were prepared using MGIEasy FS PCR-free DNA library prep set (MGI; Cat. no. 1000013455) as per manufacturer’s protocol. The WGS libraries were quality checked by the HSD 1000 Screentape and reagent (Agilent Technologies; cat. no. 5067-5585 and 5067-5584) on the Tapestation 4200 (Agilent Technologies). The single strand circularized DNA product of the library was measured with Qubit™ ssDNA HS Assay Kit (Thermo Fisher Scientific; Cat. no. Q10212) on the Qubit fluorometer 4 (Thermo Fisher Scientific). DNA nanoball (DNB) was made for each library from the single strand circularized DNA using the DNBSEQ DNB Rapid Make Reagent Kit (FCL PE150) (MGI; 1000028453) and concentration of DNB were determined by the Qubit™ ssDNA HS Assay Kit (Thermo Fisher Scientific; Cat. no. Q10212) on the Qubit fluorometer 4 (Thermo Fisher Scientific). DNBs were pooled based on the sequencing data requirement, and the pooled DNBs were sequenced 150bp paired-end (PE150) on the MGI DNBSEQ-T7 sequencer using the DNBSEQ-T7RS High-throughput Sequencing Kit (FCL PE150) V1.0 sequencing reagent (MGI; Cat. no. 1000016106) and DNBSEQ-T7RS Sequencing Flow Cell (T7 FCL) (MGI; Cat. no. 1000016269).

### Genome Assembly and Polishing

The initial assemblies were constructed using PacBio HiFi and uncorrected ONT ultra-long reads in trio mode with Verkko (v1.4.1)^20^ and Hifiasm (v0.19.6-r595)^21,22^ algorithms. Contigs were further assembled into full-length chromosomes through manual curation with 3D-DNA^63^ and a template-based approach using the CHM13 genome to determine the correct contig order for joining.

To support gap filling and manual curation, ONT ultra-long reads were also corrected using HERRO^64^ and assembled in trio mode with Verkko. In addition, all long reads were haplotype-binned and reassembled using both Hifiasm and LJA (v0.2)^65^.

Some gaps were falsely introduced during the scaffolding process, as confirmed by long-read and Hifiasm and LJA reassembled contig alignments to the assembly gap regions. These false gaps appeared as large deletions in the alignments and were subsequently removed. We considered all the remaining gaps to be real gaps. Real gaps were first closed using TGS-GapCloser (v1.2.1)^25^. The remaining gaps were manually filled by aligning corrected ultra-long reads, Verkko HERRO reads assembly, as well as Hifiasm and LJA reassemblies to the 10 kb flanking regions around each gap.

Alignments were manually examined in IGV (v2.15.2)^66^, and only reciprocal best hits were selected for gap filling. Telomeres were patched and extended using ultra-long, duplex and HiFi reads containing the (TTAGGG)n motif at either end, identified using Tel_Sequences^67^. These reads were aligned to the first or last 250 kb of each chromosome, and consensus sequences were generated using SPOA (v4.1.4)^68^ for patching, resulting in 23 gapless telomere-to-telomere chromosome level assemblies for each haplotype.

The assembly was polished through two rounds of automated processes using the T2T polishing pipeline^26^, followed by one round of manual curation and an additional round of ONT-based assembly patching. In brief, haplotype-specific binned long reads (HiFi, duplex and ultra-long) reads were mapped to their respective haplotypes using winnowmap2 (v2.03)^69^ with default platform-specific parameters. PCR-free MGI short reads were aligned using BWA MEM (v0.7.17-r1188)^70^, and the resulting sorted HiFi and short alignment files were merged using SAMtools (v1.18)^71^. SNV + indel (<50 bps) were called using DeepVariant (v1.6.1)^72^ in “HYBRID_PACBIO_ILLUMINA” for merged HiFi, short reads and in “ONT_R104” mode for duplex reads, while ultra-long reads were processed with PEPPER-Margin-DeepVariant (v0.8)^73^. SVs were identified using Sniffles2 (v2.2)^74^ and subsequently refined and merged using the Iris (v1.0.5) - Jasmin (v1.1.5)^75^. These variants were filtered using merfin (v1.0)^76^ and used for genome polishing with bcftools (v1.10.2)^77^. During the manual curation round, variants were identified as previously described and carefully examined in IGV for genome polishing. Subsequently, remaining erroneous regions within 1 kb window were merged using BEDTools (v2.30)^78^, and 1 kb flanking sequences, including the erroneous regions, were extracted. These sequences were aligned to the corresponding haplotype assemblies from corrected ultra-long assemblies using Minimap2 (v2.28)^79^ to identify reciprocal best matches. Only error-free regions from the ultra-long assemblies were used for patching (Additional methods: **Genome assembly**).

### Evaluation of Assembly

Assembly quality (QV) was assessed using hybrid k-mers derived from HiFi and MGI reads, with completeness and phasing statistics calculated using Merqury (v1.3)^27^. Assembly errors and overall quality were also evaluated using CRAQ (v1.0.9-alpha)^80^, which utilizes read clipping information, and Inspector (v1.3)^81^, which detects both large (SV) and small-scale (SNV) errors based on reads-to-assembly alignments. Additionally, coverage-based tools such as Flagger (v0.3.3)^7^ and GCI (v1.0)^82^ were employed to identify errors based on coverage inconsistencies. Haplotype assemblies were further validated using Omni-C contact maps, which were visualized in Juicebox (v1.11.08)^83^.

### Repeat Annotation and Centromere Analysis

Repeat annotation for both haplotypes was performed using RepeatMasker (v4.1.5)^84^ with a curated repeat library from Dfam (v3.7)^85^. Centromere alpha satellite (αSat) monomers were identified using Hum-AS-HMMER^86^, and high-order repeats (HORs) were classified as active/live, inactive, divergent and monomeric based on previous studies^30,31^. HSat2 and HSat3 were annotated following established methods^31^ while other centromeric satellites were identified by parsing RepeatMasker output. Structural variants within active centromeric HORs were predicted using Stv^87^.

### Genome Comparison Between I002C and CHM13

To minimize the identification of mapping-biased false structural variants (SVs), all centromeric regions, heterochromatin, and repeats ≥10 kb were masked with ’N’. Each I002C chromosome from the maternal and paternal haplotypes was aligned to the corresponding CHM13 chromosome using minimap2 with the ‘-x asm5 -r2k’ options for the contiguous alignments. SVs were identified using Syri (v1.6.3)^88^ and svim-asm (v1.0.3)^89^. A consensus SV set for each haplotype was generated by merging SVs detected by both tools using Jasmine with ‘max_dist=1000 min_support=2”. The final combined SV set for the I002C genome was generated by merging both haplotypes using jasmine ‘max_dist=1000 min_support=1’.

To identify I002C-specific SVs, they were compared against publicly available SV datasets from gnomAD (v4.1)^35^, HGSVC (v1–3)^28,36^, and HPRC^7^. Since SV coordinates for I002C and HPRC were based on CHM13, whereas gnomAD and HGSVC SVs were based on GRCh38, the latter were lifted over to CHM13 using Picard LiftOverVCF (v2.26.9)^90^. Finally, SVs from public datasets were merged using Jasmine with the parameters max_dist=1000 min_support=1.

### Assessing Reference Genome Suitability

***Read mapping performance*** We compared mapping performance for both short-read NGS data and long-read ONT data across three complete haploid reference genomes: I002C maternal haploid (includes chrY and mitochondrial genome), I002C paternal haploid (includes chrX and mitochondrial genome), and CHM13 (v2.0).

For short-read analysis, we randomly selected a subset of South Asian (SAS) individuals from the 1000 Genomes Project (1KGP), comprising 50 individuals from each SAS subpopulation: BEB, GIH, ITU, PJL, and STU. Moreover, the NGS data from the Singapore HELIOS cohort were used to facilitate a comprehensive evaluation . The corresponding CRAM files were converted to FASTQ format using samtools (v1.18). Paired-end reads from each sample were aligned separately to the four reference genomes using BWA-MEM (v0.7.17). Duplicate reads were marked using GATK MarkDuplicates (v4.1.8.1)^91^.

For long-read analysis, we used ONT data from the Singapore HELIOS cohort. The reads were aligned to each reference using minimap2 (v2.28). Mapping statistics were collected from each BAM file using samtools, and the percentage of uniquely mapped reads was calculated using the formula (NMapped reads -NMQ0) / NMapped reads.

***Variant calling*** Short-read variants were identified with DeepVariant from duplicate-marked read alignments to four reference genomes. Long-read variants were called using Clair3^92^ with the “r941_prom_sup_g5014” model. Bi-allelic SNPs were extracted with the bcftools view “-m2 -M2 -v snps -f PASS” options. Variants with read depth (DP) ≥ 10 and genotype quality (GQ) > 10 were retained. Allele frequency (AF) were calculated separately for SAS and Singapore HELIOS cohorts (Chinese and Malay). Variants with AF > 0.9 were classified as fixed or nearly fixed, while those with AF < 0.01 were considered rare.

***Structural variant calling*** To further assess reference genome performance, structural variants (SVs) were called from the NGS alignments using Manta (v1.6.0)^93^, and from ONT alignments using Sniffles (v2.6.2). To evaluate the reference specific SVs, we first projected SVs on GRCh38 coordinates using CrossMap (v0.7.0)^94^. The chain files from I002C to GRCh38 were generated following the nf-LO pipeline^95^.

A combined SV set for each haplotype was generated by merging SV across all samples using Jasmine (v1.1.5) with ‘max_dist=1000’. The same merging strategy was used to merge three reference-based SV files to identify the reference-specific SVs.

### Gene Annotation

To annotate genes on the I002C maternal and paternal genome assemblies, as part of the gene annotation pipeline (Additional file 1: Figure S30; Additional methods: **Gene annotation**), we first lifted gene models from GRCH38 (Gencode annotations (v48)) to our assembly using liftOver (conda version 377). Chain files for both haplotypes were generated from GRCH38 to the I002C assemblies using the nf-LO pipeline. This pipeline uses minimap2 for alignment and paf2chain to convert alignments into UCSC-style chain files. The resulting chain files were applied with liftOver to project gencode annotations (v48) onto the I002C assemblies. In parallel, we also applied liftoff (v1.6.3), which directly maps gene models by aligning genomic features rather than relying on precomputed chain files, allowing a more flexible transfer in structurally divergent regions. Finally, annotations from both approaches were merged using a set of predefined rules (Additional methods: **Gene annotation**), with the procedure giving greater weight to liftoff annotations.

Reference genome annotations were then extended by using long read RNA-seq data. First, reads were aligned to each haplotype-specific assembly using Minimap2. Transcript models were then constructed and quantified using Bambu (v3.10.1)^60^. Here we run Bambu with two thresholds for high precision (NDR=0.3) and high sensitivity (NDR=1). We then extracted cDNA sequences for Bambu extended annotations using each haplotype-specific assembly and aligned against GRCh38 and CHM13 separately to identify potential sequence variation for annotated transcripts and also to check whether novel transcripts identified using each haplotype-specific assembly is haplotype-specific or shared in GRCh38 or CHM13. For each transcript, we calculated the mismatch percentage and the sum of insertion and deletion bases to measure the sequence variation. Note that for this analysis, small insertion or deletion less than 5 bp were not counted, to minimize the impact of small alignment errors. Alignment of these transcript models were then processed with Bambu with trackReads set to TRUE, and without discovery and quantification. We then defined novel transcripts as haplotype-specific if there does not exist any equalMatches (i.e., not complete splicing junction chain match) to extended annotations obtained when using GRCh38 or CHM13 (including both annotated and novel transcripts), CPM > 0.04, and containing at least 3 exons.

### Ancestry Analysis

Local ancestry inference for the I002C genome was performed by calling variants relative to the GRCh38 reference using dipcall (v0.3)^96^. The resulting variant set was merged with biallelic SNPs from 1000 Genomes Project (1KGP) samples representing African (AFR), American (AMR), East Asian (EAS), European (EUR), and South Asian (SAS) super-populations. Ancestry was inferred at the super-population level using RFMix (v2.03-r0)^97^.

For the PCA analysis, we performed variant calling for the individual I002C and the 1KGP South Asian (250 SAS) samples using DeepVariant, followed by joint genotyping with GLnexus (v1.4.1)^98^. To minimize potential bias from close relatives, we excluded first- and second-degree related samples, resulting in a final dataset of 248 individuals. Variants were retained if they met the thresholds of allelic quality (AQ) > 20, read depth (DP) > 10, and genotype quality (GQ) > 20. To obtain an independent set of SNPs for PCA, we excluded long-range linkage disequilibrium (LD) regions (projected from GRCh38) and applied additional filtering with the parameters: --maf 0.1 --geno 0.03 --hwe 1e-3 --indep-pairwise 1500 150 0.2 --snps-only --autosome. PCA was then conducted in PLINK (v1.9)^99^ using the final dataset of 247 SAS samples and I002C.

### Methylation

Genome-wide and centromeric DNA methylation profiling for I002C genomes was performed using PacBio HiFi and ONT simplex reads with platform specific pipelines. To ensure reliable quantification, methylation ratios were calculated only for CpG sites supported by at least four methylation-informative reads (containing MM/ML tags). To evaluate platform concordance, Pearson correlation coefficients were computed for methylation levels across shared CpG sites and predefined genomic windows. CDR for each chromosome was manually examined and annotated as described in the previous study^31^. Haplotype-specific differentially methylated loci (DMLs) were identified by aligning parental reads (excluding homozygous reads) to the GRCh38 reference, requiring a minimum methylation difference of 97.5% between the haplotypes and at least five informative reads per site. Differentially methylated regions (DMRs) were defined as regions containing at least five DMLs and majority of DMLs within this region to have consistent methylated allele. To identify candidate genes potentially regulated by allelic methylation, DMRs were intersected with core promoter and first exon regions of experimentally validated transcripts (Additional methods: **Methylation**).

### ChrY analysis

The subregions of I002C-Y were annotated using subregion definitions from T2T-Y. The T2T-Y subregions were mapped onto I002C-Y using minimap2 (v2.28) with the parameters “-x asm20 -Y -N1 --cs -c --eqx --paf-no-hit.” To assess sequence conservation, we computed average sequence identity for each subregion based on pairwise alignments between I002C-Y and T2T-Y using nucmer (v3.1)^100^ with “-- maxmatch -l 100” parameters. For each subregion, identity was calculated as a length-weighted average across all alignments. Additionally, I002C-Y subregions were mapped onto GRCh38 chromosome Y (NCBI GCF_000001405.40) using the same method as described above, and the alignment identities were also computed accordingly.

To further characterize the paternal lineage, we determined the Y chromosome haplogroup using short read data aligned to the I002C-Y assembly. Haplogroup assignment was performed using Yleaf (v3.2.1)^101^ by comparing SNPs in the non-recombining region of the Y chromosome to those in the YFull phylogenetic reference tree (v10.01)^102^.

## Data availability

The sequencing reads have been submitted to NCBI under Project ID PRJNA1150503. The I002C assemblies are available at https://github.com/LHG-GG/I002C. Additionally, the publicly available data resources used in this study are listed in Additional file 2: Table S32, along with their corresponding access URLs.

## Acknowledgement

We want to thank the anonymous volunteers who contributed their blood samples to this study. We would also like to thank the A*STAR Microscopy Platform (AMP) and the AMP staff, especially Dr Goh Wah Ing, for the consultation, advice, and technical support. AMP is funded by Singapore’s Agency for Science, Technology & Research (A*STAR) through core funding. We also want to thank Oxford Nanopore and PacBio for their long read sequencing support. This work was supported by the funding from the A*STAR and PRECISE and the high-performance computing resources provided by the A*STAR Computational Resource Centre (A*CRC).

## Contributions

JJ.L. and M.Š. conceived the project. W.YK.H., contributed to the sample recruitment and collection. YY.S. contributed to establishing and growing cell lines. F.Y., H.W.L., S.T.L., C.C.K., contributed to library preparation and sequencing. P.S. designed the pipeline. P.S., J.L. and D.H. performed genome assemblies. P.S. performed genome polishing, evaluation, centromere and repeat analysis. J.L., P.S. performed read binning. P.S., J.L., Q.J., Z.L., L.W., L.V., D.H., X.Z., K.K., M.Š., and JJ.L. performed manual curation. P.S., Q.J. performed in-silico rDNA CN estimation. Z.L. performed dPCR experiment. Z.L., D.M., performed FISH experiments and analysis. Q.J. performed rDNA assembly. Z.L. performed methylation analysis. X.Z. performed ancestry analysis. L.V. performed chromosomeY analysis. J.L. and Y.C. performed gene annotation. P.S., W.L., performed reference justification analysis. E.W., W.Z., contributed to project management. J.C.C., contributed to the recruitment of HELIOS samples. P.T., and JJ.L., secured the funding. P.S., J.L., Q.J., L.W., Z.L., L.V., Y.C., M.Š., J.G., and JJ.L. drafted the manuscript with contribution from all authors. M.Š. and JJ.L. supervised the project and provided mentorship.

## Ethics declarations

### Ethics approval and consent to participate

All the participants were provided with informed consent for sample collection, and usage including making data publicly available via databases. Sample collection and usage were approved by SingHealth Centralised Institutional Review Board (IRB Reference: 2024– 069) and A*STAR Institutional Review Board (IRB Reference: 2019-118). All experimental methods comply with the Helsinki Declaration.

### Consent for publication

Not applicable.

## Supplementary Information

Additional File1 - Supplementary figures S1-S30

Additional File2 - Supplementary tables S1-S33

Additional methods - Detailed methodology

## References

1. Schneider, V. A. et al. Evaluation of GRCh38 and de novo haploid genome assemblies demonstrates the enduring quality of the reference assembly. Genome Res 27, 849–864 (2017).

2. Nurk, S. et al. The complete sequence of a human genome. Science 376, 44–53 (2022).

3. HG002. MarBL (2025).

4. Yang, C. et al. The complete and fully-phased diploid genome of a male Han Chinese. Cell Res 33, 745–761 (2023).

5. He, Y. et al. T2T-YAO: A Telomere-to-Telomere Assembled Diploid Reference Genome for Han Chinese. Genomics, Proteomics & Bioinformatics 21, 1085–1100 (2023).

6. Wang, T. et al. The Human Pangenome Project: a global resource to map genomic diversity. Nature 604, 437–446 (2022).

7. Liao, W.-W. et al. A draft human pangenome reference. Nature 617, 312–324 (2023).

8. Popejoy, A. B. & Fullerton, S. M. Genomics is failing on diversity. Nature 538, 161–164 (2016).

9. Fatumo, S. et al. Diversity in Genomic Studies: A Roadmap to Address the Imbalance. Nat Med 28, 243–250 (2022).

10. Yew, C.-W. et al. Genomic structure of the native inhabitants of Peninsular Malaysia and North Borneo suggests complex human population history in Southeast Asia. Human genetics 137, 161–173 (2018).

11. Wall, J. D. et al. The GenomeAsia 100K Project enables genetic discoveries across Asia. Nature 576, 106–111 (2019).

12. Wu, D. et al. Large-Scale Whole-Genome Sequencing of Three Diverse Asian Populations in Singapore. Cell 179, 736–749.e15 (2019).

13. Le, V. S. et al. A Vietnamese human genetic variation database. Human mutation 40, 1664– 1675 (2019).

14. Deng, L. et al. Analysis of five deep-sequenced trio-genomes of the Peninsular Malaysia Orang Asli and North Borneo populations. BMC Genomics 20, 842 (2019).

15. Piriyapongsa, J. et al. Pharmacogenomic landscape of the Thai population from genome sequencing of 949 individuals. Sci Rep 14, 30683 (2024).

16. Azmi, M., Chen, L., Idris, A. & Lu, Z. H. Genomic Insights of Bruneian Malays. 2022.06.01.492266 Preprint at 10.1101/2022.06.01.492266 (2022).

17. Bhattacharyya, C. et al. Mapping genetic diversity with the GenomeIndia project. Nat Genet 57, 767–773 (2025).

18. Ardiansyah, E. et al. Sequencing whole genomes of the West Javanese population in Indonesia reveals novel variants and improves imputation accuracy. Front. Genet. 15, (2025).

19. Rhie, A. et al. The complete sequence of a human Y chromosome. Nature 621, 344–354 (2023).

20. Rautiainen, M. et al. Telomere-to-telomere assembly of diploid chromosomes with Verkko. Nature Biotechnology 41, 1474–1482 (2023).

21. Cheng, H., Concepcion, G. T., Feng, X., Zhang, H. & Li, H. Haplotype-resolved de novo assembly using phased assembly graphs with hifiasm. Nat Methods 18, 170–175 (2021).

22. Cheng, H., Asri, M., Lucas, J., Koren, S. & Li, H. Scalable telomere-to-telomere assembly for diploid and polyploid genomes with double graph. Nature Methods 21, 967–970 (2024).

23. Auton, A. et al. A global reference for human genetic variation. Nature 526, 68–74 (2015).

24. Samuels, D. C. et al. Heterozygosity Ratio, a Robust Global Genomic Measure of Autozygosity and Its Association with Height and Disease Risk. Genetics 204, 893–904 (2016).

25. Xu, M. et al. TGS-GapCloser: A fast and accurate gap closer for large genomes with low coverage of error-prone long reads. GigaScience 9, giaa094 (2020).

26. Mc Cartney, A. M., et al. Chasing perfection: validation and polishing strategies for telomere- to-telomere genome assemblies. Nature Methods 19, 687–695 (2022).

27. Rhie, A., Walenz, B. P., Koren, S. & Phillippy, A. M. Merqury: reference-free quality, completeness, and phasing assessment for genome assemblies. Genome Biology 21, 245 (2020).

28. Logsdon, G. A. et al. Complex genetic variation in nearly complete human genomes. 2024.09.24.614721 Preprint at 10.1101/2024.09.24.614721 (2024).

29. Gao, S. et al. Centromere Landscapes Resolved from Hundreds of Human Genomes. Genomics Proteomics Bioinformatics 22, qzae071 (2024).

30. Miga, K. H. & Alexandrov, I. A. Variation and Evolution of Human Centromeres: A Field Guide and Perspective. Annual Review of Genetics vol. 55 583–602 (2021).

31. Altemose, N. et al. Complete genomic and epigenetic maps of human centromeres. Science 376, eabl4178.

32. Logsdon, G. A. et al. The variation and evolution of complete human centromeres. Nature 629, 136–145 (2024).

33. Šošić, M. & Šikić, M. Edlib: a C/C ++ library for fast, exact sequence alignment using edit distance. Bioinformatics 33, 1394–1395 (2017).

34. Hallast, P. et al. Assembly of 43 human Y chromosomes reveals extensive complexity and variation. Nature 621, 355–364 (2023).

35. Collins, R. L. et al. A structural variation reference for medical and population genetics. Nature 581, 444–451 (2020).

36. Ebert, P. et al. Haplotype-resolved diverse human genomes and integrated analysis of structural variation. Science 372, eabf7117 (2021).

37. Wang, X. et al. The Health for Life in Singapore (HELIOS) Study: delivering Precision Medicine research for Asian populations. medRxiv 2024.05. 14.24307259 (2024).

38. Wagner, J. et al. Curated variation benchmarks for challenging medically-relevant autosomal genes. Nat Biotechnol 40, 672–680 (2022).

39. Miller, D. T. et al. ACMG SF v3.2 list for reporting of secondary findings in clinical exome and genome sequencing: A policy statement of the American College of Medical Genetics and Genomics (ACMG). Genet Med 25, 100866 (2023).

40. Butler, M. G. Prader-Willi Syndrome: Obesity due to Genomic Imprinting. Curr Genomics 12, 204–215 (2011).

41. Buiting, K. et al. Epimutations in Prader-Willi and Angelman Syndromes: A Molecular Study of 136 Patients with an Imprinting Defect. Am J Hum Genet 72, 571–577 (2003).

42. Bieth, E. et al. Highly restricted deletion of the SNORD116 region is implicated in Prader– Willi Syndrome. European Journal of Human Genetics 23, 252–255 (2015).

43. Meng, L. et al. Truncation of Ube3a-ATS unsilences paternal Ube3a and ameliorates behavioral defects in the Angelman syndrome mouse model. PLoS genetics 9, e1004039 (2013).

44. Viljoen, D. & Ramesar, R. Evidence for paternal imprinting in familial Beckwith-Wiedemann syndrome. Journal of medical genetics 29, 221–225 (1992).

45. Chiesa, N. et al. The KCNQ1OT1 imprinting control region and non-coding RNA: new properties derived from the study of Beckwith–Wiedemann syndrome and Silver–Russell syndrome cases. Human molecular genetics 21, 10–25 (2012).

46. Ishida, M. New developments in Silver–Russell syndrome and implications for clinical practice. Epigenomics 8, 563–580 (2016).

47. Ioannides, Y., Lokulo-Sodipe, K., Mackay, D. J., Davies, J. H. & Temple, I. K. Temple syndrome: improving the recognition of an underdiagnosed chromosome 14 imprinting disorder: an analysis of 51 published cases. Journal of medical genetics 51, 495–501 (2014).

48. Kilich, G. et al. Kagami Ogata syndrome: a small deletion refines critical region for imprinting. NPJ Genomic Medicine 9, 5 (2024).

49. Yorifuji, T., Higuchi, S., Hosokawa, Y. & Kawakita, R. Chromosome 6q24-related diabetes mellitus. Clinical Pediatric Endocrinology 27, 59–65 (2018).

50. Hanna, P., Francou, B., Delemer, B., Jüppner, H. & Linglart, A. A novel familial PHP1B variant with incomplete loss of methylation at GNAS-A/B and enhanced methylation at GNAS-AS2. The Journal of Clinical Endocrinology & Metabolism 106, 2779–2787 (2021).

51. Court, F. et al. Genome-wide parent-of-origin DNA methylation analysis reveals the intricacies of human imprinting and suggests a germline methylation-independent mechanism of establishment. Genome Res 24, 554–569 (2014).

52. Joshi, R. S. et al. DNA Methylation Profiling of Uniparental Disomy Subjects Provides a Map of Parental Epigenetic Bias in the Human Genome. The American Journal of Human Genetics 99, 555–566 (2016).

53. Hernandez Mora, J. R., et al. Characterization of parent-of-origin methylation using the Illumina Infinium MethylationEPIC array platform. Epigenomics 10, 941–954 (2018).

54. Zink, F. et al. Insights into imprinting from parent-of-origin phased methylomes and transcriptomes. Nature genetics 50, 1542–1552 (2018).

55. Akbari, V. et al. Genome-wide detection of imprinted differentially methylated regions using nanopore sequencing. eLife 11, e77898.

56. Akbari, V. et al. Parent-of-origin detection and chromosome-scale haplotyping using long-read DNA methylation sequencing and Strand-seq. Cell Genomics 3, 100233 (2023).

57. Rosenski, J. et al. Atlas of imprinted and allele-specific DNA methylation in the human body. Nature Communications 16, 2141 (2025).

58. Kuhn, R. M., Haussler, D. & Kent, W. J. The UCSC genome browser and associated tools. Briefings in Bioinformatics 14, 144–161 (2013).

59. Shumate, A. & Salzberg, S. L. Liftoff: accurate mapping of gene annotations. Bioinformatics 37, 1639–1643 (2021).

60. Chen, Y. et al. Context-aware transcript quantification from long-read RNA-seq data with Bambu. Nature methods 20, 1187–1195 (2023).

61. Sarashetti, P., Lipovac, J., Tomas, F., Šikić, M. & Liu, J. Evaluating data requirements for high-quality haplotype-resolved genomes for creating robust pangenome references. Genome Biology 25, 312 (2024).

62. Lawce, H. J. & Brown, M. G. Cytogenetics: an overview. The AGT cytogenetics laboratory manual 25–85 (2017).

63. Dudchenko, O. et al. De novo assembly of the Aedes aegypti genome using Hi-C yields chromosome-length scaffolds. Science 356, 92–95 (2017).

64. Stanojević, D., Lin, D., Nurk, S., Florez de Sessions, P. & Šikić, M. Telomere-to-Telomere Phased Genome Assembly Using HERRO-Corrected Simplex Nanopore Reads. bioRxiv 2024.05.18.594796 (2024) doi:10.1101/2024.05.18.594796.

65. Bankevich, A., Bzikadze, A. V., Kolmogorov, M., Antipov, D. & Pevzner, P. A. Multiplex de Bruijn graphs enable genome assembly from long, high-fidelity reads. Nat Biotechnol 40, 1075–1081 (2022).

66. Thorvaldsdóttir, H., Robinson, J. T. & Mesirov, J. P. Integrative Genomics Viewer (IGV): high-performance genomics data visualization and exploration. Briefings in Bioinformatics 14, 178–192 (2013).

67. Sarashetti, P. T2T Sequences.

68. Vaser, R., Sović, I., Nagarajan, N. & Šikić, M. Fast and accurate de novo genome assembly from long uncorrected reads. Genome research 27, 737–746 (2017).

69. Jain, C., Rhie, A., Hansen, N. F., Koren, S. & Phillippy, A. M. Long-read mapping to repetitive reference sequences using Winnowmap2. Nature Methods 19, 705–710 (2022).

70. Li, H. Aligning sequence reads, clone sequences and assembly contigs with BWA-MEM. Preprint at 10.48550/arXiv.1303.3997 (2013).

71. Li, H. et al. The Sequence Alignment/Map format and SAMtools. Bioinformatics 25, 2078–2079 (2009).

72. Poplin, R. et al. A universal SNP and small-indel variant caller using deep neural networks. Nature Biotechnology 36, 983–987 (2018).

73. Shafin, K. et al. Haplotype-aware variant calling with PEPPER-Margin-DeepVariant enables high accuracy in nanopore long-reads. Nature Methods 18, 1322–1332 (2021).

74. Smolka, M. et al. Detection of mosaic and population-level structural variants with Sniffles2. Nature Biotechnology 42, 1571–1580 (2024).

75. Kirsche, M. et al. Jasmine and Iris: population-scale structural variant comparison and analysis. Nature Methods 20, 408–417 (2023).

76. Formenti, G. et al. Merfin: improved variant filtering, assembly evaluation and polishing via k-mer validation. Nature Methods 19, 696–704 (2022).

77. Danecek, P. et al. Twelve years of SAMtools and BCFtools. GigaScience 10, giab008 (2021).

78. Quinlan, A. R. & Hall, I. M. BEDTools: a flexible suite of utilities for comparing genomic features. Bioinformatics 26, 841–842 (2010).

79. Li, H. Minimap2: pairwise alignment for nucleotide sequences. Bioinformatics 34, 3094– 3100 (2018).

80. Li, K., Xu, P., Wang, J., Yi, X. & Jiao, Y. Identification of errors in draft genome assemblies at single-nucleotide resolution for quality assessment and improvement. Nature Communications 14, 6556 (2023).

81. Chen, Y., Zhang, Y., Wang, A. Y., Gao, M. & Chong, Z. Accurate long-read de novo assembly evaluation with Inspector. Genome Biology 22, 312 (2021).

82. Chen, Q., Yang, C., Zhang, G. & Wu, D. GCI: a continuity inspector for complete genome assembly. Bioinformatics 40, btae633 (2024).

83. Durand, N. C. et al. Juicebox Provides a Visualization System for Hi-C Contact Maps with Unlimited Zoom. Cell Systems 3, 99–101 (2016).

84. Smit, A., Hubley, R. & Green, P. RepeatMasker Open-4.0. 2013–2015. (2015).

85. Storer, J., Hubley, R., Rosen, J., Wheeler, T. J. & Smit, A. F. The Dfam community resource of transposable element families, sequence models, and genome annotations. Mobile DNA 12, 2 (2021).

86. Ryabov, F. HumAS-HMMER_for_AnVIL. (2024).

87. Ryabov, F. stv. (2024).

88. Goel, M., Sun, H., Jiao, W.-B. & Schneeberger, K. SyRI: finding genomic rearrangements and local sequence differences from whole-genome assemblies. Genome Biol 20, 277 (2019).

89. Heller, D. & Vingron, M. SVIM-asm: structural variant detection from haploid and diploid genome assemblies. Bioinformatics 36, 5519–5521 (2020).

90. Broad Institute. Picard Tools.

91. McKenna, A. et al. The Genome Analysis Toolkit: A MapReduce framework for analyzing next-generation DNA sequencing data. Genome Res 20, 1297–1303 (2010).

92. Zheng, Z. et al. Symphonizing pileup and full-alignment for deep learning-based long-read variant calling. Nature Computational Science 2, 797–803 (2022).

93. Chen, X., et al. Manta: rapid detection of structural variants and indels for germline and cancer sequencing applications. Bioinformatics 32, 1220–1222 (2016).

94. Zhao, H. et al. CrossMap: a versatile tool for coordinate conversion between genome assemblies. Bioinformatics 30, 1006–1007 (2014).

95. Chen, N.-C. & Vollger, M. nfLO.

96. Li, H. et al. A synthetic-diploid benchmark for accurate variant-calling evaluation. Nat Methods 15, 595–597 (2018).

97. Maples, B. K., Gravel, S., Kenny, E. E. & Bustamante, C. D. RFMix: A Discriminative Modeling Approach for Rapid and Robust Local-Ancestry Inference. Am J Hum Genet 93, 278–288 (2013).

98. Yun, T. et al. Accurate, scalable cohort variant calls using DeepVariant and GLnexus. Bioinformatics 36, 5582–5589 (2020).

99. Purcell, S. et al. PLINK: a tool set for whole-genome association and population-based linkage analyses. The American journal of human genetics 81, 559–575 (2007).

100. Kurtz, S. et al. Versatile and open software for comparing large genomes. Genome Biol 5, R12 (2004).

101. Ralf, A., Montiel González, D., Zhong, K. & Kayser, M. Yleaf: Software for Human Y-Chromosomal Haplogroup Inference from Next-Generation Sequencing Data. Molecular Biology and Evolution 35, 1291–1294 (2018).

102. Yfull. Haplogroup YTree v10.01.00. https://www.yfull.com/arch-10.01/tree/ (2022).

103. Koren, S. et al. Canu: scalable and accurate long-read assembly via adaptive k-mer weighting and repeat separation. Genome research 27, 722–736 (2017).

104. Schindelin, J., et al. Fiji: an open-source platform for biological-image analysis. Nature methods 9, 676–682 (2012).

105. Akalin, A. et al. methylKit: a comprehensive R package for the analysis of genome-wide DNA methylation profiles. Genome Biol 13, R87 (2012).

