## Additional File1 for "A Complete Telomere-to-Telomere Diploid Reference Genome for South Asian Population"

**Supplementary figures:**

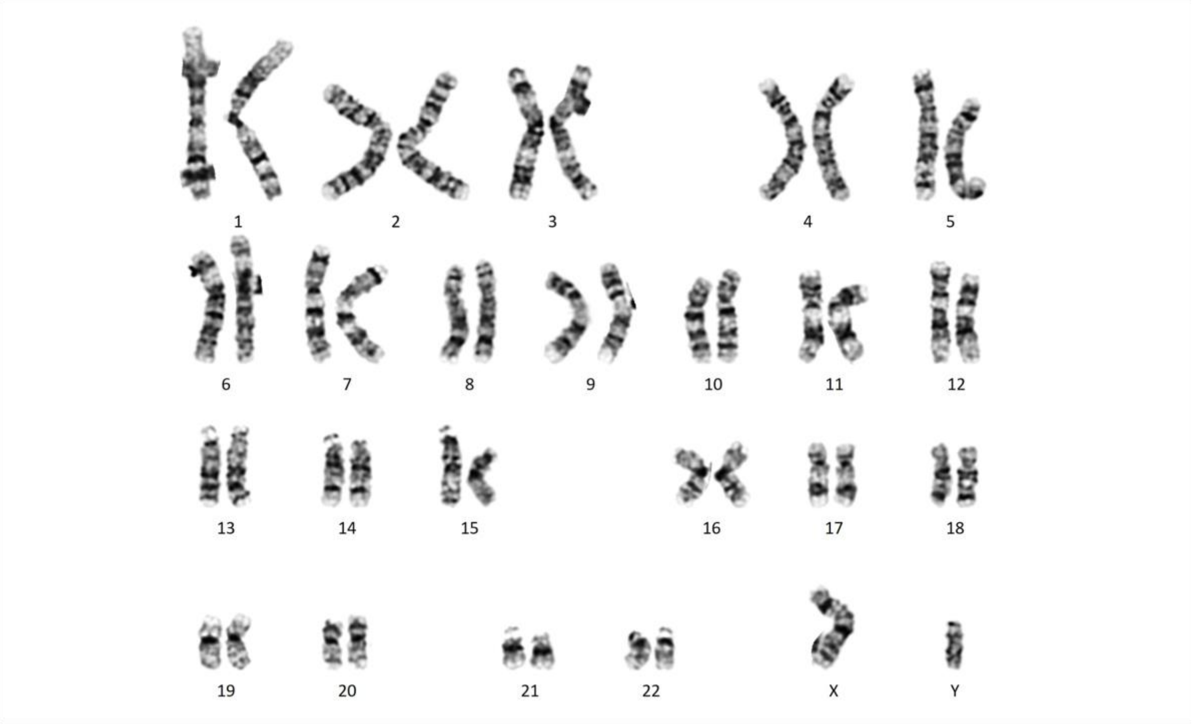


**Figure S1. Karyotype of the I002C cell line.**

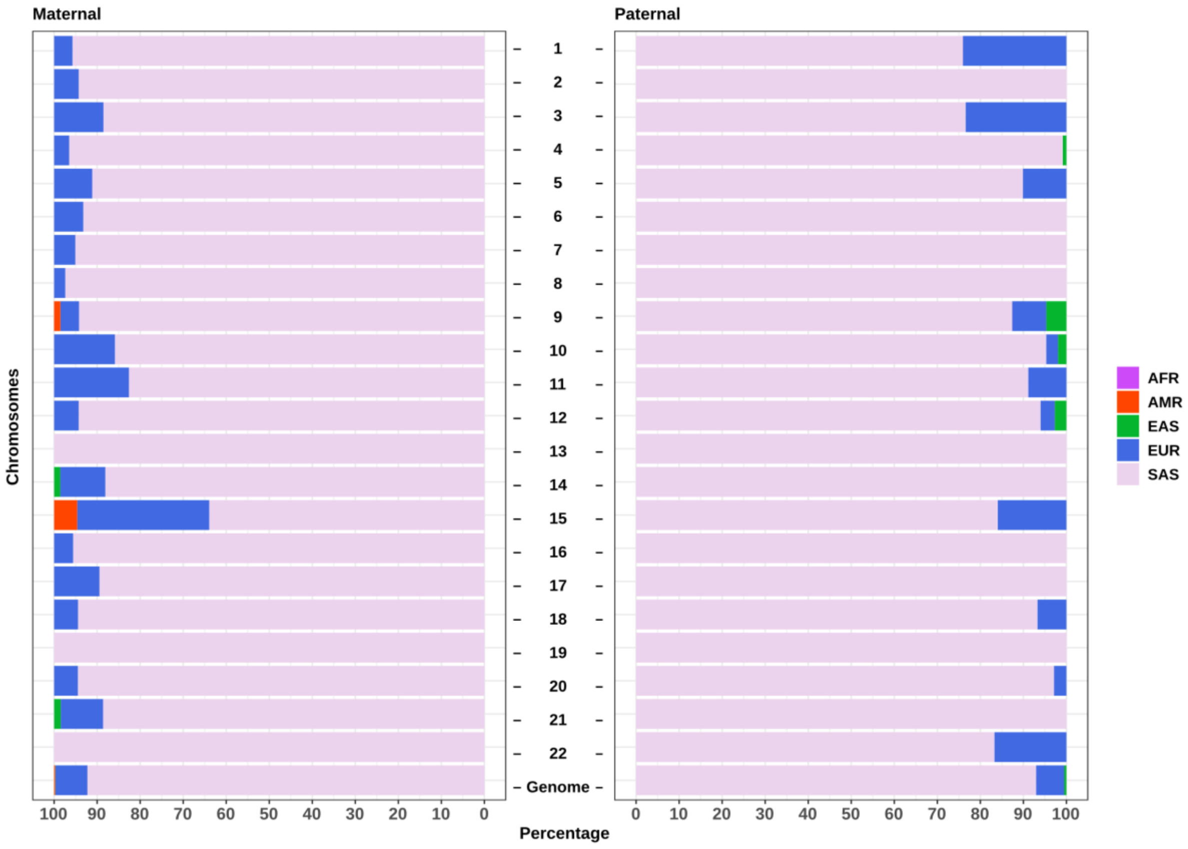


**Figure S2. Genomic distribution of ancestral components in I002C.**

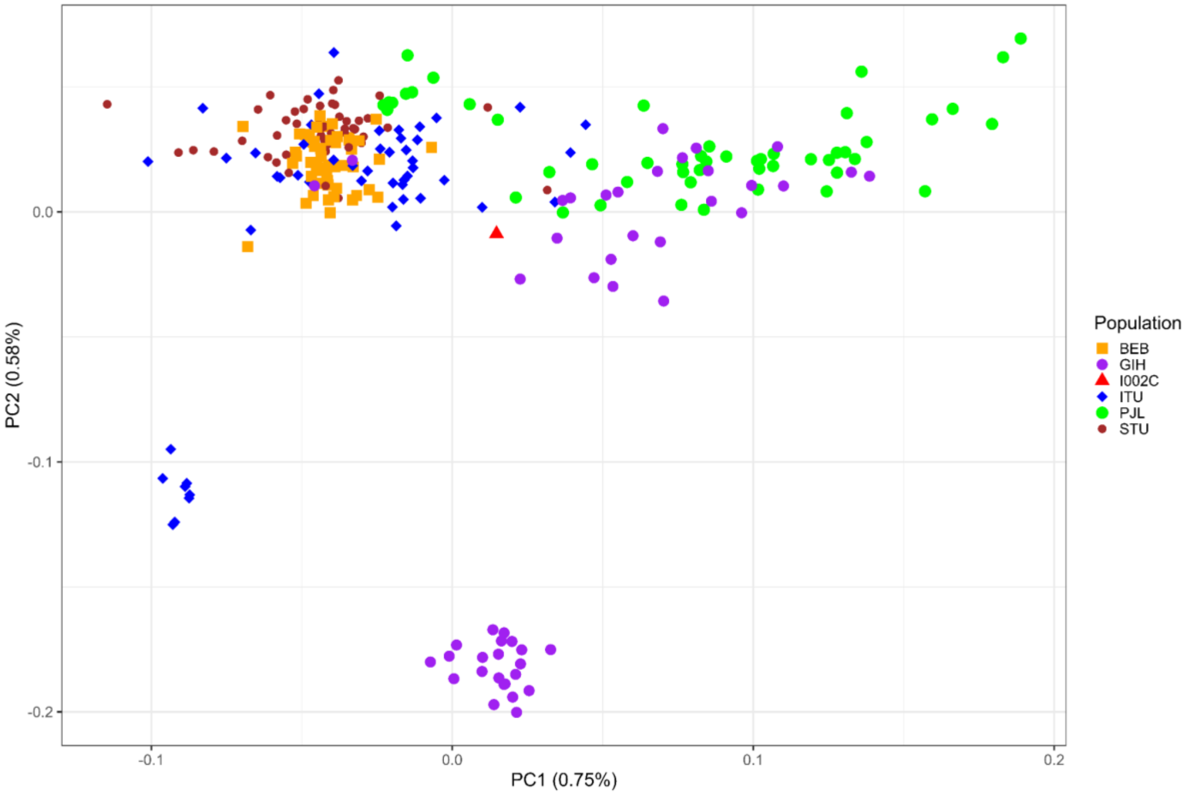


**Figure S3. PCA of I002C with 5 SAS subpopulations from 1KGP.** The plot showed GIH and ITU have two clusters, respectively. The I002C (red triangle) is closer to the PHL, and GIH sub-group, ITU sub-group.

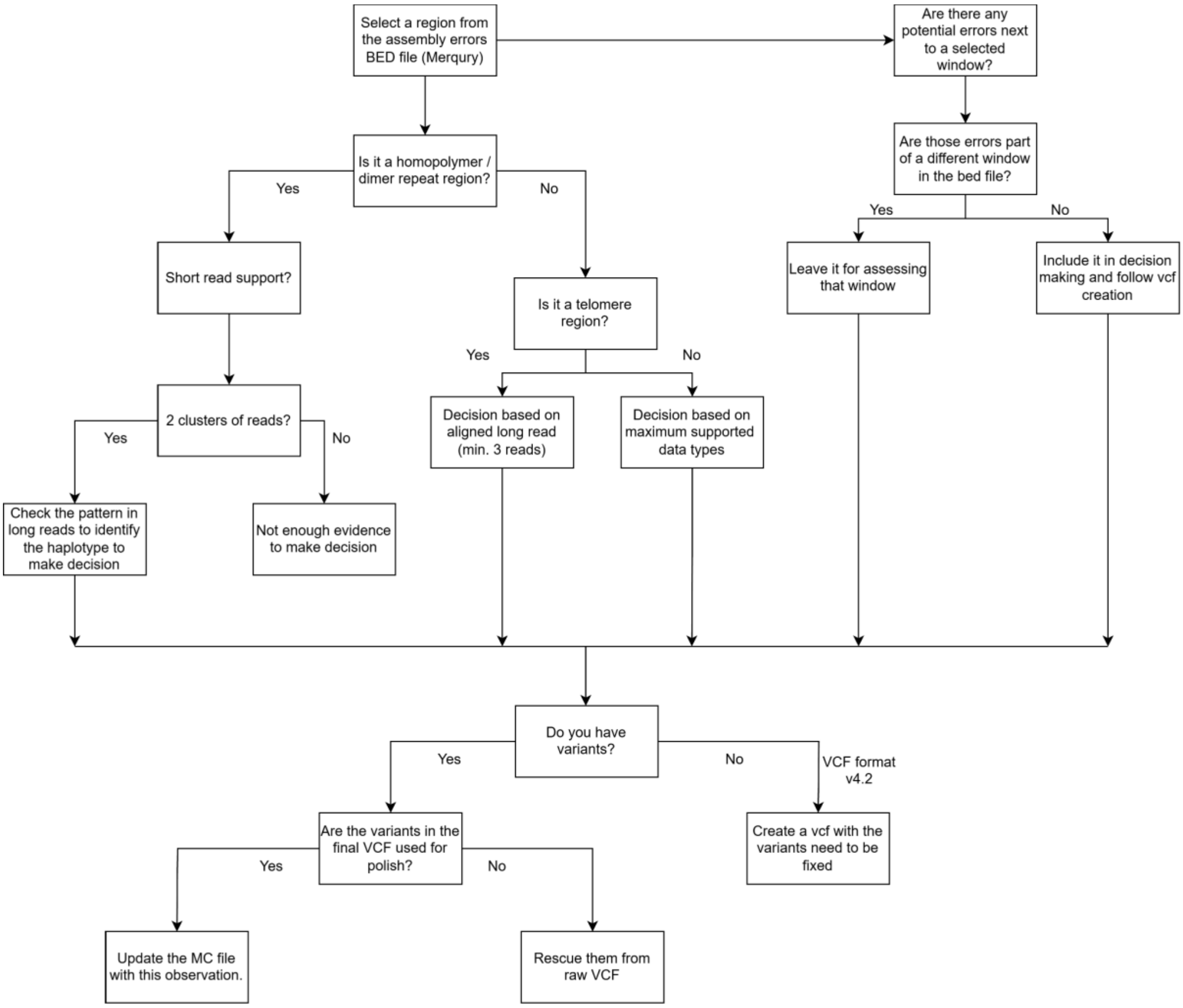


**Figure S4. Overview of manual curation steps for genome polishing.**

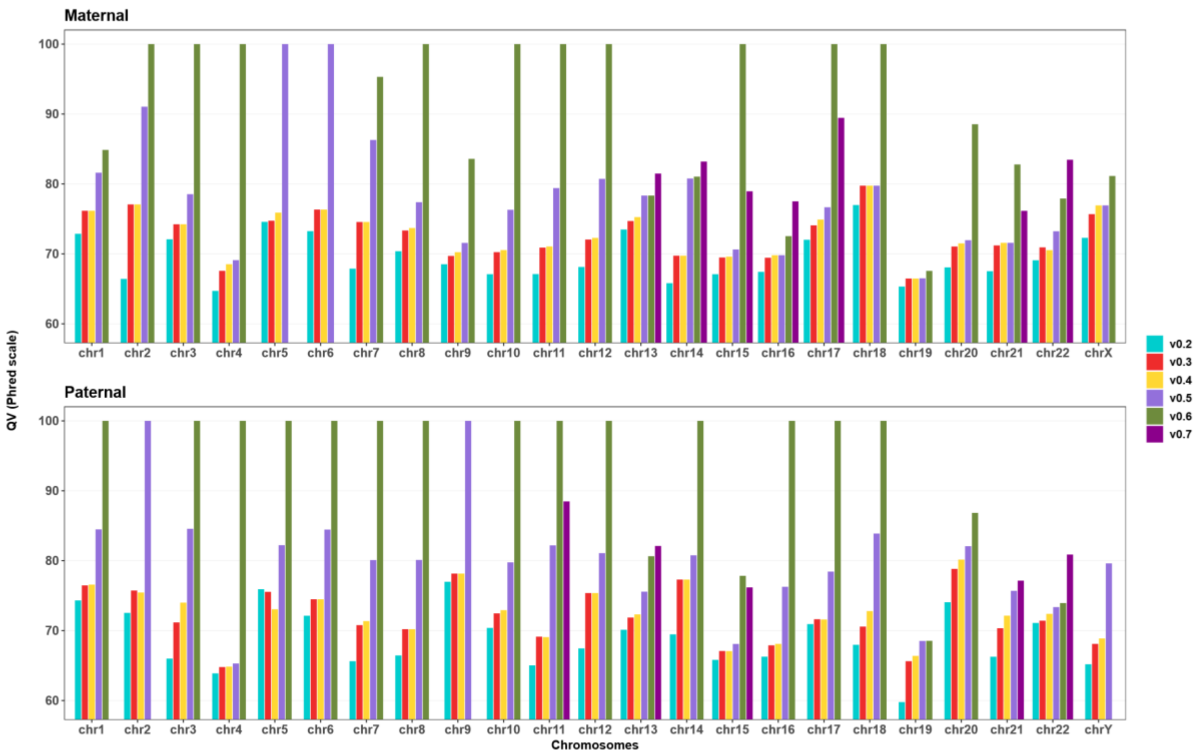


**Figure S5. Comparison of chromosome-level QV estimates across versions of I002C maternal and paternal haplotypes.**

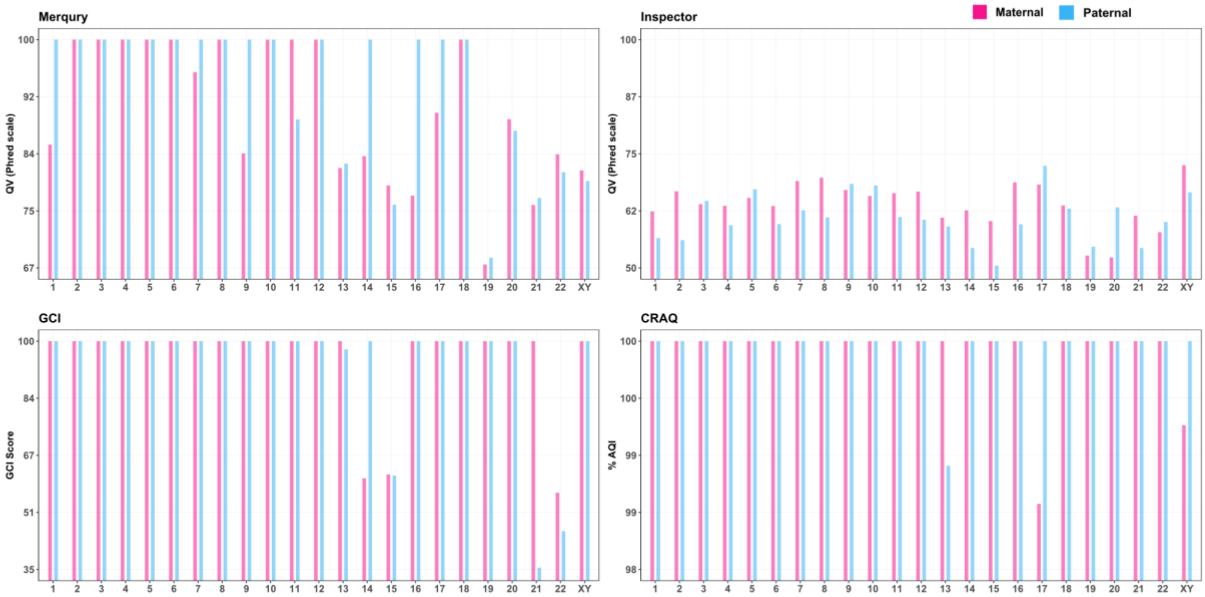


**Figure S6. Chromosome-wise assembly quality evaluation of the I002C genome across multiple tools.** Assembly quality was assessed per chromosome using four independent tools: **Merqury:** QV values are estimated from hybrid k-mers (k=21); lower the assembly specific k-mer multiplicity indicates higher assembly quality. **Inspector**: QV values are calculated from the sum of SNPs and SVs; lower variant counts indicate higher quality. **GCI (G**enome **C**ontiguity **I**ndex): Scores reflect the proportion of high quality read mappings (mapping quality >30, alignment identity (num_match_res/len_aln) >90% and clipped proportion <10%). Lower GCI scores observed in acrocentric chromosomes (13,14,15, 21 and 22) are attributed to rDNA copy numbers and challenges in uniquely mapping reads. **CRAQ: A**ssembly **Q**uality **I**ndex (AQI) scores are based on regional and structural assembly errors identification using alignment Clipping information. Higher AQI indicates fewer errors, with AQI >90 considered reference quality. Across all tools, higher scores indicate improved assembly quality and lower error rates.

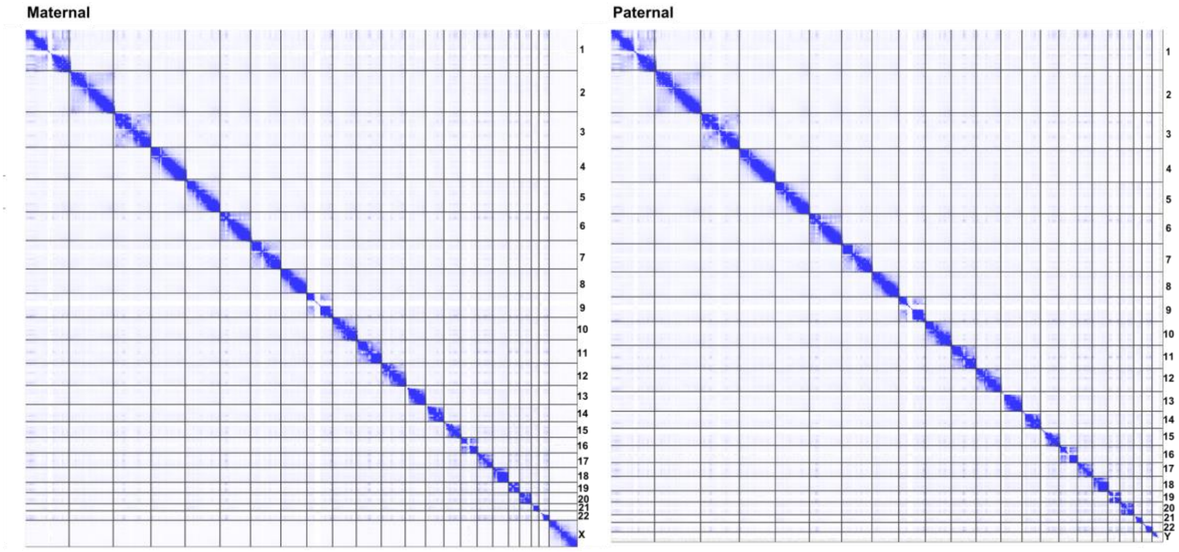


**Figure S7. Omni-C 3D contact matrix visualized using Juicebox to assess chromosome-level accuracy of haplotype-resolved I002C assemblies.**

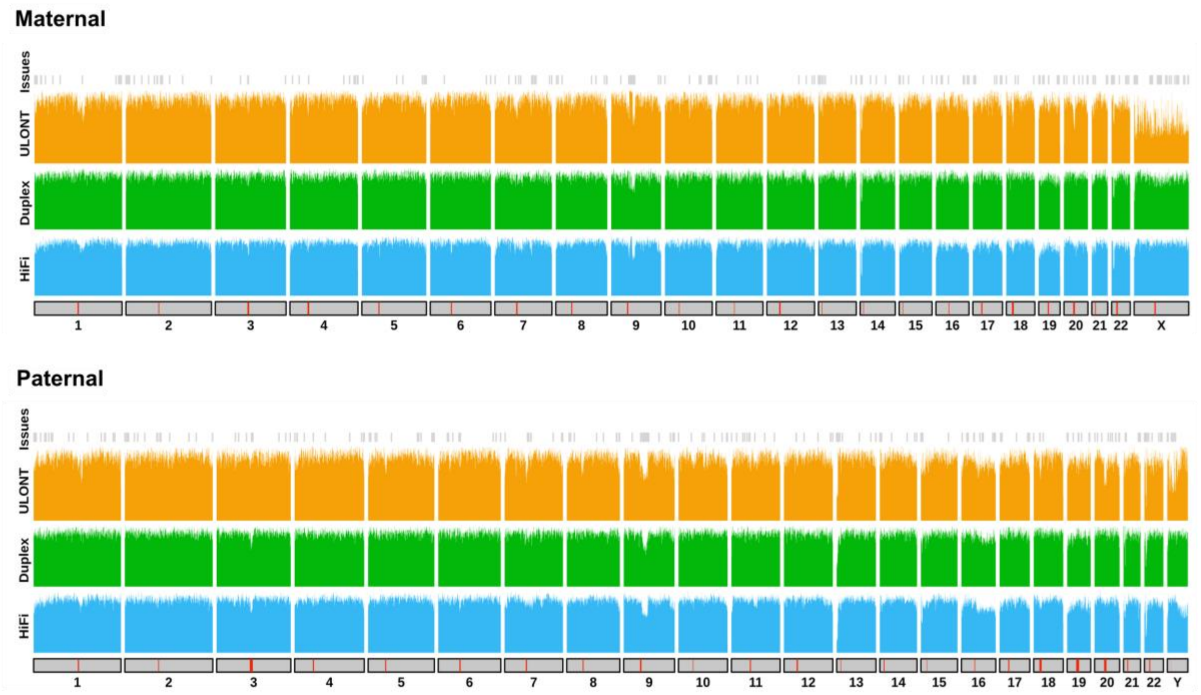


**Figure S8. Sequence coverage-based validation of haplotype-resolved I002C assemblies.** Read coverage across the I002C assemblies using binned PacBio HiFi (blue), ONT Duplex (green), and ONT ultra-long (yellow) reads. For each chromosome, centromeric regions are indicated by red bars. Coverage related issues are labelled and can be further explored by accessing from GitHub repository.

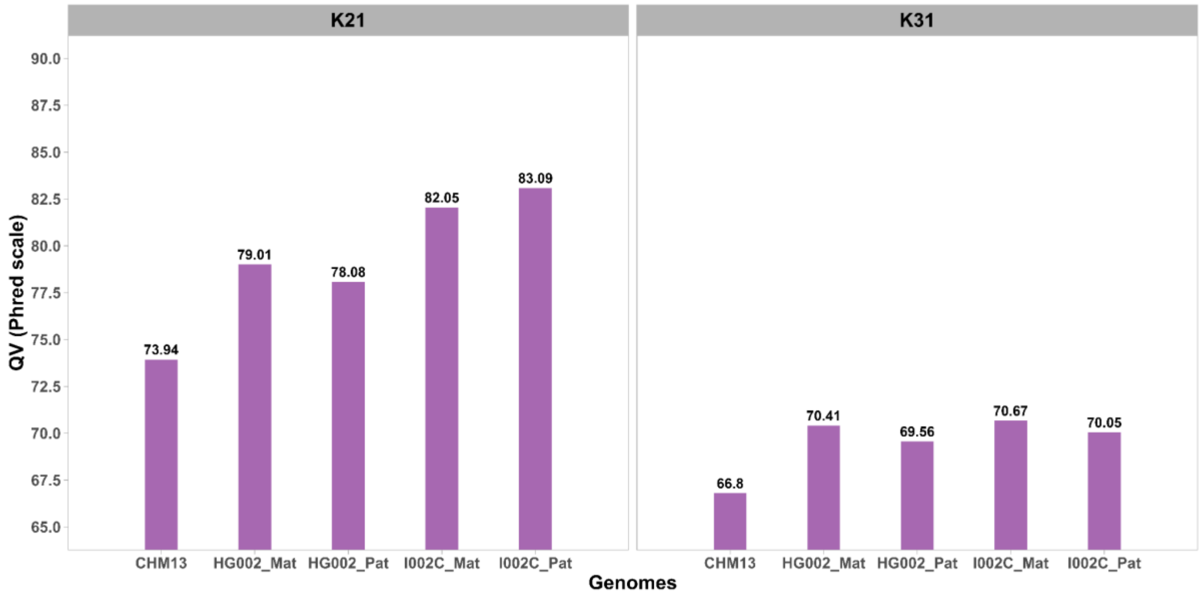


**Figure S9. Merqury quality values (QV) for various genomes using k-mer sizes of 21 and 31.** QV estimates based on 31-mers are generally lower than those from 21-mers due to increased sensitivity in homopolymer-rich regions. CN1 and Yao genomes were excluded from this analysis due to controlled-access restrictions on the sequencing data.

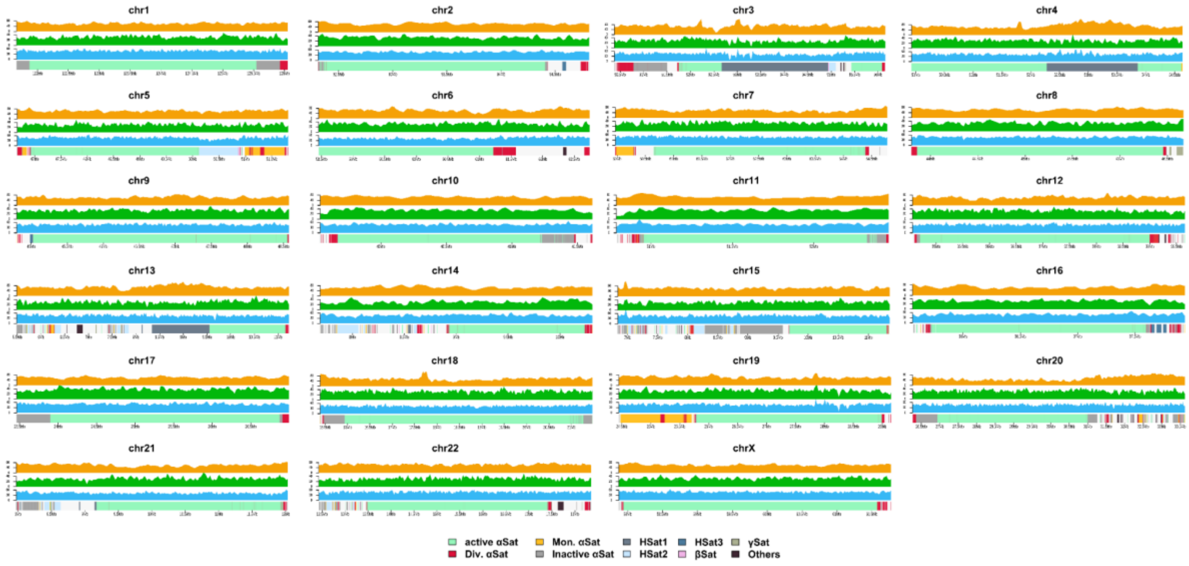


**Figure S10. Read depth profiles across centromeric regions of the I002C maternal haplotype.** Read depth was calculated and visualized across 20 Kbp sliding windows from Winnowmap alignment. For each chromosome, tracks (bottom to top) show centromere satellite annotations followed by read coverage from PacBio HiFi (blue), ONT duplex (green), and ONT ultra-long (yellow) reads.

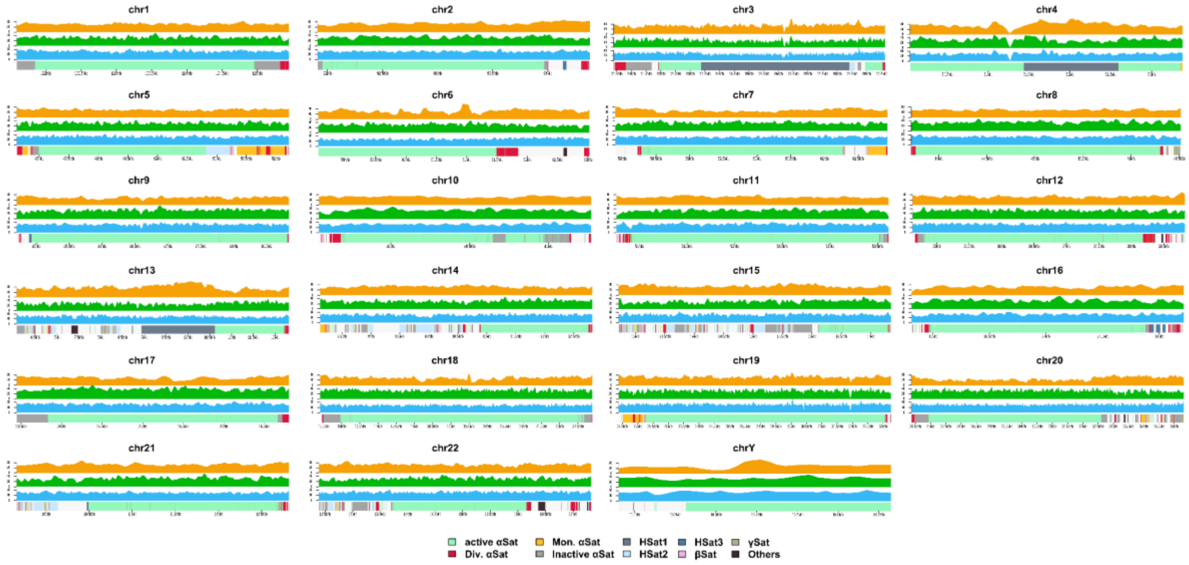


**Figure S11. Read depth profiles across centromeric regions of the I002C paternal haplotype.** Read depth was calculated and visualized across 20 Kbp sliding windows from Winnowmap alignment. For each chromosome, tracks (bottom to top) show centromere satellite annotations followed by read coverage from PacBio HiFi (blue), ONT duplex (green), and ONT ultra-long (yellow) reads.

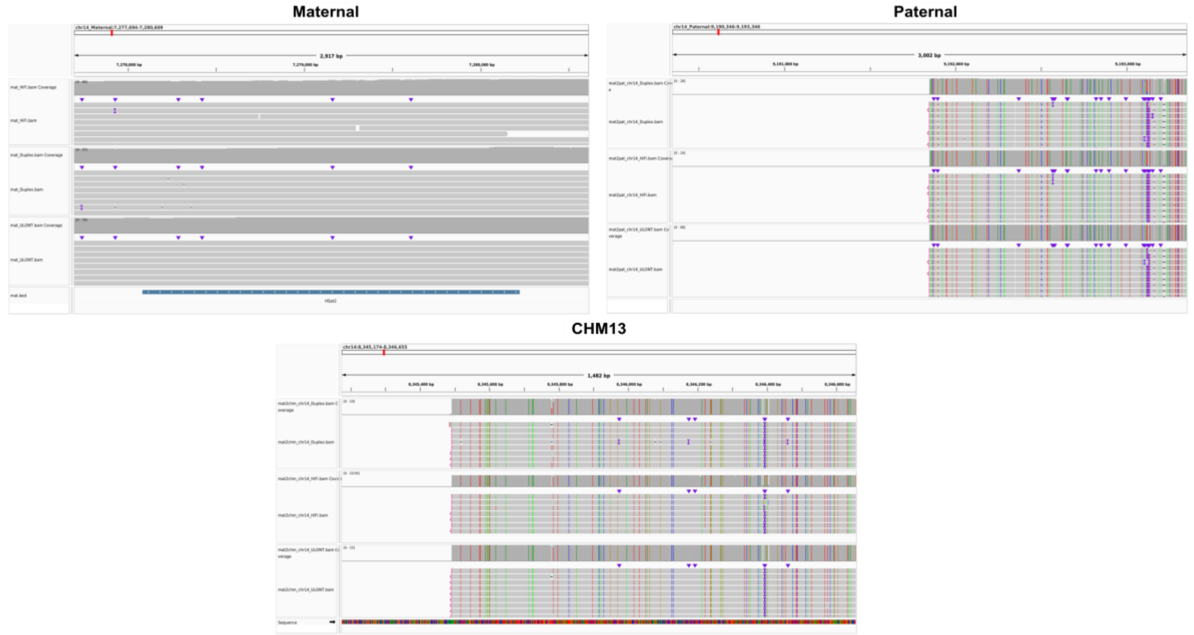


**Figure S12. Read mapping across a ~2 kb maternal-specific HSat2 region on chromosome 14.** An additional ~2 kb HSat2 array present on maternal chromosome 14 is supported by long-read coverage. Binned reads from PacBio HiFi (top track), ONT Duplex (middle), and ONT ultra-long (bottom) validate this region. Reads from this region are mapped to the paternal haplotype or CHM13, the primary alignments show partial clipping reads containing the HSat2 sequence, indicating its absence or divergence in those assemblies.

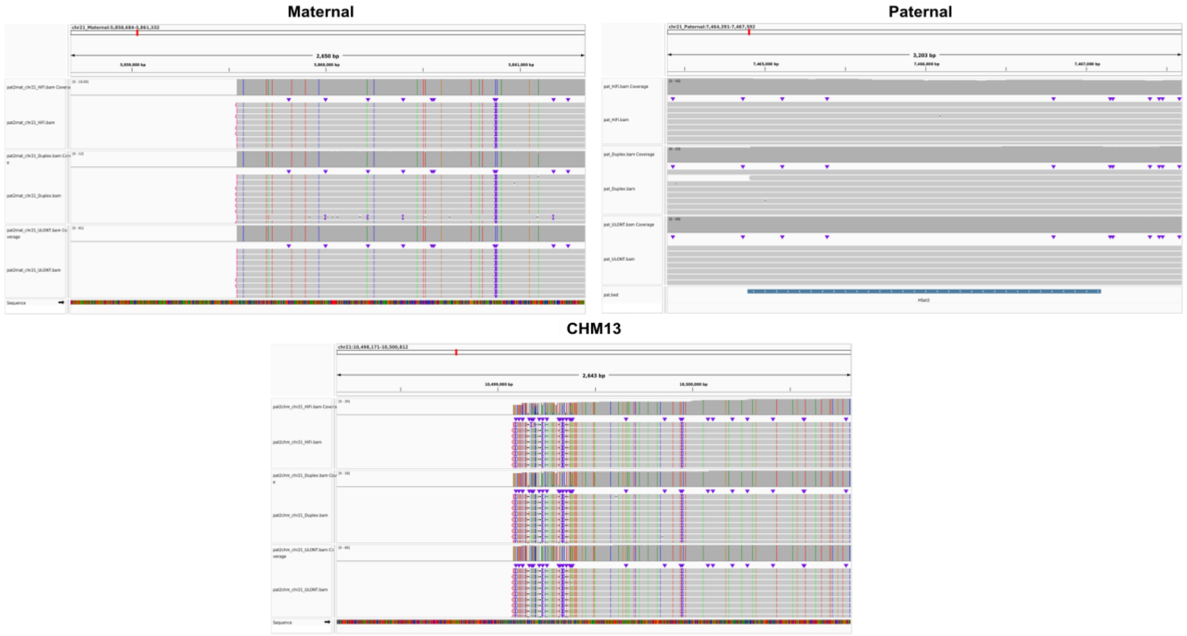


**Figure S13. Read mapping across a ~2 kb paternal-specific HSat2 region on chromosome 21.** An additional ~2 kb HSat2 array present on paternal chromosome 21 is supported by long-read coverage. Binned reads from PacBio HiFi (top track), ONT Duplex (middle), and ONT ultra-long (bottom) validate this region. Reads from this region are mapped to the maternal haplotype or CHM13, the primary alignments show partial clipping reads containing the HSat2 sequence, indicating its absence or divergence in those assemblies.

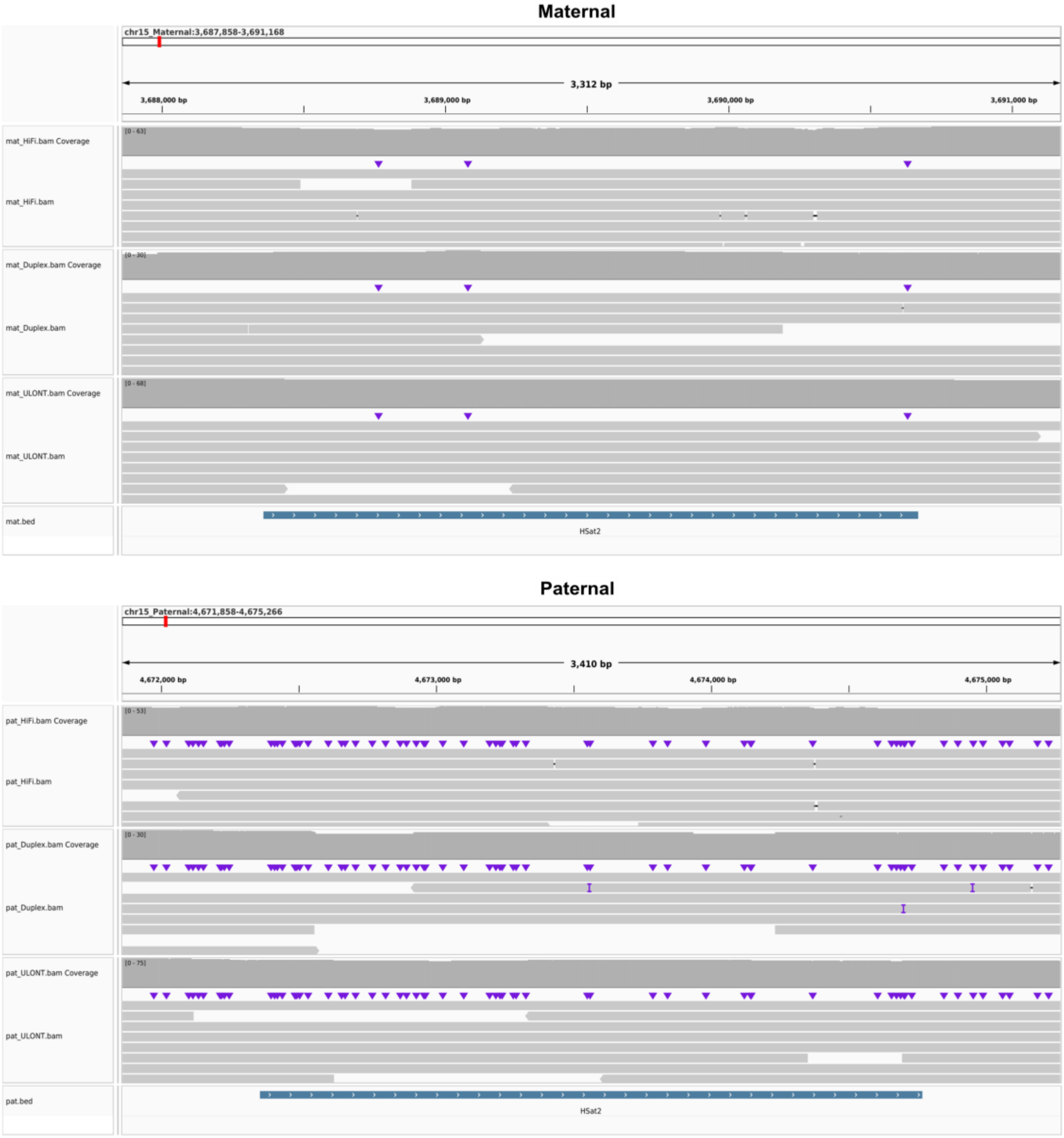


**Figure S14. Read mapping across a ~2 kb HSat2 region on chromosome 15.** An additional ~2 kb HSat2 present on both haplotype chromosome 15 is supported by long-read coverage.

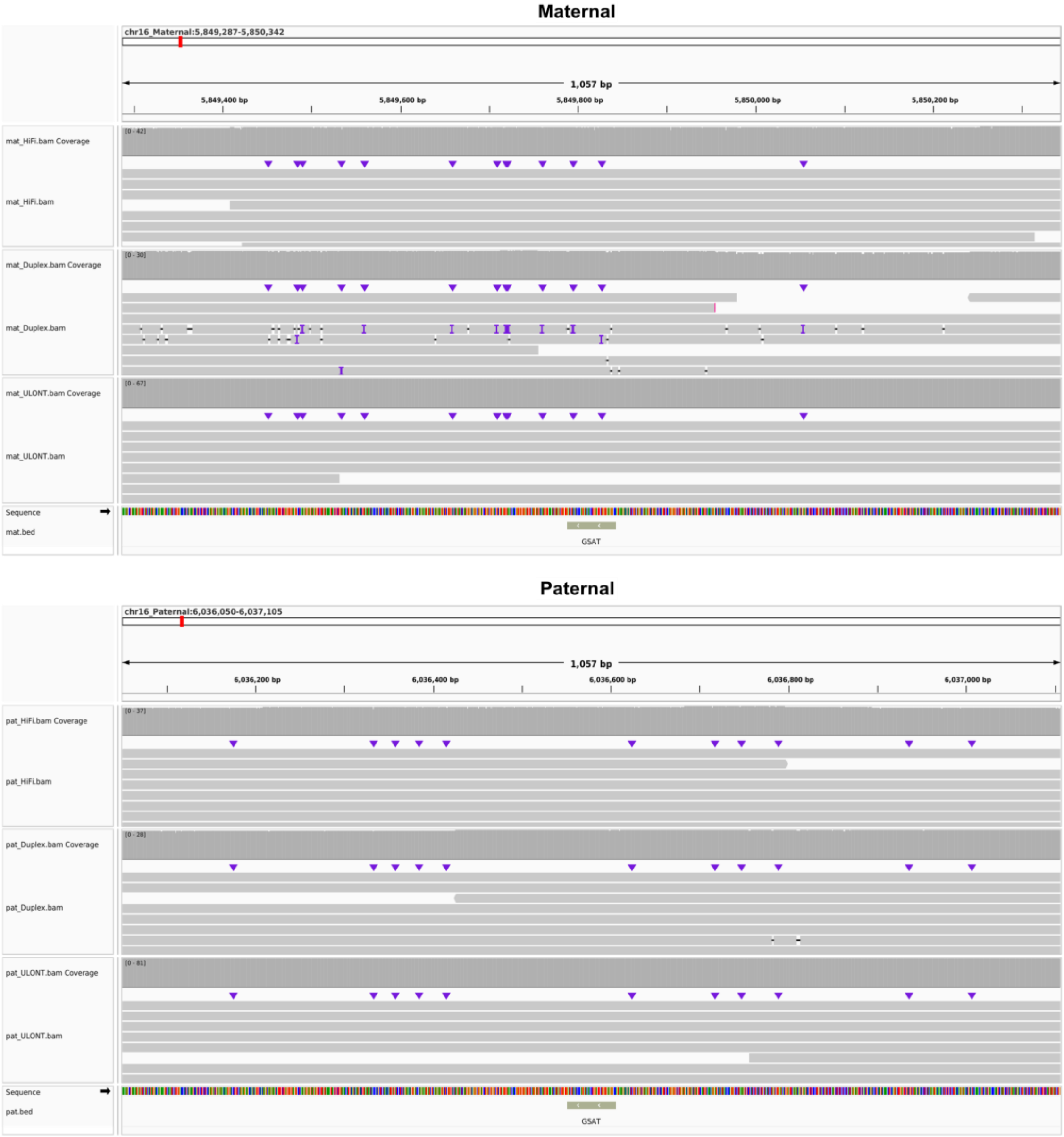


**Figure S15. Read mapping across a 55bp γSat region on chromosome 16.**

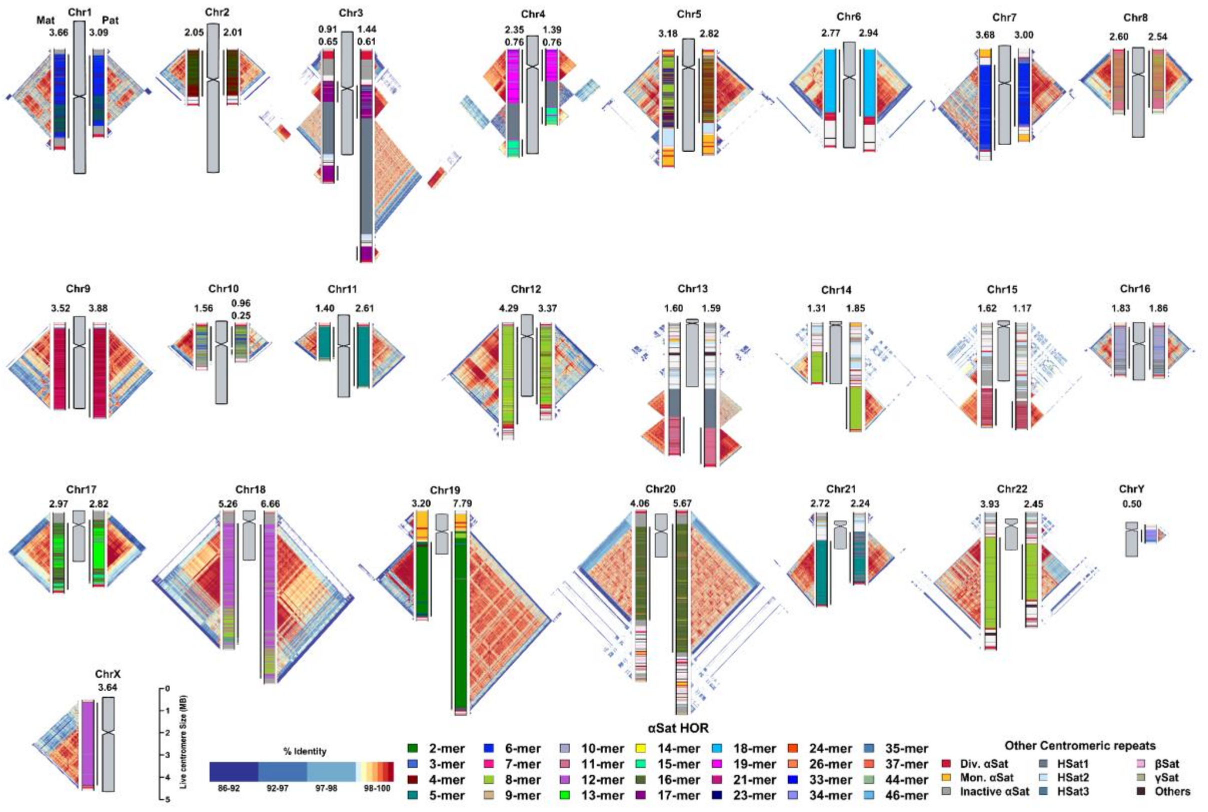

**Figure S16. Overview of centromeric variation between I002C haplotypes.** Centromeric landscapes showing higher-order repeat (HOR) structural variation in pairwise comparisons across chromosomes. For each chromosome, the maternal centromere is shown on the left and the paternal on the right. The length (in Mbp) of the active α-satellite array is indicated at the top and marked with a black line adjacent to each chromosome. Triangular heatmaps representing pairwise sequence similarity were generated using StainGlass.

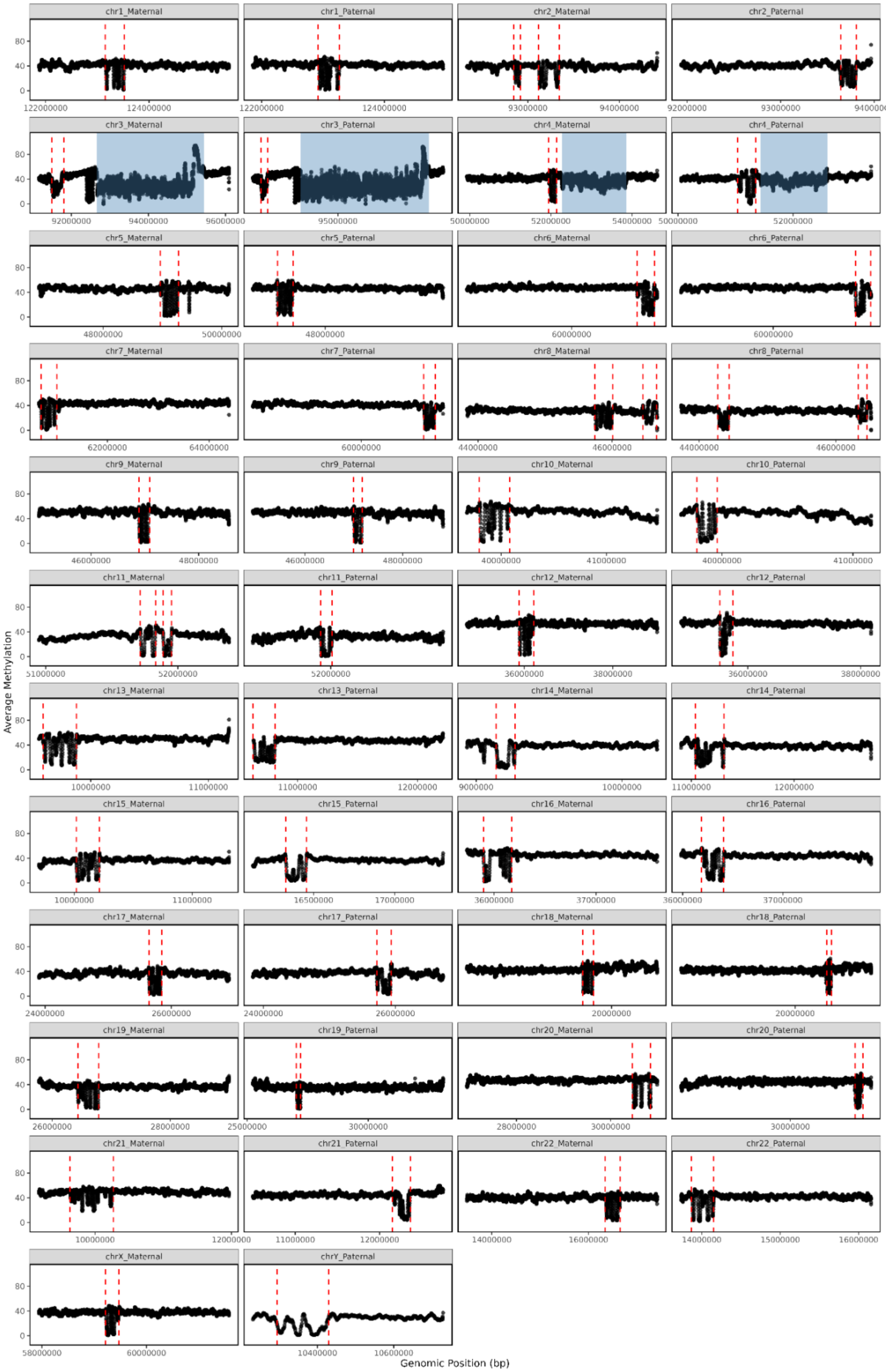

chr1_Maternal
chr1_Paternal
chr2_Maternal
chr2_Paternal
chr3_Maternal
chr3_Paternal
chr4_Maternal
chr4_Paternal
chr5_Maternal
chr5_Paternal
chr6_Maternal
chr6_Paternal
chr7_Maternal
chr7_Paternal
chr8_Maternal
chr8_Paternal
chr9_Maternal
chr9_Paternal
chr10_Maternal
chr10_Paternal
chr11_Maternal
chr11_Paternal
chr12_Maternal
chr12_Paternal
chr13_Maternal
chr13_Paternal
chr14_Maternal
chr14_Paternal
chr15_Maternal
chr15_Paternal
chr16_Maternal
chr16_Paternal
chr17_Maternal
chr17_Paternal
chr18_Maternal
chr18_Paternal
chr19_Maternal
chr19_Paternal
chr20_Maternal
chr20_Paternal
chr21_Maternal
chr21_Paternal
chr22_Maternal
chr22_Paternal
chrX_Maternal
chrY_Paternal
Average Methylation
Genomic Position (bp)

**Figure S17. Centromeric methylation dip regions across all chromosomes in the I002C genomes, separated by haplotype.** Manually annotated centromeric dip regions are indicated by red vertical dashed lines. HSat regions on chromosomes 3 and 4 that are not part of active centromeres are highlighted with a semi-transparent steel blue block for both haplotypes. Average methylation ratios were computed using homozygous and haplotype-specific reads with a sliding window of 10,000 bp and a step size of 1,000 bp.

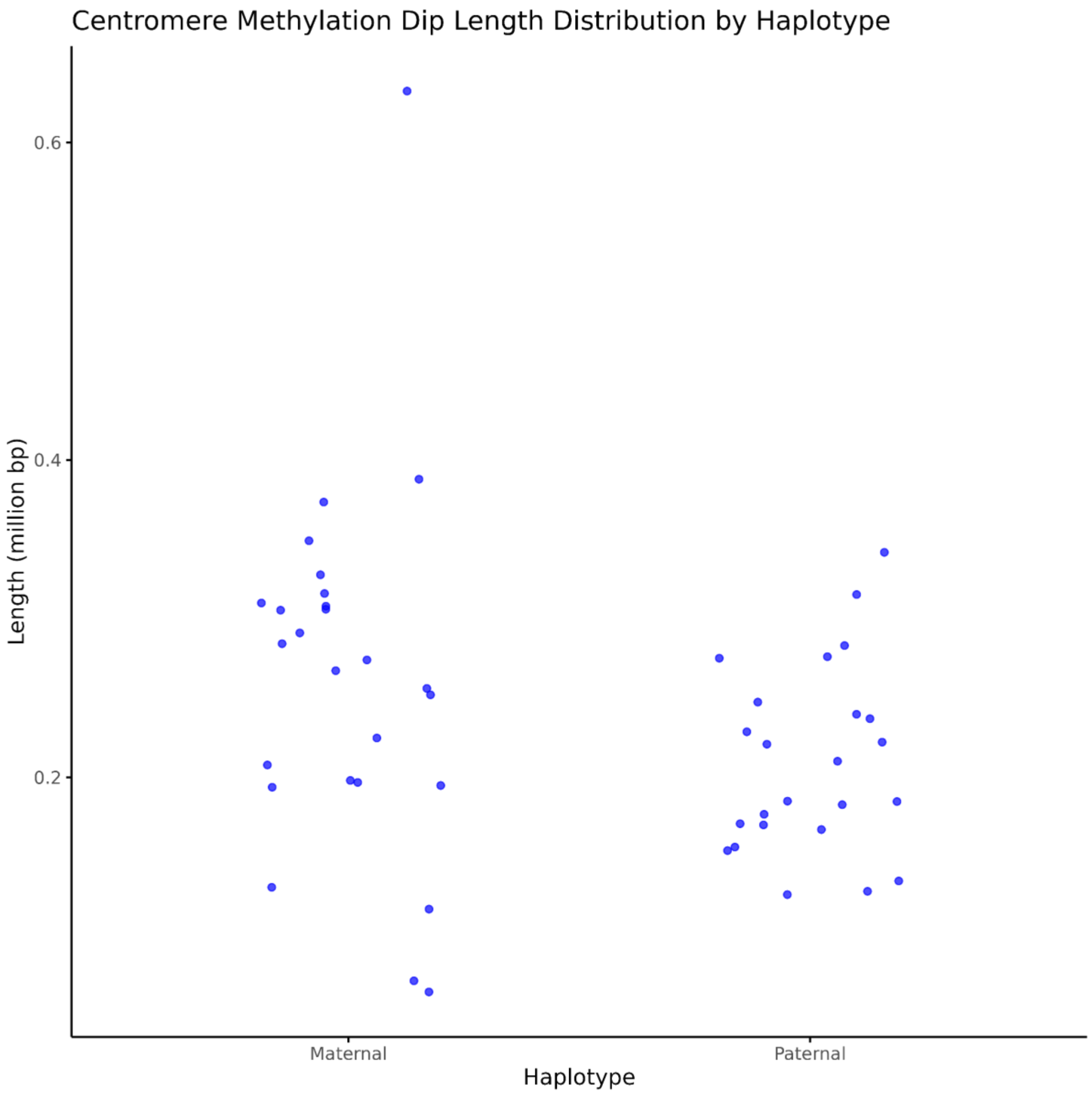


**Figure S18. Length of centromeric methylation dip regions across all chromosomes in the I002C genomes, separated by haplotype.**

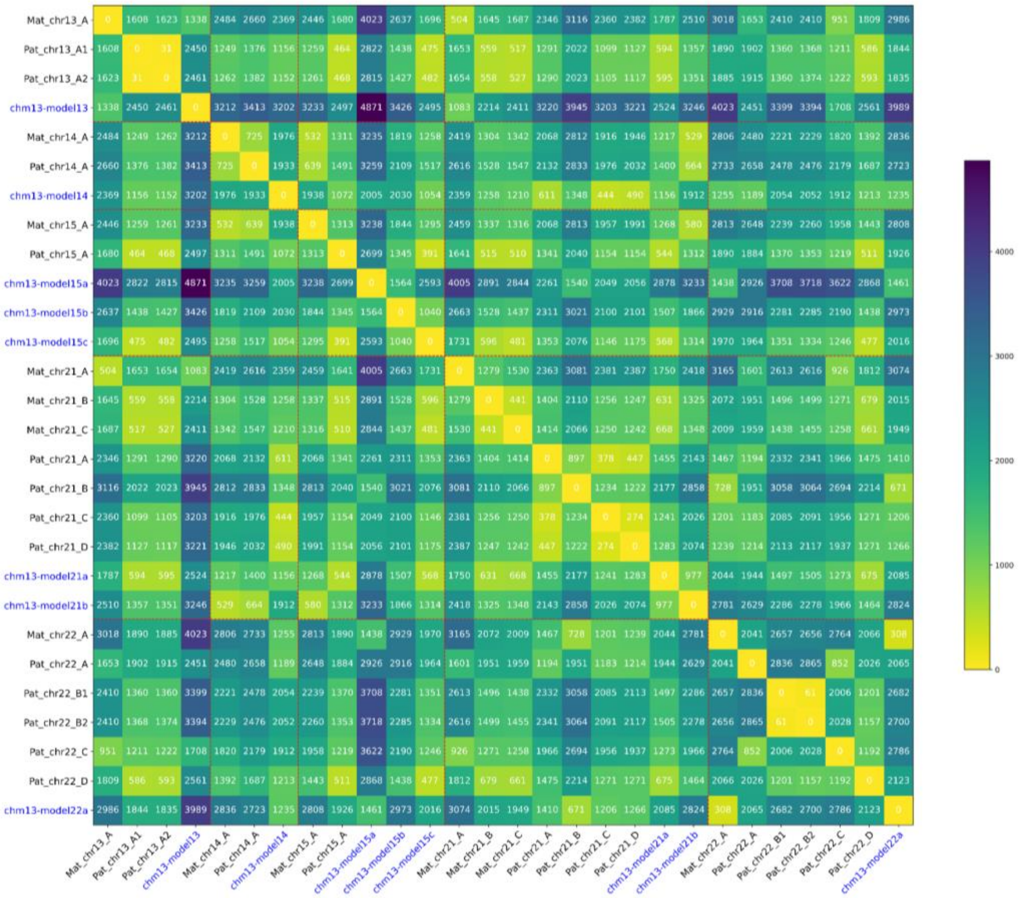


**Figure S19. Pairwise edit distance among I002C and CHM13 rDNA major morphs.** rDNA morphs from CHM13 were labelled blue. Edlib was used to calculate the pairwise edit distance.

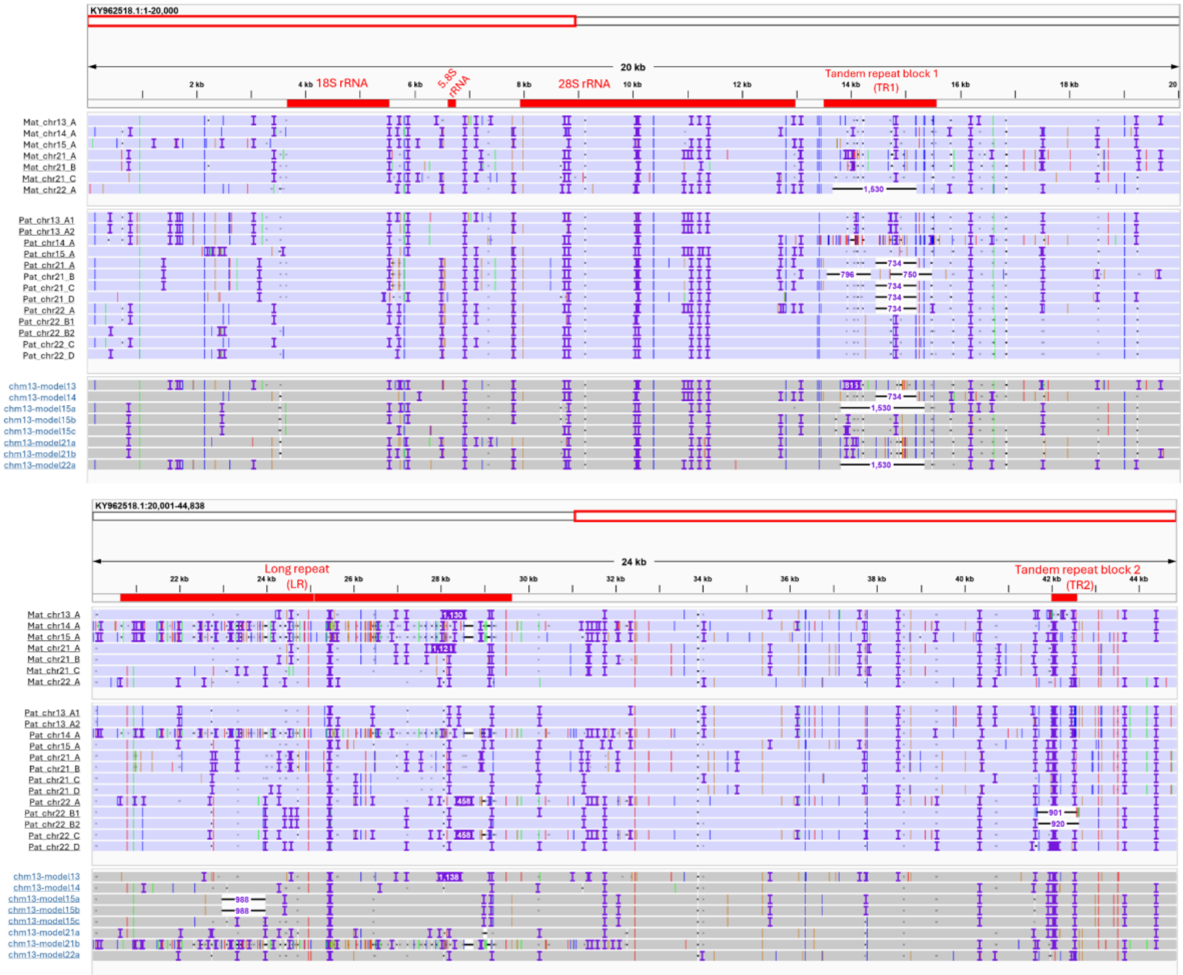


**Figure S20. Visualization of I002C and CHM13 rDNA major morphs.** Morphs were aligned against the canonical rDNA reference KY962518.1 and viewed by IGV. Large indels are enriched in the TR blocks and LR region.

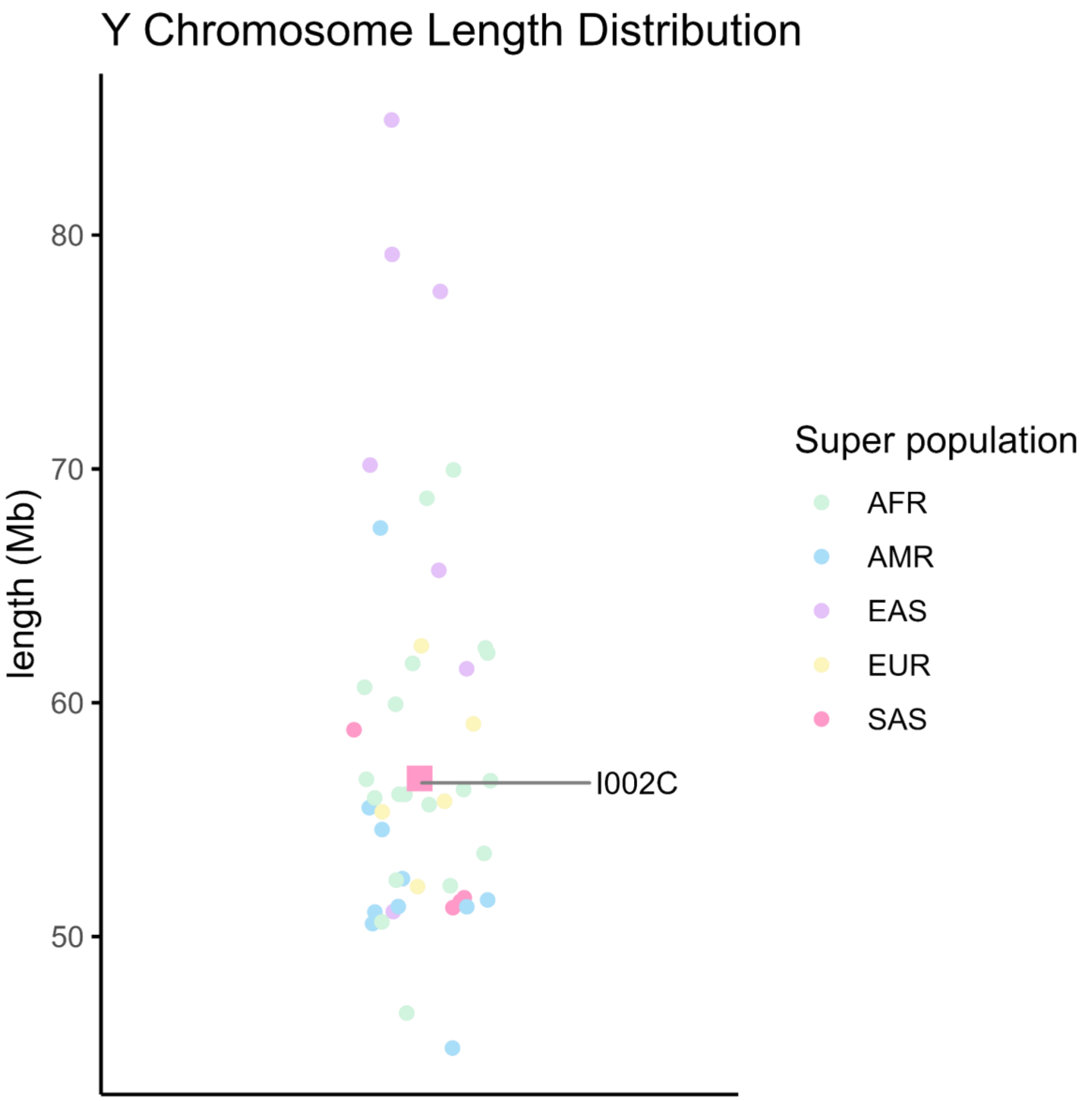


**Figure S21. Comparison of chromosome Y lengths of I002C (red square) and other human genomes.** The I002C chromosome Y length falls within the observed range across these genomes, indicating consistency with known population diversity.

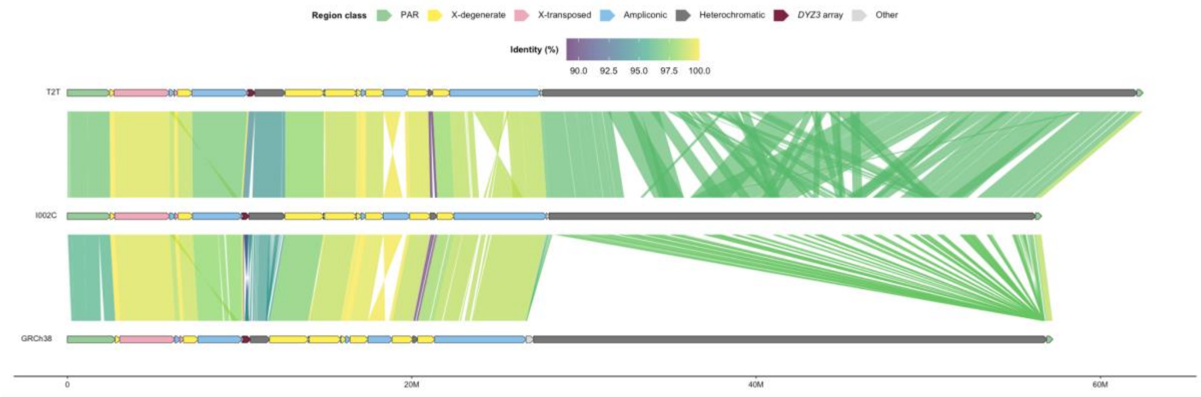


**Figure S22. Alignments of individual chrY subregions between T2T-Y (from CHM13 v2.0), I002C-Y, and GRCh38 (NCBI GCF_000001405.40).** The individual T2T-Y subregions are first mapped to the entire I002C-Y. Then, the individual I002C-Y subregions were mapped to the entire hg38-Y. Colors on the chromosomes Y represent the subregions, whereas colors on the links represent the average alignment identity between the respective subregions.

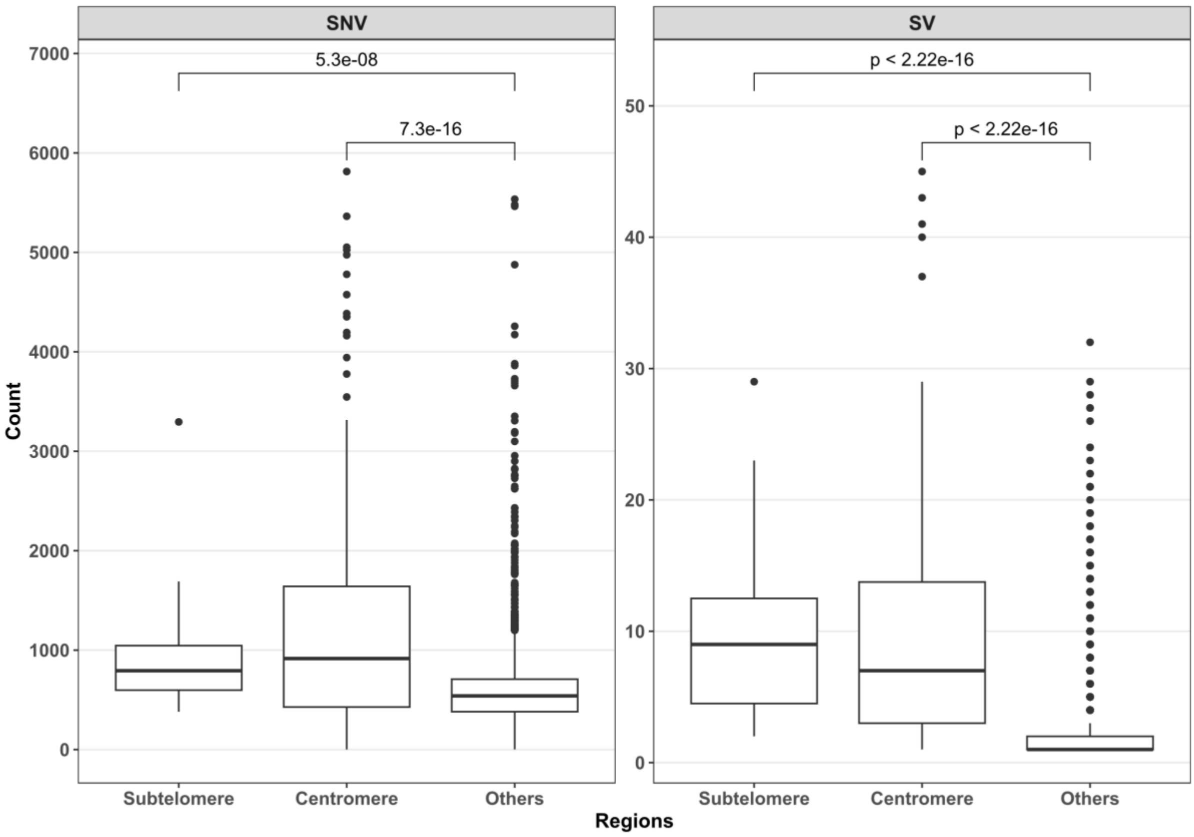


**Figure S23. Heterozygosity comparison across genomic regions in I002C.** Heterozygosity rates using SNVs and SVs counts were compared across sub-telomeric, centromeric, and other genomic regions. Rates were calculated as the number of SNVs or SVs per 500 kb window across the genome.

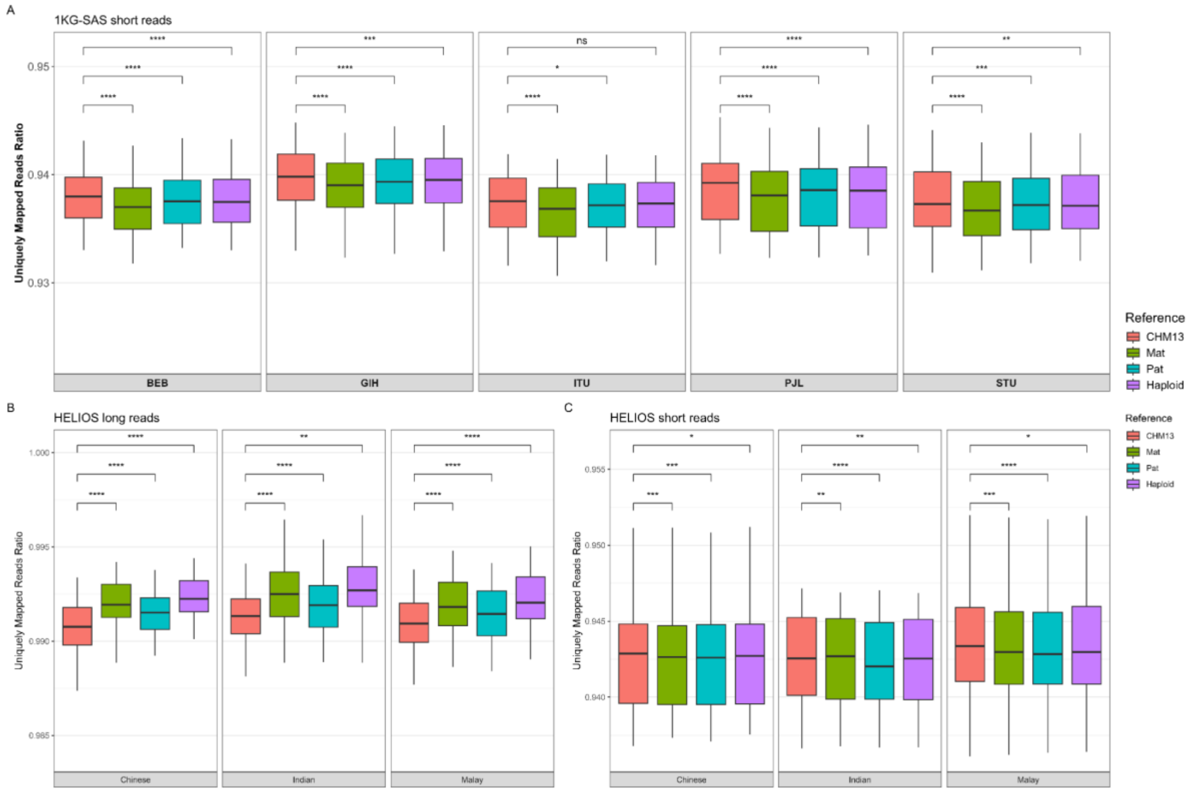


**Figure S24. Distribution of uniquely mapped reads ratio on short-read data (A, C) and long-read data (B).** The mapping ratio among 5 SAS subpopulations and 3 Singapore ethnic groups against four haploid reference genomes: CHM13, I002C Maternal (Mat), I002C Paternal (Pat) and I002C Haploid. The paired Wilcoxon test was used to assess differences in medians between the two groups. Significance levels: ns ($P > 0.05$), * ($P \leq 0.05$), ** ($P \leq 0.01$), *** ($P \leq 0.001$), **** ($P \leq 0.0001$).

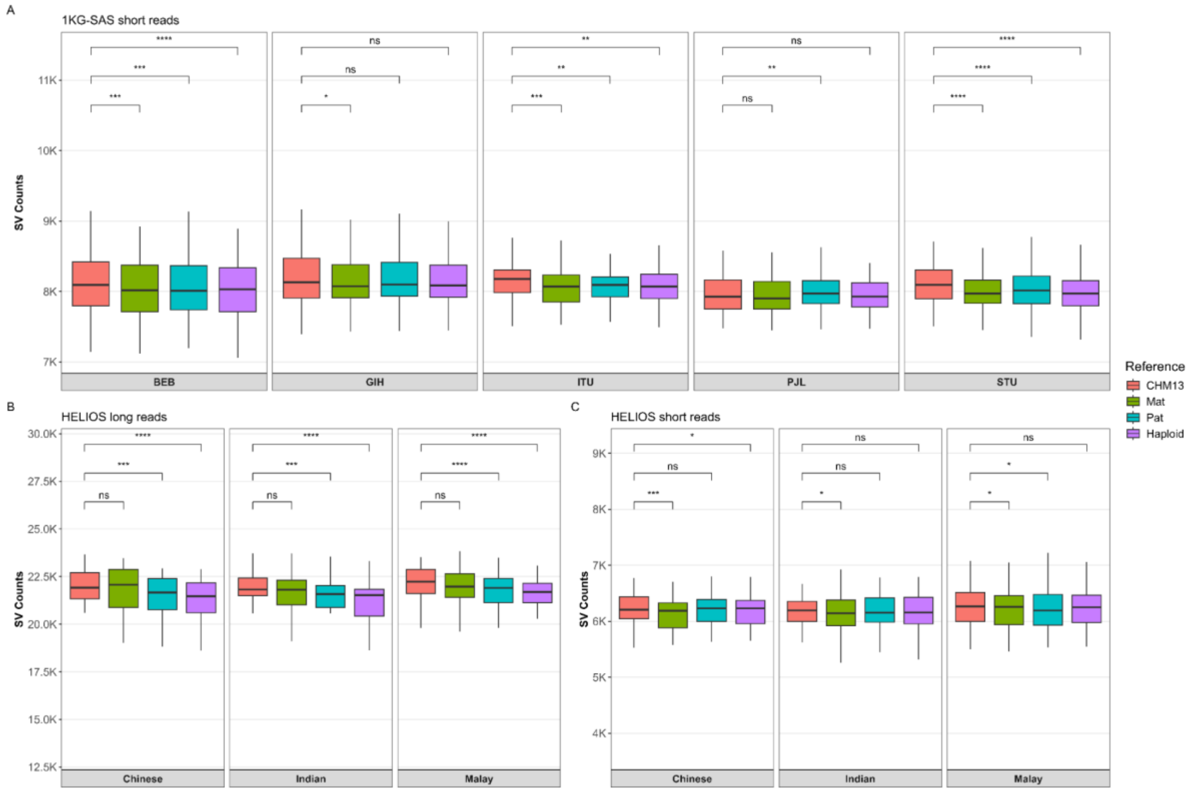


**Figure S25. Counts of SVs based on short-read data (A, C) and long-read data (B).** The comparison was performed among 5 SAS subpopulations and 3 Singapore ethnic groups against four haploid reference genomes: CHM13, I002C Maternal (Mat), I002C Paternal (Pat) and I002C Haploid. The paired Wilcoxon test was used to assess differences in medians between the two groups. Significance levels: ns (P > 0.05), * (P ≤ 0.05), ** (P ≤ 0.01), *** (P ≤ 0.001), **** (P ≤ 0.0001).

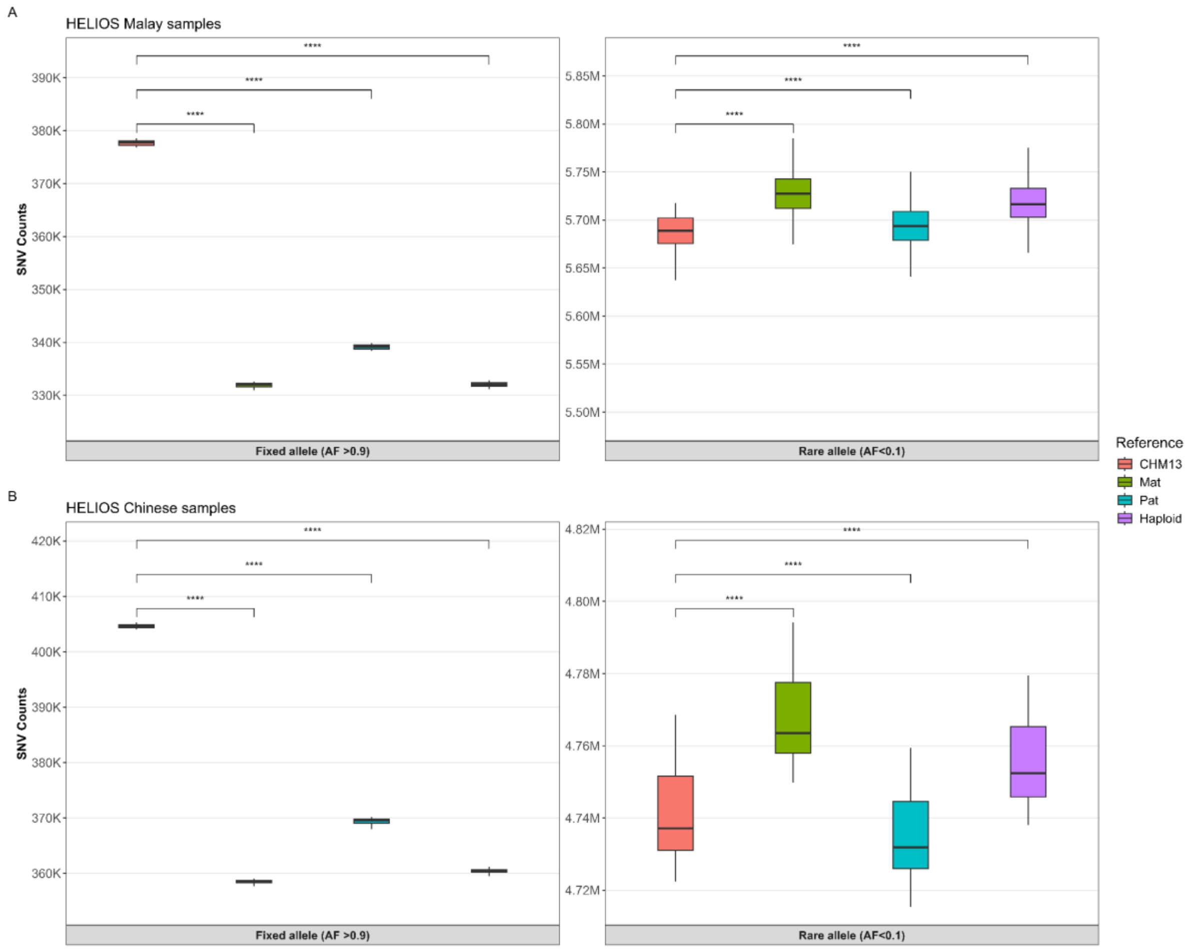


**Figure S26. Allele frequency spectrum of fixed or nearly fixed variants (AF >0.9) and rare variants (AF < 0.1).** A) Singapore Malay samples B) Singapore Chinese samples. The comparison of variants were performed against four references. The paired Wilcoxon test was used to assess differences in medians between the two groups. Significance levels: ns (P > 0.05), * (P ≤ 0.05), ** (P ≤ 0.01), *** (P ≤ 0.001), **** (P ≤ 0.0001)

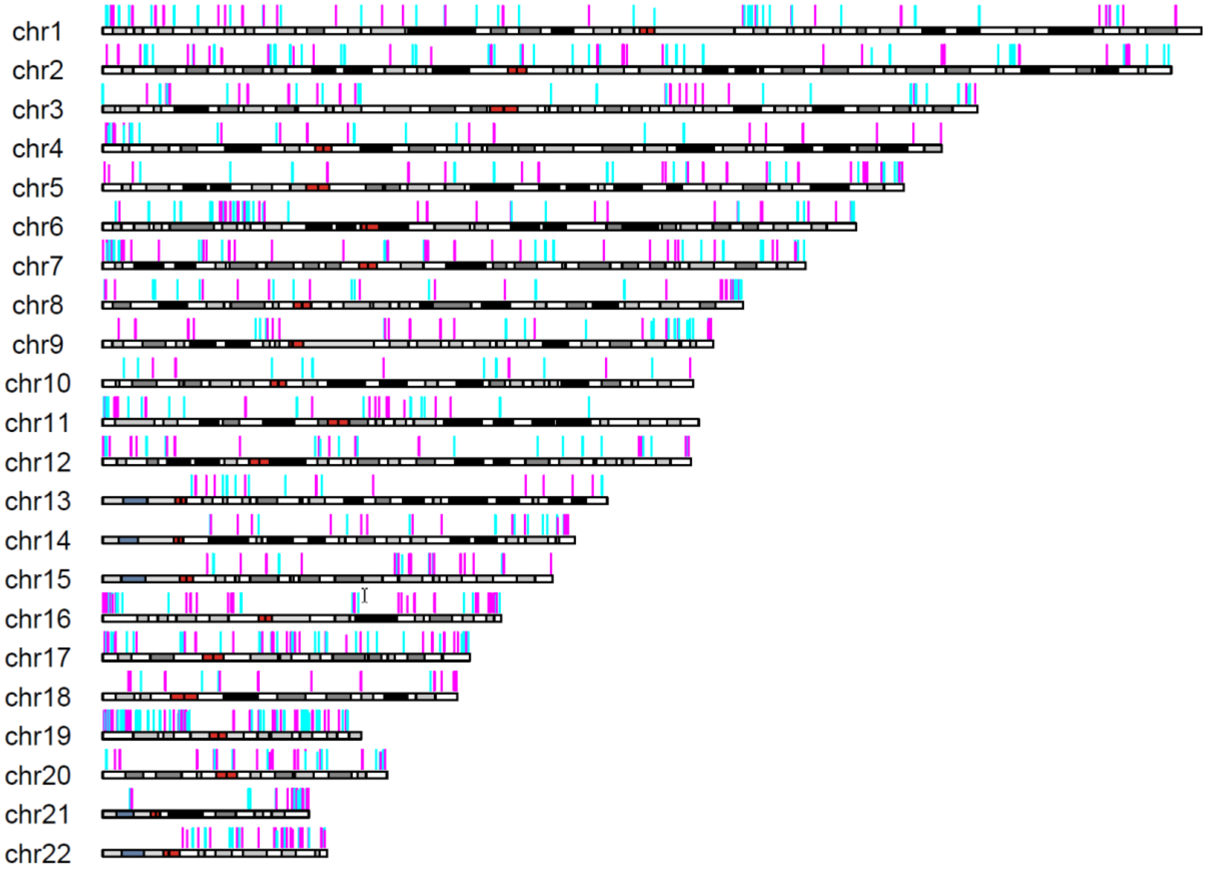


**Figure S27. Genomic distribution of differentially methylated regions (DMRs) of maternal (magenta) and paternal (cyan) origin across GRCh38 autosomal chromosomes.** DMRs were defined as clusters of at least five single-CpG differentially methylated loci (DMLs), each loci with ≥97.5% methylation difference between the two haplotypes. A total of 968 DMRs were identified.

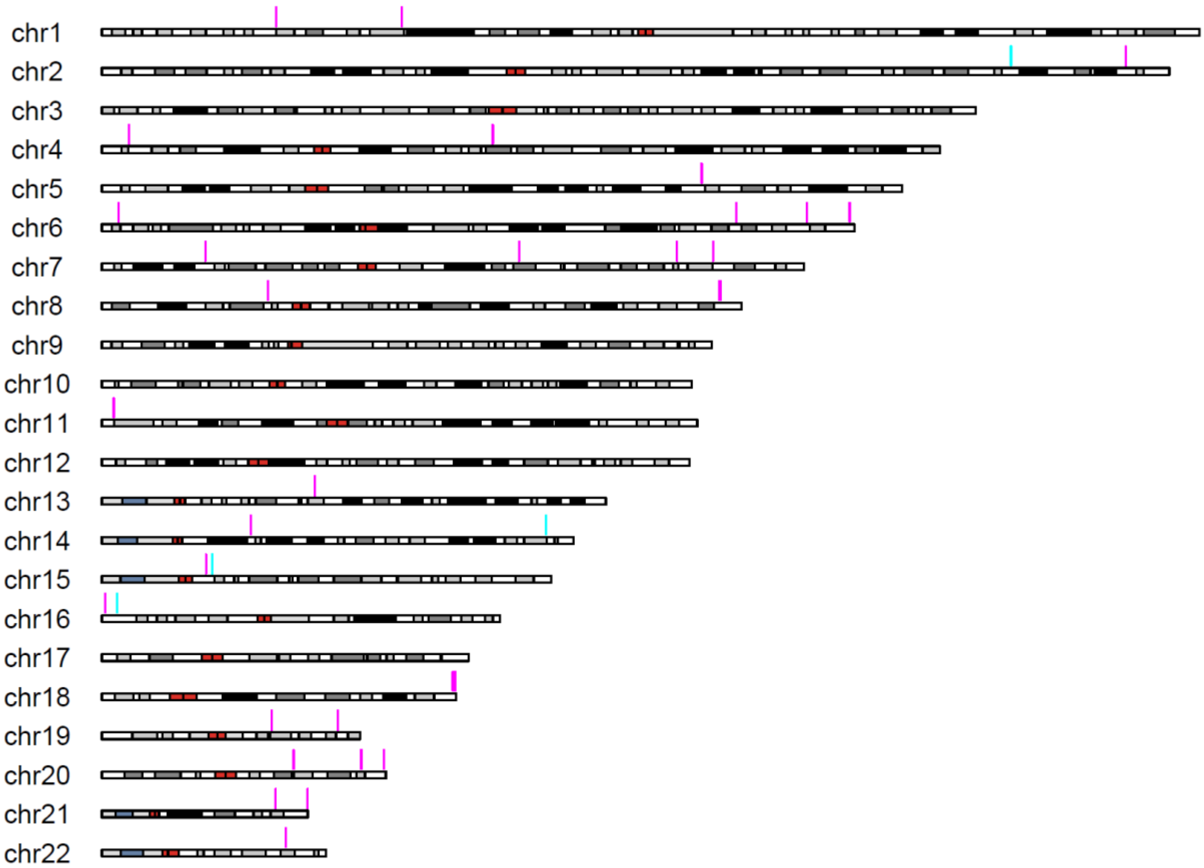


**Figure S28. Genomic distribution of 46 differentially methylated regions (DMRs) of maternal (magenta) and paternal (cyan) origin across GRCh38 autosomal chromosomes, previously reported by at least two independent studies.** DMRs were defined as clusters of at least five single-CpG differentially methylated loci (DMLs), each showing a methylation difference of ≥97.5% between the two haplotypes. All 46 regions were confirmed to have the same methylated allele as reported in previous studies.

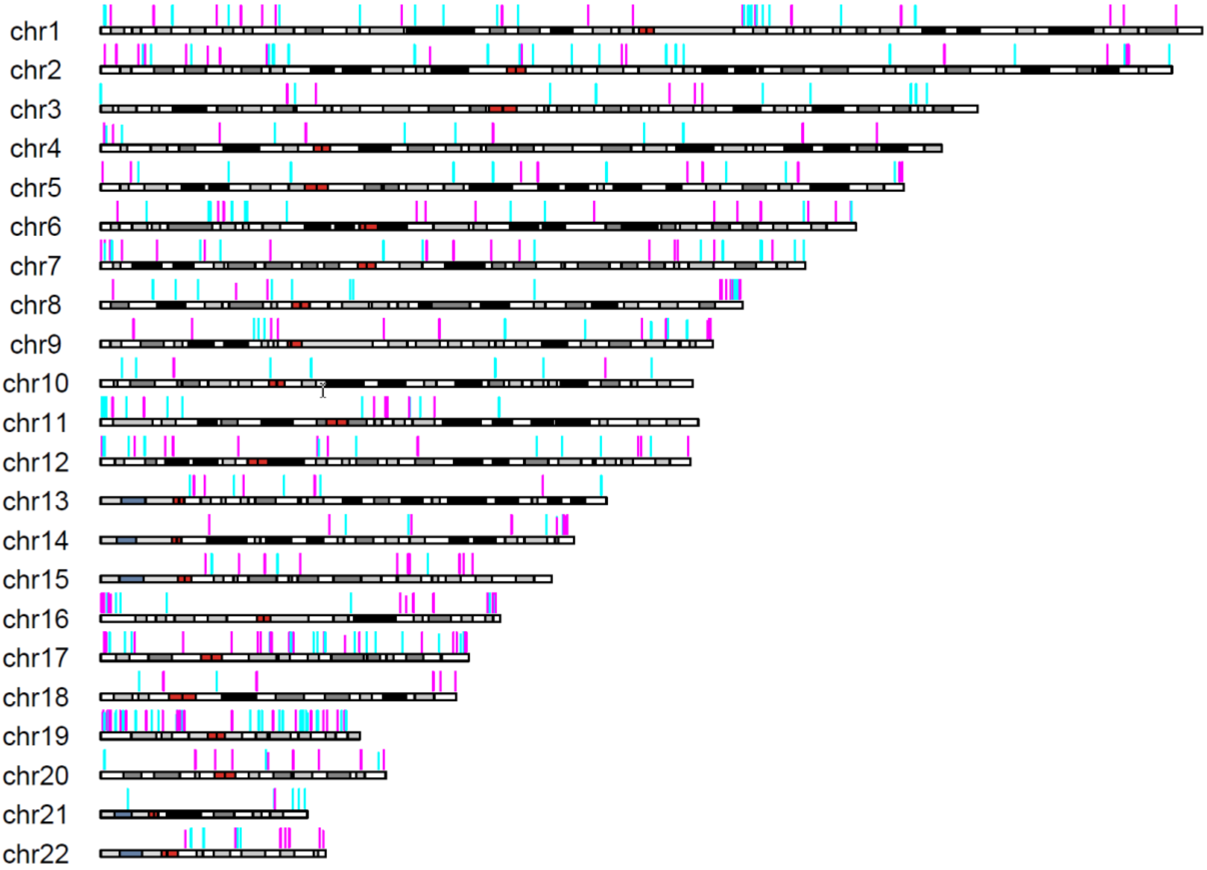


**Figure S29. Genomic distribution of subset of maternally (magenta) and paternally (cyan) derived differentially methylated regions (DMRs) intersecting core promoter or first exon regions of experimentally verified transcripts across GRCh38 chromosomes.** In total, 2,229 such transcripts were identified, corresponding to 596 distinct genes. DMRs were defined as clusters of at least five single-CpG differentially methylated loci (DMLs), each exhibiting a methylation difference of ≥97.5% between the two haplotypes. Core promoters were defined as regions spanning 300 bp upstream to 100 bp downstream of the TSS, taking transcript strand orientation into account.

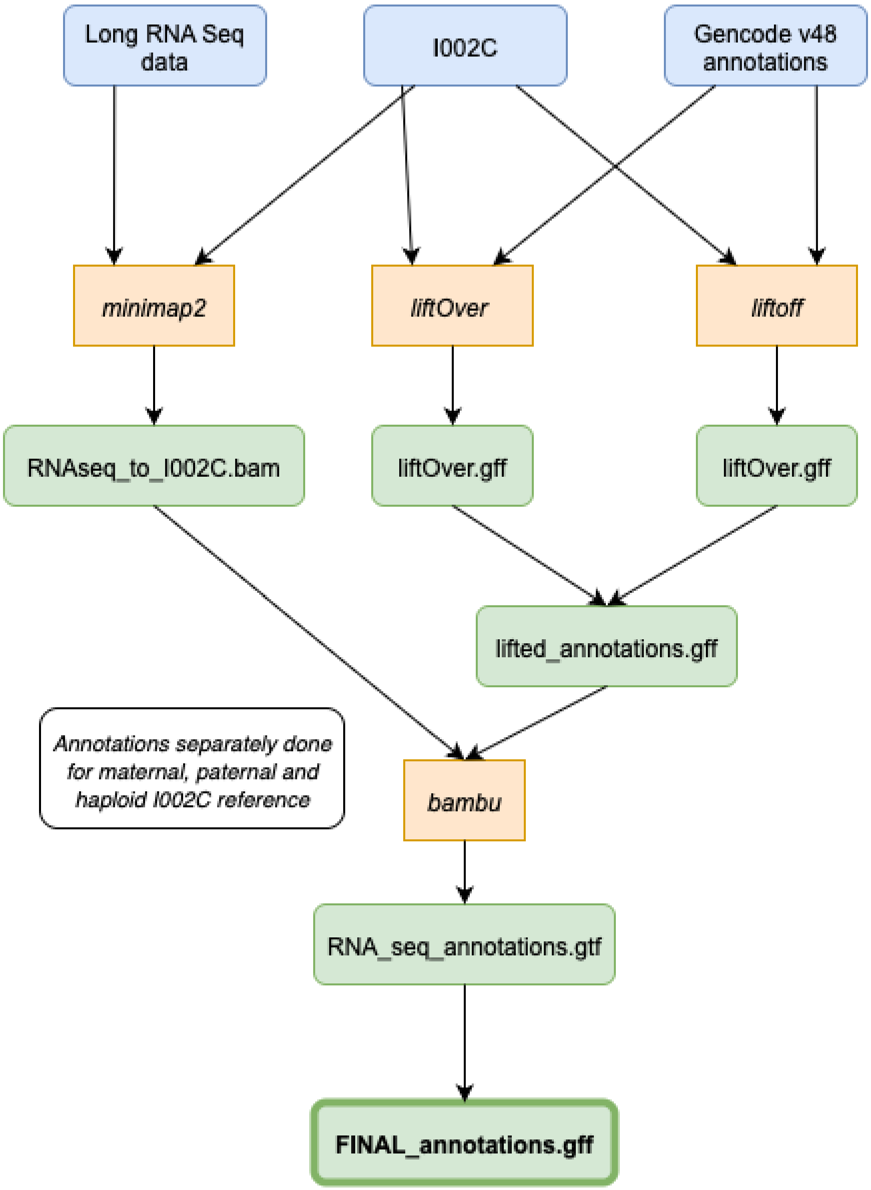


**Figure S30. Gene annotation pipeline.** The pipeline includes lifting Gencode v48 model annotations using liftoff and liftOver and combination with long RNA Seq data annotations using bambu.
