## Additional methods for "A Complete Telomere-to-Telomere Diploid Reference Genome for South Asian Population"

### Read binning

#### ONT ultra-long

Ultra-long read binning was performed using the canu (v2.4)[103] “haplotype” mode with parental-specific k-mers derived from PacBio Revio sequenced HiFi read data. We performed binning of ultra-long ONT reads longer than 50 kbp. Of the 4,449,721 reads processed, 2,189,550 were assigned to the maternal haplotype and 2,248,568 to the paternal; 11,603 reads remained unclassified and were excluded from downstream analyses.

command:

```
canu -p asm -d read_binning_out genomeSize=3g useGrid=false merylMemory=300 merylThreads=64 hapthreads=64 hapMemory=300 -haplotypePat pat_R*.fq.gz -haplotypeMat mat_R*.fq.gz -nanopore-raw UL_all_reads.fq.gz -stopAfter=haplotype
```

#### ONT Duplex

Duplex reads, longer than 10 kbp, were binned using the same approach as for the ultra-long ONT reads, applying the identical canu command described above. From 5,325,183 reads processed, 2,309,872 were assigned to the maternal haplotype and 2,236,549 to the paternal; 778,762 reads remained unbinned and were excluded from further analysis.

#### PacBio HiFi

Considering the shorter read length of PacBio HiFi reads- on average at least twice shorter than ONT duplex and approximately five times shorter than ONT ultra-long reads, we implemented a combined binning approach using *yak* (v0.1-r66-dirty https://github.com/lh3/yak) and a canu-derived trio-binning tool (v1.0.0. https://github.com/esrice/trio_binning/releases/tag/v1.0.0). Both approaches used parental k-mers from short reads. Final haplotype assignments were based on the intersection of classifications from both tools. Reads with conflicting or ambiguous classifications were assigned using a custom scoring formula. Thresholds in the formula were determined from the distributions of k-mer count sums and differences, and the third quartile was used as the cutoff in each case (1–4).

commands:

```
case 1) yak=no_count, tb=mat/pat:
      if #tb_mat + #tb_pat < threshold1 : label = yak classification
      elif abs(#tb_mat - #tb_pat) > threshold2: label = tb classification
      else discard
case 2.1) yak=ambigous and tb=mat/pat
      if abs(#tb_mat - #tb_pat) > threshold2 and #tb_mat + #tb_pat > threshold1: label = tb classification
      else discard
case 2.2) yak = a and tb=ambigous
      discard

case 3) yak=mat/pat and tb=ambigous
      if #yak_mat + #yak_pat < threshold3: label = tb classification
      elif abs(#yak_mat - #yak_pat) > threshold4: label = yak classification
      else discard

case 4) yak=mat/pat and tb=pat/mat
      discard
```

Of the 22,266,938 reads processed, 8,290,553 (37,23%) were assigned to the maternal haplotype and 7,856,353 (35,28%) to the paternal. An additional 5,764,696 (25,89%) reads lacking haplotype-specific k-mers were included in both haplotypes, while 355,336 reads (1,6%) were discarded.

commands:

```
yak count -b37 -t16 -o pat_illumina.yak <(cat pat_R1.fq.gz pat_R2.fq.gz) <(cat pat_R1.fq.gz pat_R2.fq.gz)
```

```
yak count -b37 -t16 -o mat_illumina.yak <(cat mat_R1.fq.gz
mat_R2.fq.gz) <(cat mat_R1.fq.gz mat_R2.fq.gz)

yak triobin -c1 -d 5 pat_illumina.yak mat_illumina.yak
pb_hifi_unbinned.fq.gz > yak_pb_hifi.txt

find-unique-kmers -k 21 -p 8 mat_R1.fq.gz,mat_R2.fq.gz \
 pat_R1.fq.gz,pat_R2.fq.gz

classify-by-kmers pb_hifi_unbinned.fq.gz pat_only_kmers.txt
mat_only_kmers.txt --haplotype-a-out-prefix pat.all --haplotype-b-
out-prefix mat.all --unclassified-out-prefix unclass.all >
trio_binn.out.txt
```

##### Omni-C binning

Omni-C reads were binned using the meryl (v1.4.1) tool from the Merqury[27] using parental-specific k-mers. Reads were assigned to a haplotype if they did not contain any k-mers from another haplotype.

command:

```
meryl-lookup -exclude -sequence OmniC_R1.fq.gz OmniC_R2.fq.gz -
output mat_R1.fq.gz mat_R2.fq.gz -mers pat_kmers.meryl
```

#### ONT ultra-long data correction

Binned ultra-long reads were first pre-processed with Porechop (v0.2.4, https://github.com/rrwick/Porechop) and duplex_tools (v0.3.3, https://github.com/nanoporetech/duplex-tools) to remove adapter sequences. All-vs-all alignments were then performed using Minimap2 (v2.26-r1175) to identify overlaps among reads. The read correction was carried out using the developmental version of HERRO by applying the R10 correction model.

commands:

```
minimap2 -K8g -cx ava-ont -k25 -w17 -e200 -r150 -m4000 -z200 -t
$threads --dual=yes $reads $reads
herro inference -t $threads -d 0,1,2,3 -m $model $input_reads
$output_file
```

### Genome assembly

#### Contig construction

Initial assemblies were generated using Hifiasm (v0.19.6-r595) and Verkko (v1.4.1) in trio mode with default parameters. Both assemblers produced comparable haploid genome sizes: 3.03 Gb for the maternal and 2.95 Gb for the paternal haplotype. Contigs from hifiasm assembly were gap-free, whereas the Verkko contained 723 kb and 784 kb of gaps (Ns) in the maternal and paternal haplotypes, respectively. Combining the assemblies from two assemblers yielded 20 telomere-to-telomere (T2T) contigs/scaffolds (with Ns) for the paternal haplotype (excluding chromosomes 15, 21, and 22) and 15 T2T contigs/scaffolds for the maternal haplotype (excluding chromosomes 5, 7, 9, 13, 14, 15, 16, and 22). An additional 325 contigs from Verkko assembly, totalling 15.21 Mb, were not assigned to either haplotype. Of these, 235 aligned to ribosomal DNA (rDNA) clusters, 82 to the Epstein–Barr virus (EBV) genome and the remaining 8 to high-repeat regions of non-acrocentric chromosomes were excluded. The mitochondrial genome was recovered by mapping contigs to the NC_012920.1 mitochondrial (MT) reference.

#### Scaffolding

Two distinct approaches were used for scaffolding contigs: Omni-C read–based scaffolding and a template-based method. For the maternal haplotype, binned Omni-C reads were aligned to the Hifiasm assembly and scaffolded using 3D-DNA, resulting in telomere-to-telomere (T2T) assemblies for chromosomes 5 and 7. For all remaining chromosomes in both haplotypes, a template-based approach was applied. In this method, maternal and paternal contigs from both assemblers were mapped separately to CHM13 chromosomes to determine contig order based on alignment coordinates (Additional methods, Figure 1). Importantly, CHM13 was used solely to infer contig order; neither CHM13 nor the contigs were modified during the scaffolding process. For each chromosome, contigs were merged when overlaps were detected; otherwise, they were concatenated with 1,000 bp of Ns. Scaffolding resulted in all remaining chromosomes T2T except maternal chr14, which contained telomere motifs on 3p.

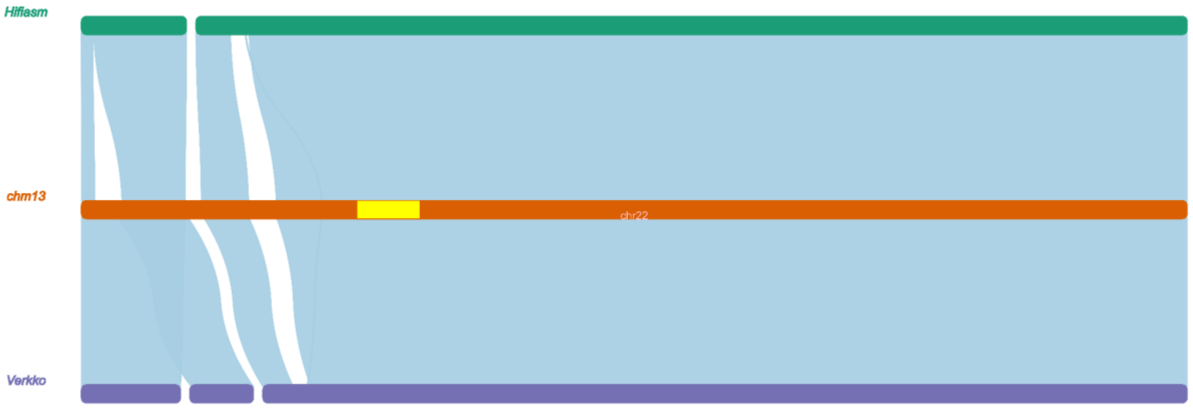


**Additional methods, Figure 1. Overview of template-based scaffolding.** Example shown for maternal chromosome 22. Contigs from Hifiasm (top) and Verkko (bottom) assemblies were independently mapped to the CHM13 chromosome (middle) to determine contig order for merging or concatenation. The yellow block indicates the centromeric region in CHM13.

**Telomere patching and extension**

Telomeric reads were extracted from binned long reads by identifying those containing at least 50 copies of the canonical human telomere repeat (TTAGGG) within 1 kb of either read end. To patch the maternal chromosome 14, telomeric ultra-long reads (>50 kb) were aligned to the first 200 kb of the assembled contig corresponding to chromosome 14. A consensus sequence was generated from the soft-clipped regions of aligned reads and used to patch the assembly. (Additional methods, Figure 2).

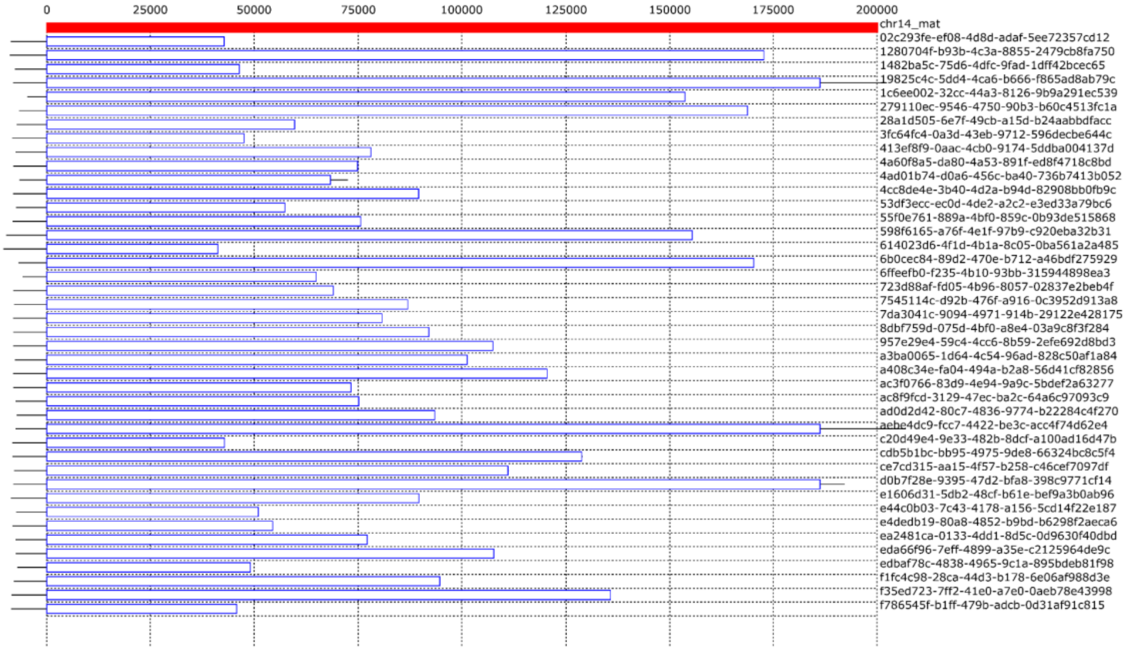


**Additional methods, Figure 2.** Telomere patching of maternal chromosome 14 using ultra-long reads.

After obtaining all 46 telomere-to-telomere (T2T) chromosomes, telomeric regions were further extended by aligning all binned telomeric long reads to the maternal and paternal assemblies, respectively, using minimap2. For each chromosome with at least three reads showing soft-clipping at the terminal bases, the soft-clipped regions were extracted. Consensus sequences were generated using SOPA and used to extend the corresponding telomeres. Read alignments were inspected before and after extension to confirm accuracy (Additional methods, Figure 3).

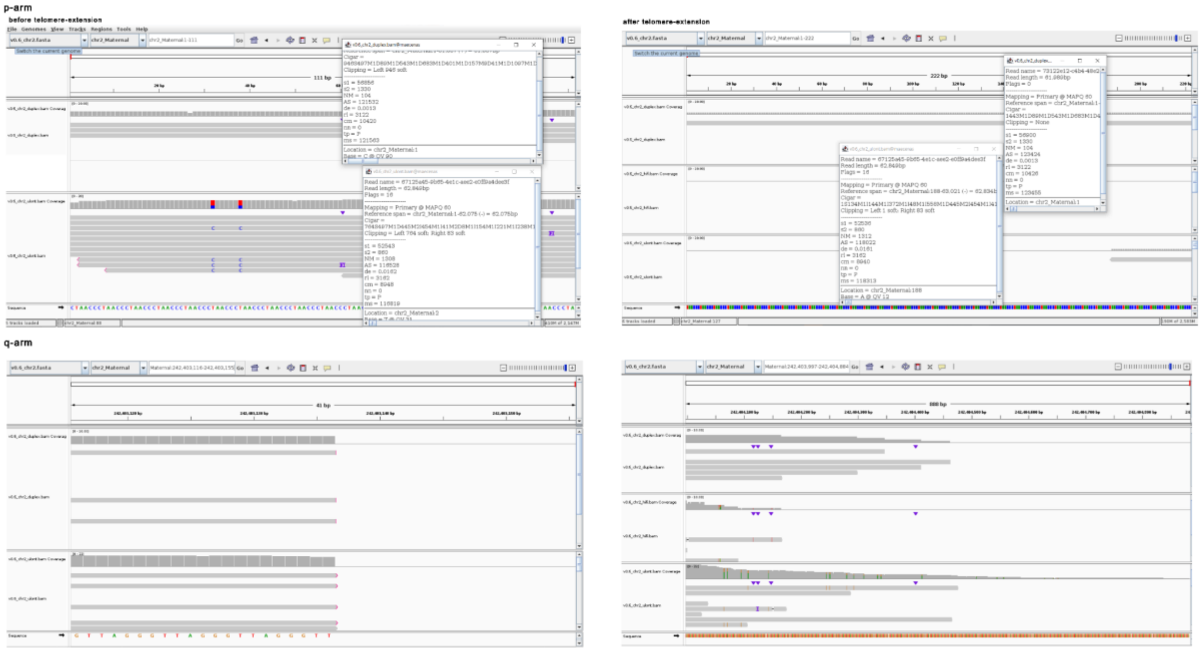


**Additional methods, Figure 3: Telomere extension workflow.** Example shown for maternal chromosome 2– Left: Telomeric reads aligned to chromosome ends showing soft-clipping at the 5′ (top) and 3′ (bottom) positions. Pop-up windows display alignment details. Right: Read alignments after telomere extension using consensus sequences derived from the soft-clipped regions.

#### Gap-filling

After the scaffolding step, the assemblies contained unresolved regions, represented by Ns—14 in the paternal haplotype and 39 in the maternal haplotype. To assess the nature of these gaps, long-read mappings were examined at the corresponding loci. Based on the mapping patterns, gaps were classified into two categories.

1) **Artificial gaps** were erroneously introduced during scaffolding and were identified by continuous, uninterrupted read coverage across the gap region after Ns were removed.
2) **Real gaps** were reflected by missing sequences and were supported by irregular read alignments, such as large insertions, large deletions, or consistent soft-clipping at the gap boundaries.

| | PAT | MAT |
|---|---|---|
| Artificial gaps | 2 | 2 |

| | | | |
|---|---|---|---|
| Real gaps | Patching | 1 | 33 |
| | Bigger Deletion | 6 | 3 |
| | TGS-GapCloser | 4 | 1 |
| | | 13 | 39 |

To resolve the real gaps, we employed two complementary approaches. First, long-read-based gap closure using TGS-GapCloser, which successfully closed five gaps. For the remaining unresolved gaps, we performed targeted reassemblies using binned reads. We generated reassemblies from duplex binned reads using LJA and hifiasm combined with RAFT, from HiFi binned reads using LJA and hifiasm, and from corrected ultra-long reads using Verkko. For each unresolved gap, we manually inspected the alignments of corrected and uncorrected ultra-long (UL) reads, duplex reads, as well as contigs from the previously mentioned reassemblies, using Integrative Genomics Viewer (IGV) to identify appropriate strategies for gap closure. In 10 cases, we identified large deletions supported by both reassemblies and long-read mappings, and we removed those regions from the assembly. In the remaining 35 cases, we identified suitable patches from the alignments and used them to successfully close the gaps.

#### rDNA

##### Reconstruction of rDNA regions in acrocentric chromosomes

rDNA regions on acrocentric chromosomes are a challenging part for de novo assembly. For I002C, rDNA segments were assembled during whole-genome assembly; however, due to the large copy number (CN), high repetitiveness, and sequence similarity, both Verkko and hifiasm over-compressed the rDNA arrays, resulting in underrepresentation of CN and variation. Therefore, we conducted a local reassembly of the rDNA regions to better reflect the dynamics of rDNA sequences. The rDNA local reassembly consists of three major steps: 1) estimating the overall and per-haplotype CN, 2) assigning rDNA-containing reads to acrocentric chromosomes through local assembly, and 3) identifying rDNA morphs for each chromosome, constructing consensus sequences, and patching morphs with estimated relative CN.

**Estimation of rDNA copy numbers for the whole genome and per haplotype**

We used two complementary methods to estimate the total rDNA CN in I002C. First, we applied a kmer-based method, following Nurk *et al*[2] for computational estimation. We counted 31-mers from I002C short-read data that matched the kmer profile of the 18S, 5.8S, and 28S regions in the canonical rDNA reference sequence (GenBank: KY962518.1), and normalized the counts by coverage. Peaks in the corresponding regions indicated that the total rDNA CN in I002C exceeds 400 copies (Additional methods, Figure 4). Second, we performed digital PCR (dPCR) to experimentally determine the rDNA CN. Using TBP1 and MRO as single-copy reference genes, we estimated an average of 209 rDNA copies per haplotype, corresponding to a total of 418 copies per diploid genome. After accounting for a potential off-target amplification on chromosome 12 in the I002C, a signature also observed in CHM13, we adjusted the estimated total acrocentric rDNA CN to 416 copies (Additional methods, Figure 5). The k-mer based and dPCR estimations of the total number of rDNA copies in I002C are consistent.

We then estimated the rDNA CN per haplotype using corrected binned ultra-long reads. By calculating the ratio of rDNA-aligned bases of each haplotype to the total rDNA-aligned bases, we estimated 259 copies for the paternal and 157 copies for the maternal haplotype.

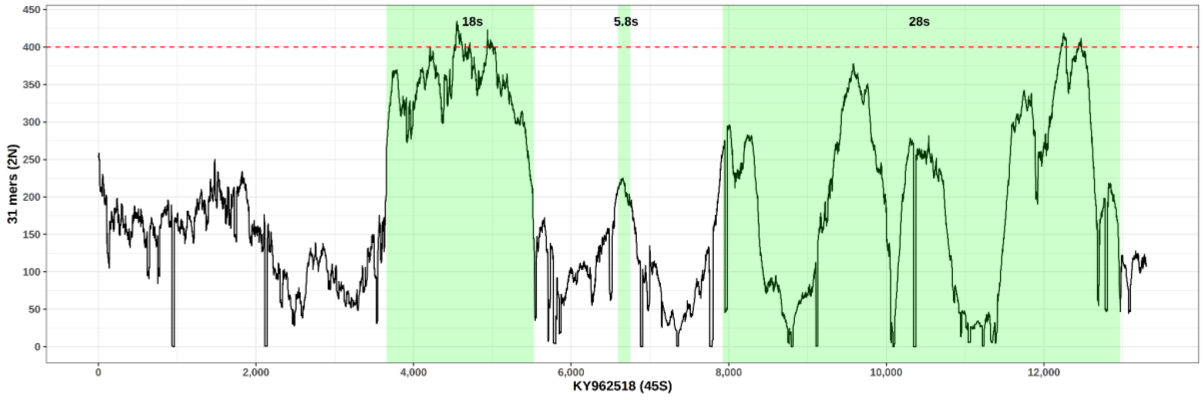


**Additional methods, Figure 4**: **kmer-based estimation of total I002C rDNA copy number**

**rDNA copy quantification by digital PCR**

Genomic DNA (gDNA) was extracted using the Gentra Puregene Kit (#158043, Qiagen, Germany) according to the manufacturer's instructions. The quantity and purity of the extracted gDNA were assessed using the Qubit dsDNA BR Assay Kit (#Q32853, ThermoFisher Scientific, USA) and the NanoDrop 2000 spectrophotometer (#ND-2000, ThermoFisher Scientific).

To fragment the tandem repeats of rDNA for digital PCR, 70 ng template DNA was digested with 1 µL of HaeIII restriction enzyme (#R0108S, NEB, USA) at 37 °C for 50 minutes, followed by heat inactivation as per the manufacturer's recommendations. A single digestion reaction was performed for all three targets, TBP1, MRO, and rDNA, to ensure consistent template processing. Digestion conditions were optimized to ensure complete and reproducible fragmentation.

Digital PCR reactions were prepared using the QIAcuity EG PCR Kit (#250113, Qiagen). For each 40 µL reaction, the mixture contained 2.5 ng of digested template gDNA, 0.4 µM of each forward and reverse primer, and 1× QIAcuity EvaGreen Master Mix, with nuclease-free water added to adjust the final volume. Master mixes for *TBP1*, *MRO*, and rDNA assays were prepared simultaneously, and the corresponding primers were added at the final step (Additional file 2: Table S32). The template input was optimized to achieve clear and distinct signal separation during digital PCR. To assess the specificity of the assays, qPCR was performed using the same reaction conditions. Melt curve analysis confirmed the presence of a single product for each primer pair, and product sizes were verified by agarose gel electrophoresis, confirming amplification of a single, correctly sized product.

Reactions were loaded into QIAcuity Nanoplate 26k 24-well plates (#250001, Qiagen) and run on the QIAcuity Digital PCR System following the manufacturer’s instructions. PCR cycling conditions included an annealing temperature of 49 °C and an elongation time of 15 seconds, with all other parameters set to default values.

Fluorescence imaging was performed using the green channel, with excitation at 463–503 nm, emission at 518–548 nm, an exposure time of 250 ms, and a gain setting of 3. Results were analyzed using QIAgility Software.

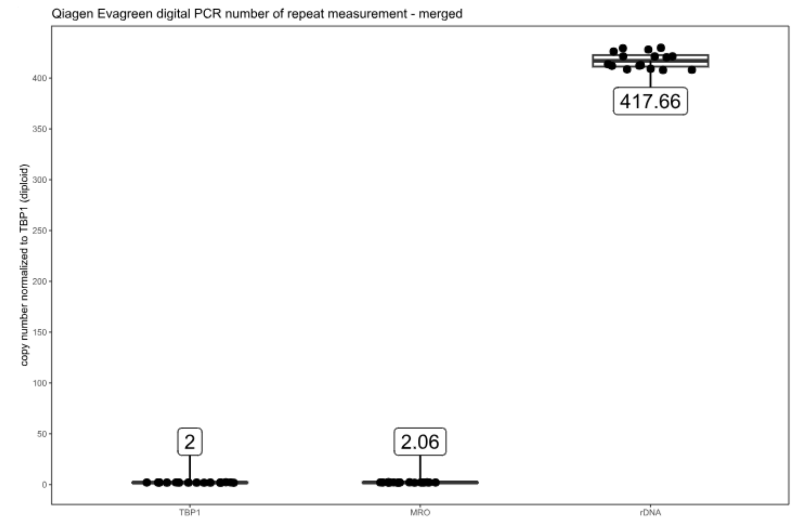


**Additional methods, Figure 5**. **Digital PCR-based estimation of rDNA copy number in diploid I002C using *TBP1* and *MRO* as reference genes.**

##### Local reassembly of rDNA arrays

We conducted the local rDNA reassembly of each haplotype by LJA. We found that the intermediate file mdbg.gfa generated by LJA provides a clear snapshot of rDNA array diversity, hence could serve as a basis for initial clustering of rDNA arrays. By empirically comparing mdbg.gfa visualizations using Bandage under different input combinations and LJA parameter settings, we identified clear clusterings for both paternal and maternal rDNA arrays (Additional methods, Figure 6). Because of the difference in complexity and diversity of paternal and maternal rDNA arrays, for the paternal assembly, we used corrected ultra-long reads spanning rDNA regions with default LJA parameters; for the maternal assembly, we used corrected ultra-long reads covering rDNA and 30 kbp flanking regions, further polished with hifiasm correction, and assembled with default LJA parameters. The presence of five clusters aligns with previous findings that rDNA arrays have greater inter-chromosome difference than intra-chromosome difference[2], suggesting they represent the five acrocentric chromosomes. We assigned the LJA-derived clusters to chromosomes by matching the variation patterns of distal junction (DJ) and proximal junctions (PJ) regions of the clusters to the I002C assembly.

Since LJA does not report read assignments to assembled contigs, we assigned corrected ultra-long reads to LJA clusters using a kmer-based approach with Merqury (k = 15). Each read was matched to the LJA-derived cluster with the fewest missing k-mers. To ensure confident assignments, we retained a read only if the top-matching cluster had at least 20% fewer missing k-mers than the second-best match. Given the important role of DJ and PJ anchoring reads in resolving rDNA arrays, we preserved all such reads and performed manual assignments by verifying small variants and SV patterns. In total, we obtained 5,271 and 3,532 corrected ultra-long reads assigned to the five paternal and five maternal acrocentric chromosomes, respectively. We estimated per-chromosome rDNA CN by integrating the proportion of reads supporting each cluster with the preliminary rDNA FISH experiment results. The final estimates showed strong concordance with our high-resolution FISH measurements (Spearman's rho = 0.97, $p = 2.4*10^{-6}$).

We use the clustered reads to identify the rDNA morphs for each chromosome. rDNA repeat units were grouped into morphs based on pairwise identity using UMAP. The pairwise identity between rDNA units was obtained from the "Number of residue matches" field of PAF-format output generated through minimap2 all-vs-all alignments. Due to the extreme complexity and high similarity of rDNA arrays, we prioritized the assembly to major morphs. For each chromosome, we kept only morphs supported by ≥ 10% of reads. Consensus sequences for major morphs were generated using high-quality supported reads (QV > 60) by SPOA. In cases where clusters exhibited minimal divergence (edit distance < 200) but had strong read support, we maintained these minor distinctions and designated them as sub-morphs under the corresponding major morph. Morphs near DJ/PJ boundaries were preserved regardless of coverage due to their critical role in array organization. CN per morph was estimated based on the number of supporting reads. The per-chromosome rDNA sequence was reassembled by concatenating the estimated CN of each morph and then integrated into the I002C final assembly.

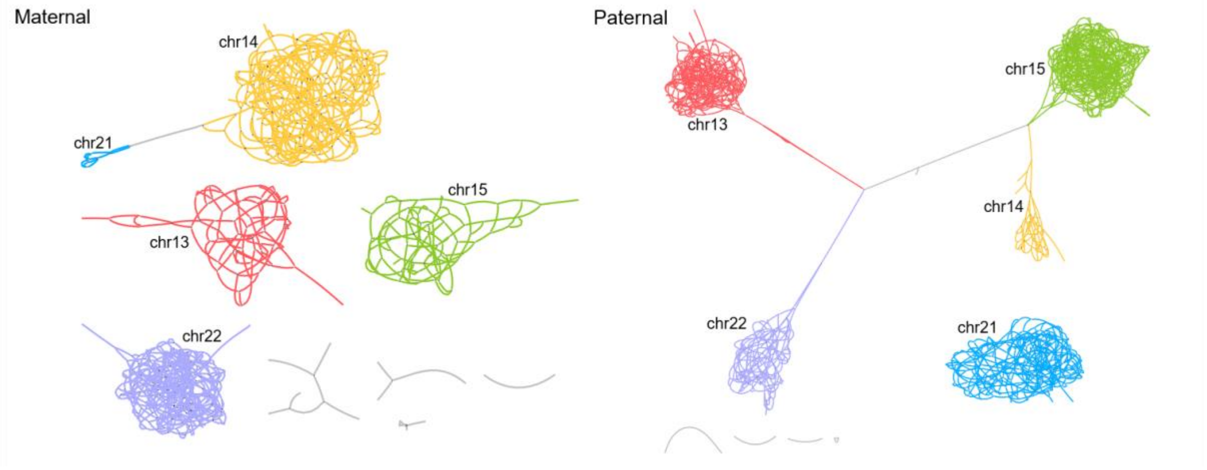


**Additional methods, Figure 6. Assembly graphs of local rDNA assembly by LJA.** Bandage visualization of the intermediate file (mdbg.gfa) generated by LJA for the maternal and paternal local rDNA assemblies, respectively.

#### Maternal chromosome 21 rDNA sequence

Maternal chromosome 21 rDNA was excluded from reassembly because its sequence assembled during the initial draft genome assembly (v0.1) was valid, comprising ten rDNA units that can be classified into three major morphs based on an edit distance threshold of 200. We evaluated and validated the completeness of the maternal chromosome 21 rDNA cluster sequence by two approaches. Firstly, we identified two overlapping ultra-long reads with DJ/PJ-anchoring sequences that spanned the entire initial maternal chromosome 21 rDNA assembly. Secondly, the rDNA CN of maternal chromosome 21 in the initial assembly was consistent with both coverage-based estimates and FISH results.

We also preserved the maternal chromosome 13 rDNA sequence from the initial assembly, as the reassembled sequence was nearly identical in both morph composition and total CN, despite FISH results suggesting a lower CN. The major morph dominated the assembly, and no ultra-long reads were observed to extend beyond the consecutive repeats of this morph, making it challenging to identify overlapping reads that could confirm the assembly continuity.

#### Estimating rDNA copy number per acrocentric chromosome from Fluorescence in situ Hybridization (FISH)

***Metaphase spread and Fluorescence in situ Hybridization (FISH)*** For the preparation of metaphase spreads, cells were seeded on 100 mm dishes and grown for 48 hrs to 60-80% confluency. KaryoMAX™ Colcemid™ (#15212012, Gibco) at 0.2 µg/mL was then added to arrest cells in metaphase, and cells were incubated for 2 hrs. Cells were harvested and incubated with hypotonic 0.075 M KCl afterwards for 7 minutes at 37 °C. A small amount of Carnoy's Fixative (3:1 Methanol: Glacial Acetic Acid) was then titrated to the cell suspension and further incubated at 37 °C for 3 mins. Afterwards, the cells were centrifuged at 200 *g* for 5 mins and washed a further two times with Carnoy's Fixative. Fixed cells were dropped onto slides using a humidified slide warmer and air-dried for 1 min.

Slides were dehydrated in subsequent 70%, 85%, and 100% ethanol washes, 2 minutes for each concentration. After air-drying, a mixture of fluorescently labeled rDNA probe (#RP11-450E20-gold, Empire Genomics, USA), Prader-Willi/Angelman (SNRPN) probe (#LPU005, CytoCell, UK), and Prenatal 13 and 21 Enumeration probe (#LPA003, CytoCell), were added per slide and covered with a 22x22 mm coverslip and sealed with rubber cement. Slides were then baked at 75 °C for 5 mins on a slide warmer and then incubated overnight in a humidified chamber at 37 °C for at least 16 hrs. Coverslips were then removed, and slides were washed with a wash buffer (0.4X SSC+0.3% IGEPAL® CA-630, Merck-Sigma Aldrich #I8896) at 73 °C for 2 mins, followed by a second wash with 2X SSC+0.1% IGEPAL® CA-630 at 37 °C for 2 mins. Slides were then incubated with a 1:2000 Hoechst33342 (#H3570, ThermoFisher) for 15 mins, washed with 1X PBS for 3 mins, air dried, and mounted with VECTASHIELD® PLUS Antifade Mounting Medium (#H-1900, Vector Labs). Images were acquired with a widefield Nikon Ti-E system equipped with Hamamatsu ORCA Flash 4.0 v2 camera and 60x CFI Plan Apo VC 1.40 NA oil immersion objective lens. The channels used were: channel 1: 535/50 nm bandpass excitation, 565 nm longpass dichroic and 610/75 nm bandpass emission filters; channel 2: 500/20 nm bandpass excitation, 515 nm longpass dichroic and 535/30 nm bandpass emission filters; channel 3: 350/50 nm bandpass excitation, 400 nm longpass dichroic and 460/50 nm bandpass emission filters. The images were deconvolved with the Huygens Professional software version 18.04.0p2 (Scientific Volume Imaging).

***Estimating rDNA copy number per acrocentric chromosome from FISH images***

Acrocentric chromosomes were reliably identified based on a combination of distinct fluorescence labeling and morphological features. The Prader-Willi/Angelman region probe (SNRPN) was used to label chromosome 15 with red and green fluorescence signals, while the Prenatal 13 and 21 Enumeration probe distinctly labeled chromosomes 13 and 21 with green and orange signals. Orange and red signals were visualized using channel 1, whereas green signals were detected via channel 2. The Rhodamine 6G (gold)-labeled rDNA probe was visible in both channel 1 and 2, and consistently localized to the p arm (Additional methods, Figure 7). Most acrocentric chromosomes could be unambiguously identified in each image. In rare instances where acrocentric chromosomes appeared folded or overlapped with another chromosome, such cases were excluded from the final quantification.

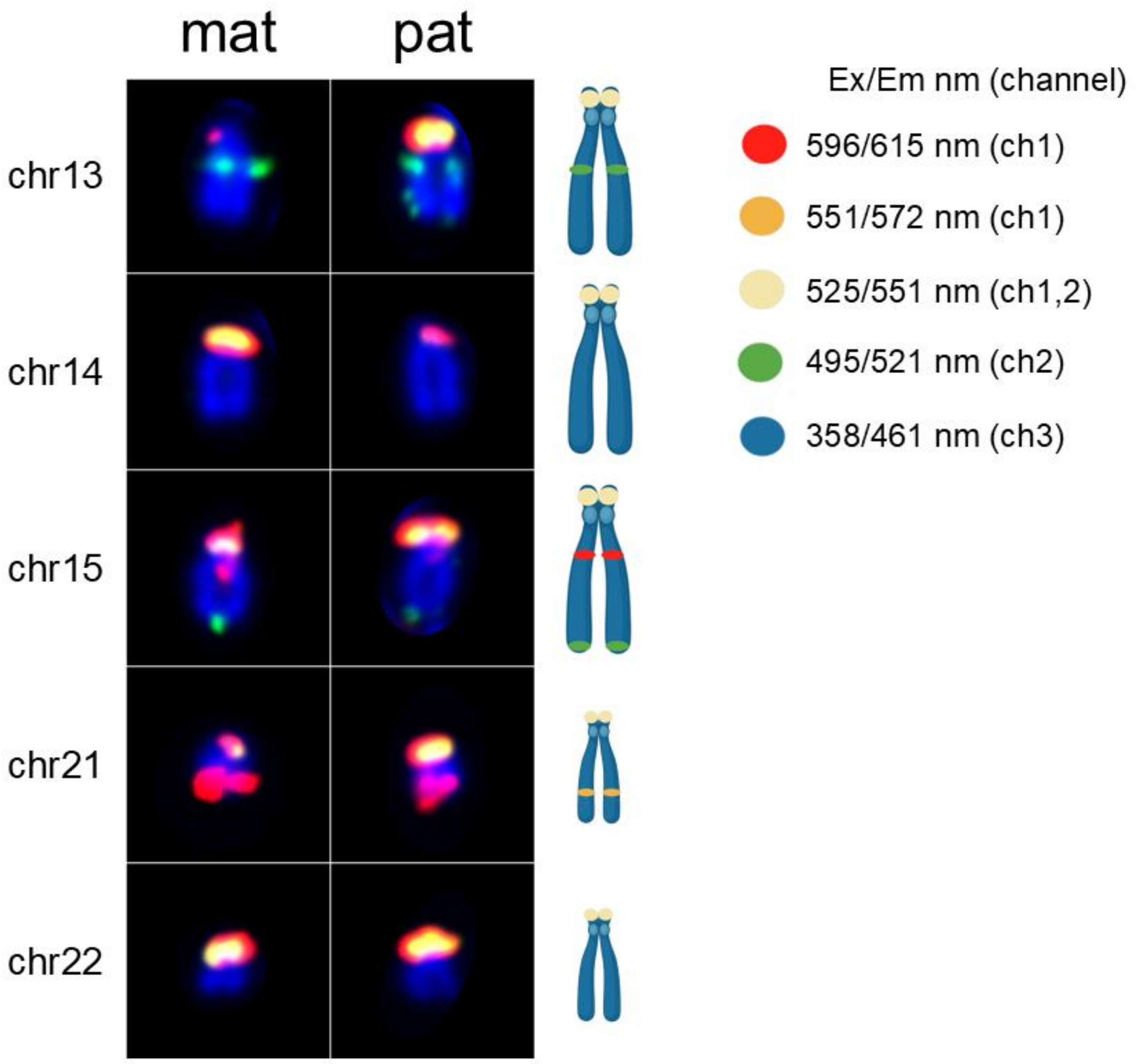


**Additional methods, Figure 7. Fluorescent In Situ Hybridization (FISH) karyogram of acrocentric chromosomes from an I002C metaphase spread, labeled using a mixture of fluorescently tagged probes.** Acrocentric chromosomes were identified based on a combination of distinct fluorescence labeling patterns and morphological features. Based on approximate chromosome size, the acrocentric chromosomes were grouped into two categories: larger (chromosomes 13, 14, and 15) and smaller (chromosomes 21 and 22). All acrocentric chromosomes display gold signals at the p-arm termini, corresponding to rDNA regions. Chromosome pairs were distinguished as follows: Chromosome 13 was identified by a green signal near the center of the chromosome and its large chromosome size. Chromosome 14 was identified as the only chromosome in the large group without additional fluorescence markers. Chromosome 15 was recognized by the presence of a red signal near the center, a green signal at the q-arm terminus, and its classification in the large chromosome group. Chromosome 21 was identified by a red signal near the q-arm terminus and its small size. Chromosome 22 was identified as the only chromosome in the small group without additional fluorescence markers. Each chromosome is present in two copies, consistent with a diploid metaphase spread. Chromosomes were counterstained with Hoechst33342. Images on black backgrounds show actual examples from the metaphase spread, while the schematic on the white background illustrates the conceptual design of probe combinations and chromosome sizes. Excitation and emission wavelengths, along with the respective fluorescence detection channels, are indicated in the accompanying legend.

To quantify rDNA fluorescence signals, images were analyzed using FIJI (ImageJ 1.54p)[104]. Elliptical regions of interest (ROIs) were manually drawn around each rDNA signal, and multiple elliptical ROIs were also drawn randomly at the termini of non-acrocentric chromosomes to represent background fluorescence. The mean background intensity was calculated across these regions after excluding the highest and lowest values to minimize outlier effects. For each rDNA ROI, the background-corrected integrated intensity was computed by subtracting the product of the mean background intensity and ROI area from the raw integrated signal intensity.

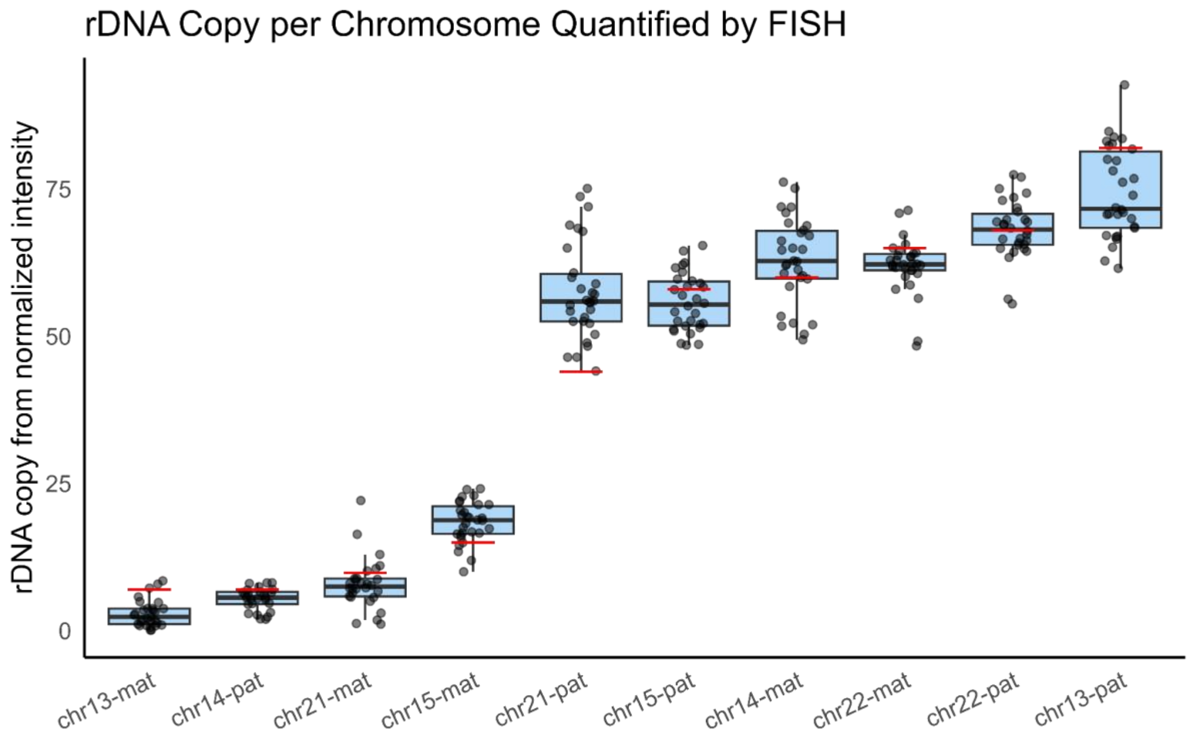


**Additional methods, Figure 8. rDNA copy number per chromosome, quantified from the main FISH experiments.** For each metaphase spread, rDNA fluorescence intensity was measured once per acrocentric chromosome. The intensities were normalized to the total rDNA signal across all chromosomes in the same spread and then scaled to absolute copy number using digital PCR measurements (Additional methods, Figure 5). Each dot represents a background-corrected rDNA measurement from a single metaphase spread. Red horizontal line segments indicate the number of rDNA copies patched into the I002C v0.7 reference genome.

The total background-corrected fluorescence intensity from all rDNA loci was then normalized to the total rDNA CN independently estimated by digital PCR. This enabled the derivation of an average unit rDNA fluorescence intensity, which was subsequently used to estimate rDNA CN for each acrocentric chromosome (Additional methods, Figure 8). Notably, background-corrected signals from the randomly drawn ROIs consistently corresponded to less than one rDNA copy in the final images (Additional methods, Figure 9), underscoring the low background noise in the images. Due to the consistently similar rDNA signal intensities between the paternal and maternal copies of chromosome 22 across all images, these homologs could not be reliably distinguished by FISH alone. Therefore, haplotype-specific CN estimates for chromosome 22 were assigned with the aid of binned coverage estimation detailed in the previous section.

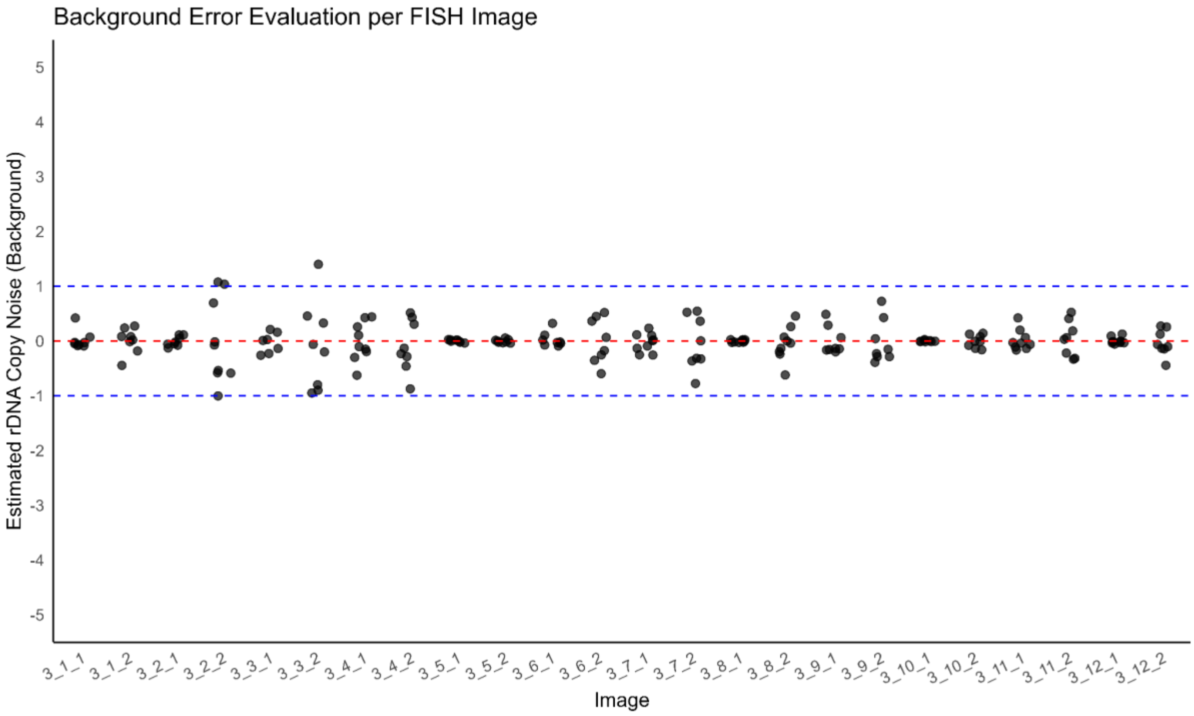


**Additional methods, Figure 9. Estimation of baseline noise and technical variability in rDNA copy number quantification**, based on measurements from randomly drawn regions of interest (ROIs) located at the ends of non-acrocentric chromosomes.

##### Genome polishing and version progression

Genome polishing was carried out in five iterative steps using a combinatorial approach. This included two rounds of automated polishing, one round of manual curation, one round of assembly-based patching, and a final step in which rDNA CN was corrected followed by an additional round of automated polishing. Automated polishing was performed genome-wide, whereas manual curation and assembly-based patching were restricted to error-prone loci identified by Merqury using hybrid k-mers generated from PacBio HiFi and MGI short reads (https://github.com/arangrhie/T2T-Polish/tree/master/merqury)

**Versions 0.1 and 0.2**

Initial assemblies produced by Hifiasm and Verkko were labelled as v0.1. Following scaffolding and telomere patching, the assembly containing all 46 T2T chromosomes was marked as v0.2.

**From v0.2 to v0.4**

Version v0.4 was generated after two rounds of automated polishing using binned long reads and unbinned short reads, following the T2T polishing pipeline (https://github.com/arangrhie/T2T-Polish/blob/master/doc/T2T_polishing_case_study.md).

**From v0.4 to v0.5**

Alignment and variant call files (SNVs and SVs) were generated as previously described. Error loci were inspected in IGV, and selected variants were used for manual polishing. All curated variants and decisions are provided (https://docs.google.com/spreadsheets/d/1Ij5S-RcrHaJY2hl8Zct_m0j7NLoAHckU/edit?usp=drive_link&ouid=108523434832954281275&rtpof=true&sd=true)

**From v0.5 to v0.6**

Remaining error-prone regions in v0.5 were evaluated against corrected ultra-long read assemblies (alt_assembly) generated with Verkko in trio mode. For each chromosome, 1 kb flanking sequences surrounding the target region were extracted and aligned to the corresponding chromosome in the alt_assembly using nucmer. The matched region from alt_assembly was then mapped back to the v0.5 assembly to assess whether it was mapped as a reciprocal best match at the same locus. For regions lacking a one-to-one reciprocal match, the flanking sequence was extended up to 10 kb. Regions lacking a unique reciprocal alignment were excluded from further consideration. For candidate regions showing one-to-one mapping, sequence quality in the alt_assembly was evaluated based on hybrid k-mer concordance and the presence of uniform read coverage without structural inconsistencies. Segments with higher support in the alt_assembly were selected for patching into the main assembly. Following replacement, long-read alignments were visually inspected to confirm correctness and integrity (Additional methods, Figure 10).

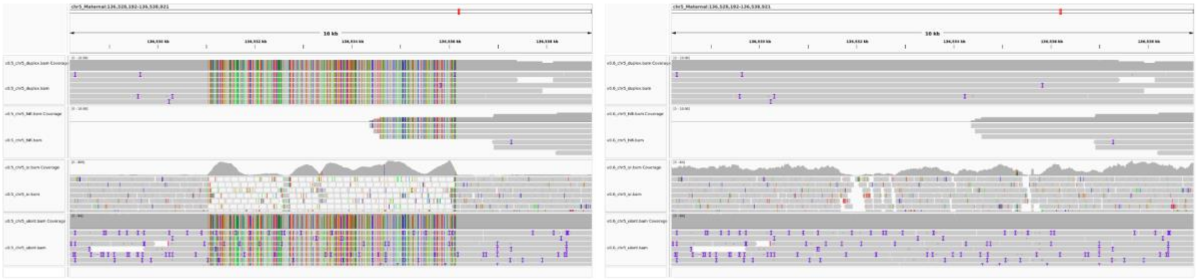

**Additional methods, Figure 10: Correction of assembly errors using alternate assembly derived patches.** An example is shown for a locus on maternal chromosome 5. Left: Read alignments at the error region in v0.5, showing inconsistent consensus between reads and assembly. Right: The same region post-patching in v0.6, demonstrating uniform read coverage and resolved consensus.

**From v0.6 to v0.7**

Based on genome-wide estimates of ribosomal DNA (rDNA) copy number, rDNA repeat units were appended to the short arms of each acrocentric chromosome across both haplotypes, following the approach described previously.

#### Methylation

Raw Nanopore R10.4.1 signals were basecalled using Dorado (v0.7.2, https://github.com/nanoporetech/dorado) with the model dna_r10.4.1_e8.2_400bps_sup@v4.1.0_5mCG_5hmCG@v2 under default parameters. This model estimates single-read methylation probabilities for cytosines in the CpG context. Reads were mapped to reference genomes with the Dorado *aligner* function. Per-read methylation data were aggregated across overlapping reads using modbam2bed (v0.9.1, https://github.com/epi2me-labs/modbam2bed) to compute per-site methylation ratios. At each locus, the methylation ratio was calculated as the number of methylated reads divided by the sum of methylated and unmethylated reads, considering only reads with explicit methylation (MM/ML) tags.

For PacBio Revio sequencing, HiFi reads with methylation information inferred from DNA polymerase kinetics were aligned to the reference genome using pbmm2 (v1.17.0, https://github.com/PacificBiosciences/pbmm2) in HiFi mode with default settings. Per-site methylation probabilities were extracted using pb-CpG-tools (v2.3.2, https://github.com/PacificBiosciences/pb-CpG-tools) with the pileup_calling_model.v1.tflite model and default parameters. The methylation ratios reported are derived from PacBio's internal algorithm, which correlates well with the ratio of methylated to total reads at a given locus.

For genome-wide and centromeric methylation analyses, Nanopore and PacBio reads were aligned to a combined I002C reference genome comprising the paternal and maternal haplotypes, the mitochondrial genome, and the EBV genome (NC_007605.1), given that I002C is a lymphoblastoid cell line immortalized by EBV. For genome-wide methylation analysis from both long reads sequencing platforms, loci covered by fewer than four reads containing MM/ML tags were excluded to reduce noise. For sliding window analysis, genomic windows were generated using the BEDTools (v2.30) *makewindows* function, with a step size equal to one-tenth of the respective window size. Average methylation levels within each window were calculated using the BEDTools *map* function. The Pearson correlation coefficient between Nanopore and PacBio methylation levels was computed in R (v4.3.2), both at common single CpG loci and across predefined genomic window sizes of 250/500/1000/1500/2000/5000/10,000/20,000/25,000/30,000/35,000/50,000/75,000/90,000/100,000/150,000 bp. Centromeric DNA methylation dip regions (CDRs), as defined in a previous study[31], were manually examined and annotated across all chromosomes.

For allelic methylation analyses, only haplotype-specific reads (excluding homozygous reads) were used. These reads were separately aligned to the GRCh38 + EBV hybrid reference genome to ensure consistent coordinate systems for both paternal and maternal haplotypes, enabling differential methylation analysis using methylKit (v1.24.0)[105]. Analyses focused on autosomal CpG motifs only, since sex chromosomes lack homologous pairs in this male donor. Differentially methylated loci (DMLs) were identified based on a minimum methylation difference of 97.5% between paternal and maternal alleles (with methylation ratios ranging from 0 to 100%) and a q-value cutoff of 0.01. To reduce noise, loci with fewer than five reads carrying methylation tags were excluded from analysis.

To discover novel uniformly differentially methylated regions (DMRs), the genome was binned into 200 bp sliding windows with a 100 bp step size. Within each window, directional consistency was assessed by calculating the proportion of hypermethylated loci among all DMLs. Windows with inconsistent directionality (i.e., 25–75% hypermethylated sites) were excluded. The remaining windows were classified as either hyper- or hypomethylated. They were merged separately into larger regions using the BEDTools *merge* function, considering only adjacent windows with the same methylation direction within a 1,000 bp window. This ensured that hyper- and hypomethylated regions were not combined into mixed-direction segments. Regions containing fewer than five DMLs were discarded. To further ensure directional consistency after merging, regions were re-evaluated, and those with 25–75% hypermethylated sites were excluded.

Identified DMRs were annotated to regulatory elements using the BEDTools *intersect* function with ENCODE candidate cis-regulatory elements (cCREs; downloaded from the UCSC Genome Browser, September 2025). To assess whether the observed overlap was greater than expected by chance, permutation testing was performed by randomly shuffling DMR coordinates across the genome with the BEDTools *shuffle* function while constraining placement to the same chromosome. Each DMR was counted once if overlapping any cCRE, regardless of whether multiple element types were intersected. This randomization procedure was repeated 1,000 times to generate a null distribution of overlap counts, from which the expected mean and empirical p-value were calculated (p-value defined as the proportion of permutations with an overlap count ≥ the observed value, with +1 correction). Identified DMRs were also compared with previously reported imprinting regions using the BEDTools *intersect*. For comparison, only regions reported in two or more independent studies were included. To identify potential imprinting genes, the DMRs were intersected with the core promoter and first exon regions of all transcripts. Core promoters were defined as 300 bp upstream and 100 bp downstream of experimentally validated transcription start sites (TSS), based on GENCODE v47 annotation in UCSC-style hg38 coordinates, with transcript strand taken into consideration. Monoallelic expression was defined as transcripts from the predominant allele comprising ≥92.5% of the total isoform expression, based on cDNA Nanopore sequencing analyzed with Bambu, with a minimum total isoform expression count of one. Concordant allele-specific methylation and transcription were defined as methylation of the core promoter and/or first exon of one allele, accompanied by expression from the other.

#### Gene annotation

*liftOver (conda version 377)*

```
nf-LO/main.nf --target I002C_mat.fasta --source GRCh38.p14.genome.fa --
outdir dir -profile local --aligner minimap2 --max-memory 800 --max_cpus
128
nf-LO/main.nf --target I002C_pat.fasta --source GRCh38.p14.genome.fa --
outdir dir -profile local --aligner minimap2 --max-memory 800 --max_cpus
128
```

```
liftOver -gff gencode.v48.annotation.gff3 liftover.chain liftover.mat.gff3
liftover.mat.unmap.gff3
liftOver -gff gencode.v48.annotation.gff3 liftover.chain liftover.pat.gff3
liftover.pat.unmap.gff3
```

*liftoff (version v1.6.3)*

```
liftoff -g gencode.v48.annotation.gff3 -o gv38ToMat.gff3 -u
gv38ToMat.unmap.gff3 -p 64 I002C_mat.fasta
liftoff -g gencode.v48.annotation.gff3 -o gv38ToPat.gff3 -u
gv38ToPat.unmap.gff3 -p 64 I002C_pat.fasta
```

*liftoff + liftOver combination*

```
python combine_liftoff_liftover.py gv38ToMat.gff3 liftover.mat.gff3
gencode.v48.annotation.gff3 mat.liftoff_liftover.gff3
python combine_liftoff_liftover.py gv38ToPat.gff3 liftover.pat.gff3
gencode.v48.annotation.gff3 pat.liftoff_liftover.gff3
```

```
combine_liftoff_liftover.py rules:
      case 1) liftoff and Liftover assign different strands:
              -  keep liftoff if coverage and sequence identity >= 0.85
              -  otherwise keep liftOver
      case 2) transcript names are inconsistent compared to the original
annotation:
              -  trust the original GFF
              -  if the annotation lifts on a completely different
                 chromosome than in liftoff/liftOver, discard it
      case 3) both sources agree on the chromosome and strand but different
coordinates:
              -  if they do not overlap -> keep both
              -  otherwise compare total exon length and keep the longer
                 one
```

*minimap2 alignment for each reference*

```
minimap2 -t 8 -ax splice GRCh38.p14.genome.fa I002C_cDNA_ONT.fastq.gz |
samtools view -bh -O BAM -o GRCh38_cDNA.bam
minimap2 index -@ 32 GRCh38_cDNA.bam

minimap2 -t 8 -ax splice I002C_mat.fasta I002C_cDNA_ONT.fastq.gz  |
samtools view -bh -O BAM -o I002C_mat_cDNA.bam
minimap2 index -@ 32 I002C_mat_cDNA.bam

minimap2 -t 8 -ax splice I002C_pat.fasta I002C_cDNA_ONT.fastq.gz  |
samtools view -bh -O BAM -o I002C_pat_cDNA.bam
minimap2 index -@ 32 I002C_pat_cDNA.bam
```

*Bambu processing (in R)*

```
input.bam <- “GRCh38_cDNA.bam”
gtf.file <- “gencode.v48.annotation.gff3”
```

```
ref_file <- "GRCh38.p14.genome.fa"
ndr_value <- 0.3
bambuAnnotations <- prepareAnnotations(gtf.file)
se <- bambu(reads = input.bam,
            annotations = bambuAnnotations,
                  genome = ref_file,
                  stranded = FALSE,
                  ncore = 4,
                  NDR = ndr_value)

writeBambuOutput(se, file = "grch38_bambu_output")
```

For haplotype-specific analysis:

```
# input.bam should be changed to I002C_mat_cDNA.bam, I002C_pat_cDNA.bam respectively
# gtf.file should be replaced by, mat.liftoff_liftover.gff3, pat.liftoff_liftover.gff3 respectively
# ref_file should be I002C_mat.fasta, I002C_pat.fasta respectively
# ndr_value: 0.3 and 1
# output_folder
```

For a more sensitive analysis:

ndr_value can be changed to 1
